# TimeTeller: a tool to probe the circadian clock as a multigene dynamical system

**DOI:** 10.1101/2023.03.14.532177

**Authors:** Denise Vlachou, Maria Veretennikova, Laura Usselmann, Vadim Vasilyev, Sascha Ott, Georg A. Bjarnason, Robert Dallmann, Francis Levi, David A. Rand

## Abstract

Recent studies have established that the circadian clock influences onset, progression and therapeutic outcomes in a number of diseases including cancer and heart diseases. Therefore, there is a need for tools to measure the functional state of the molecular circadian clock and its downstream targets in patients. Moreover, the clock is a multi-dimensional stochastic oscillator and there are few tools for analysing it as a system. In this paper we consider the methodology behind TimeTeller, a machine learning tool that analyses the clock as a system and aims to estimate circadian clock function from a single transcriptome by modelling the multi-dimensional state of the clock. We demonstrate its potential for clock systems assessment by applying it to mouse, baboon and human microarray and RNA-seq data and show how to visualise and quantify the global structure of the clock, quantitatively stratify individual transcriptomic samples by clock dysfunction and globally compare clocks across individuals, conditions and tissues thus highlighting its potential relevance for advancing circadian medicine.

## 1 Introduction

The mammalian cell-endogenous circadian clock temporally regulates tissue-specific gene expression in cells driving rhythmic daily variation in metabolic, endocrine, and behavioural functions. Indeed, around half of all mammalian genes are expressed with a circadian rhythm in at least one tissue [1, 2] and approximately 50% of all current drugs target the product of a circadian gene [3]. Moreover, recent studies demonstrated that the circadian clock influences therapeutic outcomes in a number of diseases including heart disease and cancer [4, 5, 6, 7, 8, 9, 10], and that disruption of the normal circadian rhythm and sleep (e.g., through shift work) is associated with higher risk of obesity, hypertension, diabetes, chronic heart disease, stroke and cancer [11, 12, 13, 14]. There is therefore a rapidly growing interest in developing circadian medicine tools that aid the incorporation of time in order to provide safer and more efficacious therapeutics. As a result a number of phase-estimation algorithms have been designed to estimate the molecular clock phase of the circadian clock, i.e., its “internal time”, from the measured levels of rhythmic gene expression [15, 16, 17, 18, 3, 19, 20, 21, 22, 23]. If the true timing of a sample is known, then divergence between the estimated timing and the true timing indicate the possible presence of clock dysfunction and, indeed, this internal phase has been proposed as a clinically actionable biomarker [24].

The core mammalian circadian clock involves a more than a dozen genes [25] and therefore the regulatory system is a high dimensional stochastic dynamical system. Since emergent systems properties such as oscillation, synchronisation, entrainment, phase-locking, robustness, flexibility and temperature compensation are critical for the functioning of the clock, tools that enable the analysis of the circadian clock’s systems properties are very much needed. However, probing the global behaviour of such a system is a highly non-trivial task and almost all analysis of clock data focuses on individual components and connections. On the other hand, a substantial amount of data is becoming available including whole transcritome time-series that should facilitate such systems analysis using mathematical modelling, statistics and machine learning.

Although we will provide a tool for estimating the phase of the clock, our main aim is to to delve deeper into the clock and its structure and quantify clock functionality independently in individual samples. We believe that it is difficult to do this, and perhaps impossible, without taking advantage of the clock’s structure as a stochastic dynamical system and the corollary that it therefore has a well-defined probabilistic structure describing the relationship between time and multidimensional gene state.

We want to quantify functionality in individual transcriptomic samples such as those from patients. To be useful in such a context this must be done in a way that allows comparison with the results on other similar transcriptomic samples. In particular, the results on a given test sample should be independent of those on other test samples and should not depend upon the particular test dataset being considered. This is crucial for reproducibility, comparability and interpretability. Even for timing estimation alone this does not seem possible with the phase-estimation algorithms mentioned above apart from TimeSignature [19] which requires two samples. However, the key point differentiating TimeTeller from TimeSignature and the other algorithms is that, apart from identifying timing deviations, these do not provide any other assessments of functionality or other quality controls on the individual timing assessments. This is essentially also true for ZeitZieger [16] but with the caveat that it, like TimeTeller, uses a likelihood curve that it might be considered could be used in a similar way to TimeTeller’s to assess functionality. However, although differences in ZeitZieger’s likelihood between WT/control and perturbed clocks in controlled lab situations has been discussed [16], it has not been proposed or statistically analysed as a measure of dysfunction and, moreover, has not been used as such when ZeitZieger has been employed to analyse timing variation in populations [26, 24, 27]. Moreover, analysis by ZeitZeiger of new data as described in [16] involves renormalizing and batch-correcting this data with the training data and then retraining, resulting in a different predictive model every time and therefore sacrificing the reproducibility, comparability and interpretability discussed above.

It is difficult to compare timing accuracy of these algorithms with TimeTeller as they have been used on different datasets collected under different conditions with different levels of genetic heterogeneity, but we show below that TimeTeller compares well in that its typical median timing error even on genetically heterogeneous human and baboon control data is around one hour and often significantly less. Finally, we note that any claim for reproducibility, comparability and interpretability for an algorithm must involve testing on external samples that are independent of the training data and, while this has previously been relatively rare, we provide many examples of this for TimeTeller. Regarding such external samples, one novel development we make is to develop a test of timing precision of our algorithm on two large cancer datasets for which we do not have time labelling. This shows that even though these data sets contain a large amount of clock dysfunction TimeTeller can nevertheless extract from the data the timing of samples with very good accuracy.

Statistical theory has clear messages about how to link the probability structure of the dynamical system to functionality that facilitates such an exploration. One such message is that the gene-gene correlations in the expression of the clock genes and the way this changes as a function of time will have a definite structure set by the dynamics, and this contains very important information about functionality. Indeed, one significant attempt [28] at a systemic approach to define molecular clock disruption has used pair-wise correlations between clock genes across large transcriptomic datasets. This showed greater dysfunction at the dataset level in solid tumours compared to healthy tissue. However, this approach compared datasets as a whole and, as the authors pointed out, it did not lend itself to assessing clock function in single samples.

Our aim in this paper therefore is to develop a tool that (i) provides a multidimensional picture of the clock’s dynamics and structure that integrates the behaviour of multiple genes, (ii) provides a quantitative analysis at the systems level of clock data, (iii) enables a quantitative comparison of different clocks and (iv) enables a quantitative assessment of clock dysfunction both in the core clock and in downstream target genes. We propose a tool that can determine the presence of a dysfunction causing perturbation from just one sample and that can stratify individuals on the basis of clock functionality, and, thus, might be useful to develop as a clinically actionable biomarker. For example, we show that such a stratification can enable the identification of clock and other genes that are differentially expressed between samples that have better and worse clocks. Finally, we consider new methods for comparing clocks across different individuals, tissues and conditions, identifying a “molecular chronotype” associated with these, and uncovering the effect of clock perturbations on downstream genes.

The above probability structure of the dynamical system is primarily described by the joint probability distribution *P*(*t, g*) of the external time *t* and the expression state *g* of the core clock genes or some representative subset of them. TimeTeller requires time-series data for training, but once trained, it can be used to analyse single samples independently. The training data will come from a clock with high functionality and test data will be compared to this. Timeteller estimates key aspects of *P*(*t, g*) for this training clock and uses these to address the problem of measuring how well the clock gene state *g* represents the external time *t* in test samples. Using this approach we get a much better idea of the nature of the dysfunction being observed and, in this way, we can identify and measure three types of clock dysfunction. It is a definite advantage if statistical methods used to analyse clock data can be related to this underlying structure in a transparent and reasoned way so that one can understand in what sense the methods estimate the important underlying probability distributions. Current methods for phase estimation do not have this property. In particular, our methods provide much more information on the reliability of our quantification.

The broad range of transcriptomes from microarray and RNA-seq data that we use throughout the paper is detailed in SI Sect. S1, as are the methods used to prepare the data for use with TimeTeller. We use different microarray or RNA-seq training datasets for mouse, baboon and human tissues. It is important to stress here that with the currently available data we will have to make and justify some assumptions on the cross-validity of data from different tissues in order to combine the data. For example, in order to estimate the probability model for a particular tissue we would ideally like to use training data that is only from that tissue. In particular, this is not possible for the mouse and baboon datasets as adequate amounts are not currently available and we therefore have to pool data from several tissues. To do this we choose an appropriate rhythmic gene panel with good cross-tissue synchronicity and, after validation of this, use normalisation to overcome tissue differences in the way explained below and in the SI Sect. S6. For our human data we pool across individuals rather than tissues. Another potential limitation comes from the fact that our current RNA-seq training data is only available at a few training timepoints around the day. Nevertheless, even with these handicaps we obtain very informative results and provide plenty of evidence that the approaches adopted work well. As more data becomes available this situation can only improve.

## 2 Results

### 2.1 Training with genetically homogeneous and heterogeneous data

The data that is used to prepare TimeTeller’s probability model is referred to as the *training data*. The data that is then analysed using this probability model is called the *test data*. In this paper we use four different training datasets and more details about these are in SI Sect. S1. For mouse training data we will use the microarray and RNA-seq datasets of Zhang *et al*. [1] which quantified the transcriptomes of 12 mouse tissues every 2 hours over 48 hours with microarrays, and every 6 hours using RNA-seq. We use 8 of these tissues excluding 3 brain tissues and white fat as the noise to amplitude ratios are higher for them. TimeTeller will also be trained on a human dataset, namely that of Bjarnason *et al*. ([29], Sect. S1.4) which used microarrays to analyse punch biopsies of oral mucosa taken every four hours over 24 hours from 10 healthy volunteers (five males). This data will be particularly interesting since, unlike the mouse training datasets, it is from genetically heterogeneous individuals and TimeTeller can be used to investigate this. Finally, we will utilise training data taken from the baboon dataset of Mure *et al*. [2] which includes 33 tissues from one animal every 2 hours for 24 hours. This is doubly challenging in that the baboons are genetically heterogeneous and we find that the multiple tissues considered have disparate expression and timing. In addition to the analysis of real data we have also used simulated data obtained by developing a stochastic version of a relatively detailed published model of the mammalian circadian clock [30] for design, testing and evaluation of the algorithm (Sects. S11 and S12).

#### 2.1.1 Choice of a clock representative gene panel

For a given training dataset we firstly choose the panel of *G* rhythmic genes that TimeTeller will use. This is called the *rhythmic expression profile* (REP). For a given transcriptomics sample the expression levels *g*_*k*_, *k* = 1, …, *G*, of these genes are collected into a vector *g* = (*g*_1_, …, *g*_*G*_) which we will call the *rhythmic expression vector* (REV). The user is free to to choose the genes in the REP and may have a particular reason to include or leave out a particular gene. However, in this study we firstly carry out an analysis of both the rhythmicity and synchronicity across tissues or individuals in our datasets to guide our choice. This analysis which is detailed in Sect. S8 is important to ensure the choice of a panel of genes with good circadian rhythmicity combined with minimal variation across the relevant tissues or individuals. We use this analysis to guide the choice of our REP from the highly ranked genes usually including a small number that are not considered to be core clock genes and then checking that any results do not depend on the choice of these. It is also necessary to try and ensure that the choice of the REP and rhythmic properties of the genes included provide a faithful representation of the clock state even though it does not contain all the core clock genes. Our approach to this is explained in Sect. S5.

#### 2.1.2 Timecourse and intergene normalisation

When combining training data from multiple tissues, for each gene in the REP we study the variation across the tissues in that gene’s expression time-series. This analysis (Sect. S6.1) shows that for RNA-seq data this variation is significantly greater than that found, for example, in the Affymetrix MoGene 1.0 ST and GeneChip Human Genome U133 Plus 2.0 microarray platforms that we have analysed. Therefore, for the RNA-seq training data, it is usually necessary to carry out a further normalisation to align the timecourses of the tissues.

Each of our training data sets is organised into time series for each gene in the REP with times *t*_*k*_, *k* = 1, …, *K*, that are usually independent of the particular gene. We can normalise the data by replacing each of these time series by a normalised version which has mean expression zero and standard deviation 1 (see SI Sect. S6.1). We call this approach *timecourse normalisation*. Following timecourse normalisation of a training dataset, if we wish to test an independent test sample REV from a given tissue and gene we will have to normalise the REV using the offsets and scalings that were used in the timecourse normalisation of the training data for this tissue and gene. Such normalisation of test data is called *timecourse-matched*.

There is, however, a cost in using timecourse and timecourse-matched normalisation because the test data has to be normalised using the adjustments calculated for the training data. This means that one can only use test data for tissues where we have a training time-series. Moreover, when using timecourse-matched normalisation on test data it is crucial that the training data are produced by the same transcriptomics platform (e.g., RNA-seq).

Intergene normalisation avoids this. When timecourse normalisation is unnecessary or is impossible because we do not have a training data set for the test data tissue, the data is normalised using *intergene normalisation* where if *g* = (*g*_*i*_) is a REV the normalised levels are given by *ĝ*_*i*_ = (*g*_*i*_ *−μ*)*/σ* where *μ* and *σ*^2^ are the mean and variance of the entries *g*_*i*_. It is also possible to usefully combine timecourse and intergene normalisation (see Table 1). Though use of timecourse normalisation typically improves timing performance, we will show that intergene normalisation can also be remarkably effective (e.g., see Table 1).

**Table 1:** Mean and median absolute timing errors for the training datasets. Column 1 shows the normalisation used. Columns 2 and 3 show respectively the mean and median absolute timing error. Column 5 and 6 show the mean and median absolute timing error after a correction is made using the timing displacement for the tissues or individuals as relevant. **A-E**. The timing errors for the three training datasets when a leave-one-out cross-validation approach was used. For the Zhang *et al*. microarray data we compare using all the data (2h resolution) to only a subset giving 6h resolution. **F**. Timing results for Zhang *et al*. RNA-seq test data when Zhang *et al*. microarray is used as training data. **G**. As **F**. but with datasets swapped.

| A. Zhang *et al.* 2014 Microarray, mouse |  |  |  |  |
| --- | --- | --- | --- | --- |
| normalisation | mean | median | corr. mean | corr. median |
| intergene | 1.39h | 0.93h | 1.35h | 0.94h |
| timecourse (2h) | 0.89h | 0.70h | 0.77h | 0.52h |
| timecourse (6h) | 0.78h | 0.63h | 0.61h | 0.53h |
| both | 0.90h | 0.60h | 0.84h | 0.59h |
| B. Zhang *et al.* 2014 RNA-seq mouse |  |  |  |  |
| intergene | 1.60h | 0.80h | 1.51h | 0.82h |
| timecourse | 0.68h | 0.46h | 0.58h | 0.59h |
| both | 0.64h | 0.27h | 0.66h | 0.56h |
| C. Bjarnason *et al.* human |  |  |  |  |
| intergene | 1.52h | 0.73h | 0.95h | 0.75h |
| timecourse | 1.14h | 0.86h | 0.70h | 0.62h |
| both | 1.06h | 0.48h | 0.67h | 0.48h |
| D. Mure *et al.* trained on central 18 tissues, baboon |  |  |  |  |
| intergene | 2.43h | 1.87h | 2.27h | 1.51h |
| timecourse | 1.49h | 1.11h | 1.21h | 0.90h |
| both | 1.49h | 0.90h | 1.23h | 0.86h |
| E. Mure *et al.* trained on all 33 tissues, baboon |  |  |  |  |
| intergene | 2.53h | 1.84h | 2.36h | 1.65h |
| timecourse | 1.53h | 1.13h | 1.24h | 0.88h |
| both | 1.53h | 0.94h | 1.27h | 0.92h |
| F. Test: Zhang *et al.* RNA-seq. Training: Zhang *et al.* microarray |  |  |  |  |
| timecourse | 0.83h | 0.51h | – | – |
| G. Test: Zhang *et al.* microarray. Training: Zhang *et al.* RNA-seq |  |  |  |  |
| timecourse | 0.86h | 0.67h | – | – |

We can also apply such timecourse normalisation to test data when this contains a time series; as many experimental model datasets do. However, this removes some amplitude information from the test data. Although some cross-tissue relative amplitude information is removed by timecourse normalisation, when test data is analysed using timecourse-matched normalisation any change in amplitude in the test data compared to the training data is maintained. This is not the case if test data is timecourse normalised.

Similar considerations to the above apply when combining data across individuals instead of tissues as we do with the Bjarnason *et al*. human data. Timecourse normalisation can also be very useful when analysing microarray data and we have found it necessary when the training and test data come from different microarray platforms (e.g., as in Fig. S19).

Table S5 summarises the normalisations that were used for all the analyses shown in the Figs. 1-5.

**Figure 1:**
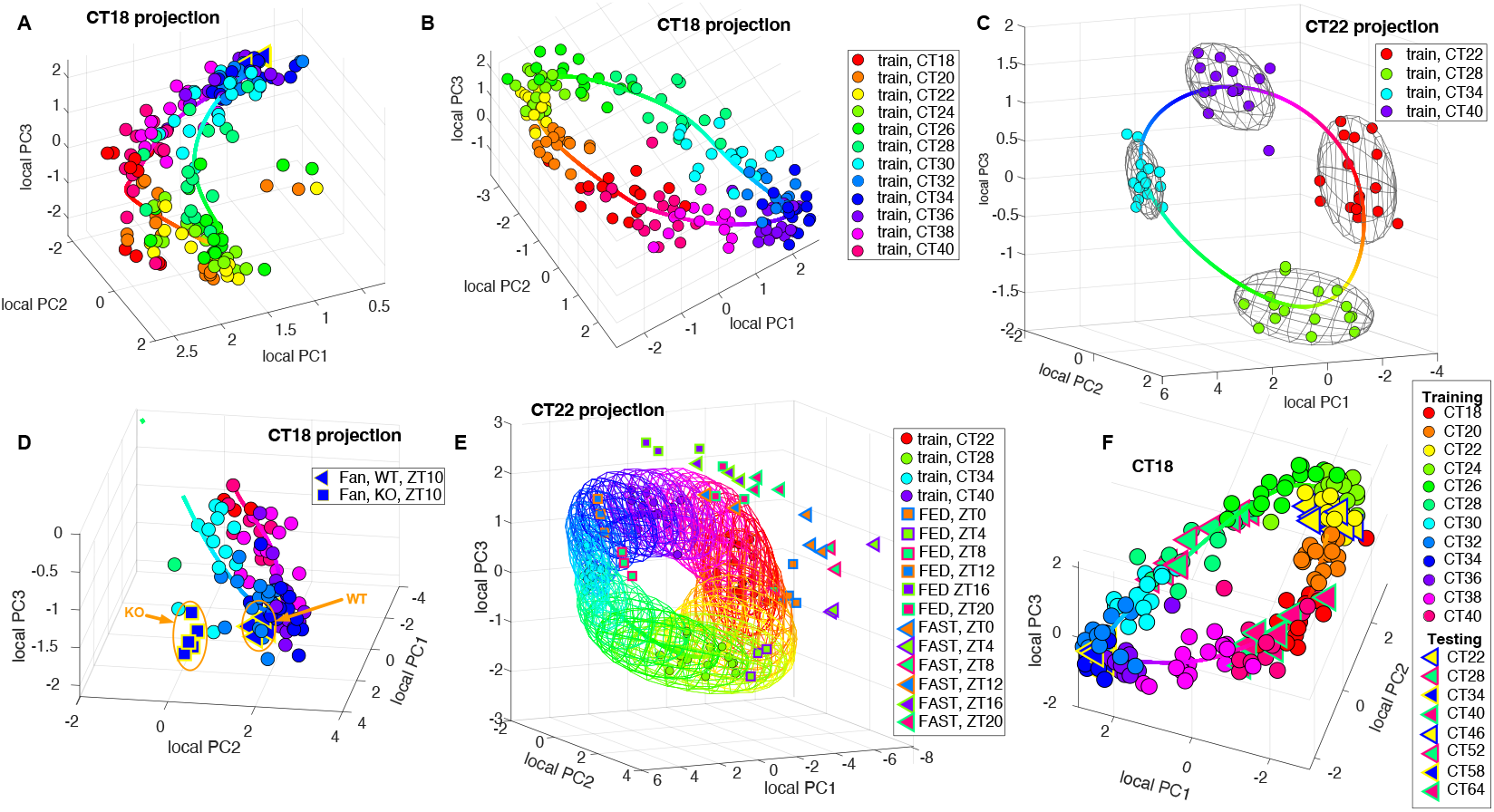
The color of datapoints etc (when not black) corresponds to the time when the data was sampled. This coloring is done in a consistent way across all figures. **A-F**. Using local PCA projection to visualise data. The identity of each data point can be read off from the legends to the right of each example. Only one projection for each example is shown but the differently timed projections have a similar quality. Examples showing all of the projections are in the SI (e.g., Figs. S9 S13). **A**. The projection using the CT18 local PCs of the Zhang *et al*. microarray data with intergene normalisation only. **B**. The CT18 local PCs of the Zhang *et al*. microarray data but using timecourse normalisation. **C**. The projection using the CT22 local PCs of the Zhang RNA-seq data with timecourse normalisation. The ellipsoids shown are of the form (*x − μ*)^*T*^ Σ^−1^(*x − μ*) = *ε* where *μ* is the mean of the estimated *P*(*g*|*t*) and Σ is its estimated covariance with *ε* chosen so that the ellipsoid should contain 97.3% of the training data (i.e. 3 standard deviations). This enables visualisation of the variation and covariation in the data. **D**. A detail from a projection of the the Fang *et al*. test data together with the Zhang *et al*. training data as in A showing coherence of the Fang *et al*. WT data and the gap between this and the KO data. **E**. The Kinouchi *et al*. RNA-seq skeletal muscle test data for FED and FAST mice plotted against the Zhang *et al*. RNA-seq training data with ellipsoids as in **C** added for a larger number of times around the day and colored according to the time. The divergence of the FAST data from the training data is clear. **F**. This shows a visualisation of both the test and training data when testing Zhang *et al*. RNA-seq data using Zhang *et al*. microarray data for training. Timecourse normalisation is used.

#### 2.1.3 Estimating the training clock’s statistical structure

We mentioned above that the joint distribution *P*(*t, g*) of time *t* and clock gene state *g* characterises the statistical structure of a clock. For us this distribution is always associated with the training clock while test data is analysed using this training clock probability distribution. In fact, rather than *P*(*t, g*) we will mainly be interested in two conditional distributions associated with it.

As the time *t* proceeds across the day the expression levels of the clock genes *g*_*i*_ = *g*_*i*_(*t*) oscillate in a coherent way and, since the clock is stochastic, this can be best described by the related probability distributions *P*(*g*|*t*) and *P*(*t*|*g*). If *g* denotes the REV state of the clock *g* = (*g*_*i*_), these distributions tell us respectively the distribution of *g* when the time is *t* and the probability distribution of times *t* that are found when the gene expression state of the clock is *g*. The distribution *P*(*t*|*g*) is a critical quantity because the cell has to use some function of the state of the gene products as a surrogate for *t* and the variance of *P*(*t*|*g*) tells us how well you can tell the time by just seeing the clock gene state. Moreover, *P*(*g*|*t*) is important because if *g* comes from the training clock at time *t* then we expect *P*(*g*|*t*) to be high so if *g* comes from a test clock and *P*(*g*|*t*) is low then we see that (*t, g*) differs from typical states in the training clock.

TimeTeller aims to use the training data to estimate *P*(*g*|*t*) for all times *t* across the day as explained in Sects. S2-S4. Moreover, *P*(*g*|*t*) and *P*(*t*|*g*) are related by Bayes’ law and in the case of clocks this boils down to the fact (since times *t* are equally probable) that, as functions of time *t, P* (*t*|*g*) is approximately proportional to *P*(*g*|*t*). Therefore, for any clock gene state *g* from training or test data we can use knowledge of *P*(*g*|*t*) to determine the temporal shape of *P*(*t*|*g*). Furthermore, as we explain in Sect. S4, variance of *P*(*t*|*g*) depends crucially on the covariance structure of the clock genes, i.e. the covariance matrix of *P*(*g*|*t*). Our tool is constructed to use this understanding. Finally, we note that the stochastic dynamics of the system around its periodic attractor modified by measurement noise sets the nontrivial structure of this covariance matrix. From theoretical considerations [31], if the measurement noise is not too large, we can expect that the covariance matrix has rapidly decaying eigenvalues, an observation that will justify our dimension reduction from *G* to less dimensions that is discussed below (also see Fig. S2).

### 2.2 Multidimensional visualisation provides important information about phenotype

The training and test data REVs consist of *G*-dimensional vectors of expression levels. When constructing the probability model, the TimeTeller algorithm projects this into fewer dimensions using a local version of principal component analysis using the the singular value decomposition (SVD) algorithm (Sect. S7, Figs. S1 & S2). This gives a different projection for each time in the dataset and the algorithm extends this to all times around the day. If for the *G*-dimensional data the distributions *P*(*g*|*t*) are approximately multivariate normal (MVN) then the corresponding distributions of the projected data optimise the capture of the dominant gene-gene correlations after projection (SI Sect. S2). We find that for our datasets *d* = 3 is sufficient for this (e.g., see Fig. S2) and the resulting 3-dimensional model of the clock provides a very informative visualisation.

Figs. 1A & B show such a visualisation for the mouse multi-organ microarray training data from Zhang *et al*. [1] when intergene (Fig. 1A) and timecourse normalisation (Fig. 1B) are applied, while Fig. 1C shows the timecourse-normalised RNA-seq training data from the same study. TimeTeller actually produces such a local projection visualisation for each time in the training dataset as shown in Fig. S9 but normally inspection of just one of these is adequate and we only show one in Fig. 1. With each such visualisation we also show the curve given by the means of the estimated distributions *P*(*g*|*t*) as *t* varies over the day. Also in such plots we often provide for a sample of times *t* an ellipsoid showing the covariance structure of the estimated distribution *P*(*g*|*t*) (see caption of Fig. 1). We color the training data points and mean curve by time with a color coding as given in the legend of Fig. 1 using the sample time for the data points. The same color coding is used throughout the paper.

Fig. 1D plots microarray test data from Fang *et al*. [32] comparing it with the Zhang *et al*. microarray training data in Fig. 1A. This test data compares liver samples of *Nr1d1* (*Rev-erbα*) KO and WT mice entrained to light-dark (LD)12:12 cycles. The gene *Nr1d1* is a core clock gene of the mammalian circadian clock important in one of the interlocked feedback loops of the clock and a key link to metabolism [33]. Knocking it out leaves a functional but perturbed clock when compared to WT mice [32]. Since *Nr1d1* is a member of the default REP it would not be surprising that TimeTeller could distinguish *Nr1d1* KO mice from WT mice, and indeed this is the case. Therefore, for this validation, we exclude *Nr1d1* from the REP genes. The visualisation shows that while the WT data appears to fit well with the training data, the KO data has a consistent substantial difference. The visualisation is able to detect this apparent difference in each single sample and shows the coherence across the four KO samples. It suggests that the *Nr1d1* KO mice have a significantly perturbed clock when compared to WT mice (Fig. 1D) but that it is still somewhat functional as it gives approximately correct timing and the level of sample variation between WT and KO is similar. We investigate this further below.

The next test data we visualise in this figure is from Kinouchi *et al*. [34]. This contains samples analysed by RNA-seq from mouse skeletal muscle taken around the clock in the presence of a light/dark cycle, i.e., entrained conditions, [34], and compares mice that had been fed *ad libitum* (FED) with mice that had been starved for exactly 24 hrs prior to point of sampling (FAST). In Fig. 1E we see that, while the FED samples align nicely with the RNA-seq training data, the FAST samples are substantially perturbed. One the other hand, the FAST samples show consistency in that for a given sample time they tend to cluster together. A similar viualisation for the liver samples from[34] is given in SI Sect. S9.4. It should be noted that the test samples from Kinouchi *et al*. have been collected in LD whereas the training dataset was collected in constant conditions. Interestingly, there is little difference between FED (control) and training dataset mice, which might be due to the fact that the free-running period of these WT C57Bl/6 mice is very close to 24 hours.

Other examples demonstrating the utility of such visualisation are discussed below.

### 2.3 Analysis of single test samples

TimeTeller’s estimate of *P*(*t*|*g*) from the training data is used to analyse test data. For a normalised test data REV *g* our estimate *L*_*g*_(*t*) of *P*(*t*|*g*), which we regard as a function of *t*, is referred to as the *likelihood curve* (LC) for the corresponding transcriptomics sample. The quantities for functionality assessment are associated with this LC. For example, our estimate of the time when the independent sample with this REV *g* was taken is the time *T* at which the estimated likelihood function *L*_*g*_(*t*) *≈ P* (*t*|*g*) is maximal i.e., the maximum likelihood estimate. Given *T*, we define the *likelihood ratio function* (LRF) as *R*_*g*_(*t*) = *L*_*g*_(*t*)*/L*_*g*_(*T*) i.e., it is the LC but normalised so that the value at the maximum is 1.

Fig. 2A shows the estimated LRFs for the Zhang *et al*. microarray data but where we have translated them in time so that each LRF’s highest peak is at 12noon. When we compare many LRFs visually we usually centre then in this way because this makes comparison of the shapes easier. Many examples of estimated LCs and LRFs can be seen in in Figs. 1-5 and the SI. the resulting predicted timing plotted against the sample time is shown in Fig. 2A together with the times corrected to allow for the chronotype explained in Sect. 2.3.1. LCs for the Bjarnason *et al*. human training data are shown in Figs. 2C and S14. These show the general form of the LCs and demonstrate that one can clearly observe qualitative differences between one individual’s LC and those of the others.

**Figure 2:**
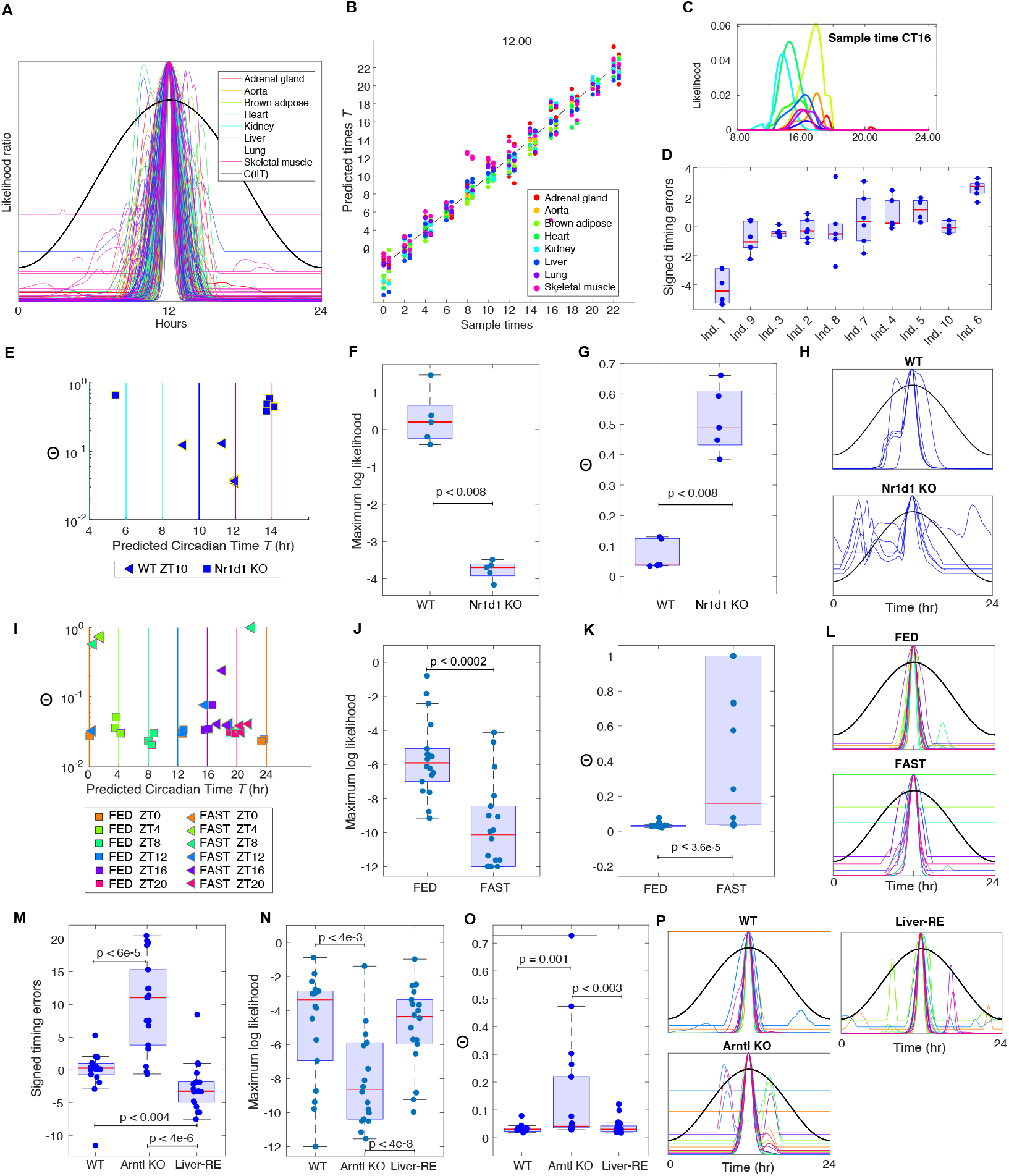
**A**. The centred LRFs for the Zhang *et al*. mouse microarray data. They are centred in that the maximum of the curve is moved to noon. This makes the shapes of the curves clearer and more comparable. The black curve is the curve *C*(*t*|*T*) for *T* = 12 (Sect. S2.2) that is used in the calculation of Θ. Θ is the proportion of time that the LRF spends above this curve. **B** Analysis of the timing for the Zhang *et al*. mouse microarray data with each data point assigned the color corresponding to the tissue. For each time the points over the sample time are for uncorrected timing and the points moved slightly to the right show the corrected timings (i.e. the predicted timing when corrected by the timing displacement for the tissue). A leave-one-tissue out cross-validation approach was used. **C**,**D**. Analysis of the Bjarnason *et al*. data. **C**. Examples of the likelihood curves. **D**. Boxplots showing the timings errors found for each individual ordered by their means. This shows the substantial timing displacements of some individuals. **E-H**. Analysis of the Fang *et al*. data using *l*_thresh_ = −5. **E**. Plots of the Θ value against the estimated time *T*. The vertical lines show the true time with colours indicating the sampling time. WT timings are close to the true sample times and the KO times deviate from them. **F &N G**. Boxplots of the maximum likelihood and Θ values showing significant differences between the WT and KO groups with p-values from the Wilcoxon rank sum test calculated using the Matlab ranksum function. Note that the smallest MLs are around *e*^−4^ which is why *l*_thresh_ was taken to be -5. Taking *l*_thresh_ = −4 gives entirely similar results. **H**. The centred LRFs for the WT and KO samples. **I-L** Analysis of the Kinouchi *et al*. skeletal muscle data. This analysis used a logthresh of −12. The plots J-L are as for F-H but for the Kinouchi *et al*. data. **I-L** Analysis of the Koronowski *et al*. data. comparing WT, Arntl KO and Liver-RE data using *l*_thresh_ = −12. **I** The signed error boxplots show the timing dysfunction in the KO data as well as good recovery in the reconstituted Liver-RE clock but with a clear phase advance. **J**,**K**,**L** Boxplots of ML and Θ values, and centred LRFs for the three genotypes.

#### 2.3.1 Timing errors and timing deviations in the training data

For each training dataset we used an appropriate leave-one-out cross-validation approach to compare the actual time *T*_*a*_ of sample collection with the estimated time *T* and evaluated the timing errors *T − T*_*a*_ for each sample. The mean absolute timing errors (MAEs) for the training datasets are shown in Table 1.

We then analysed how the mean timing error varies with tissue, individual or condition to see if there is a consistent timing deviation for any of these. When these deviations are clear and statistically significant we call the mean of them the *timing displacement* of the tissue, individual or condition. We show below that in all but one of our training datasets the observed timing displacement are associated with coherent phase changes in the genes. Therefore, in assessing the performance of TimeTeller the timing errors should be corrected to take account of this. The timing displacements of the different mouse tissue in the Zhang *et al*. data are relatively small (Fig. S10F) but, for the more genetically heterogeneous human population of the Bjarnason *et al*. data, we found significant timing displacements on the individual level (Fig. 2E). When the apparent errors are adjusted for this they are often substantially reduced (Fig. 2D and Table 1). For the Bjarnason *et al*. human data this reduction is of the order of 50%. Table 1 shows that timecourse and timecourse then intergene (both) normalisations are performing significantly better than intergene alone.

It is difficult to compare performance with that of the published algorithms mentioned above as they have been used on different datasets collected under different conditions and there has been relatively little work on time-stamped genetically heterogeneous data. The Zhang *et al*. dataset was also analysed by ZeitZeiger and the mean absolute errors on cross-validation were between 0.6h and 1.1h [16]. On these tissues the results for timecourse normalisation with TimeTeller are very similar to those of ZeitZeiger (Table S6). Moreover, TimeTeller’s timing errors for the genetically heterogeneous human data compare well with those found in other studies which typically have a median absolute error (MdAE) greater than 1.4h. For example, in the study [24] the 1-sample method had a MdAE of 1.6h and the 2-sample method had a MdAE of 1.4h-1.7h and when CYCLOPS was validated against pre-frontal cortex biopsies with annotated time in [3] the MdAE was 1.69h. In an impressive application to data from four distinct human studies TimeSignature [19] reported MdAEs between 1.21h and 1.49h although this requires two samples for each individual. While TimeTeller’s timecourse normalised results for the genetically heterogeneous Bjarnason *et al*. and Mure *et al*. data (Table 1) compare favourably with these results we do not wish to claim timing superiority as there is great heterogeneity in the studies giving rise to the data that was analysed and in the transcriptomics platforms employed.

#### 2.3.2 Maximum likelihood ML

Given a test sample REV *g*, the value of ML = *L*_*g*_(*T*) (i.e. the maximum likelihood of *g*) is a key diagnostic as, if *M* denotes the maximum value of the distribution *P*(*·*|*T*), we can regard *λ* = log(ML*/M*) as a likelihood ratio test statistic for a pure significance test of the hypothesis that *g* is drawn from the training clock. Thus a low value of ML relative to the values obtained by training or control data is indicative of the fact that *g* comes from a clock that is substantially different. We refer to dysfunction of this kind as *low ML* (lowML). An initial evaluations of the ML values for both training and test data is a key first step of an analysis using TimeTeller.

#### 2.3.3 Dysfunction metric Θ

Statistical theory tells us how to estimate the confidence interval for the maximum likelihood estimator *T* for any given degree of confidence using the LRF (Sect. S4). The variance of *T* arises because *g* is a random sample from the clock at time *t* and we want to know how *T* will vary with other such samples because high variance implies imprecise timing. We call such dysfunction *high variance timing* (highTvar). The Cramér-Rao Theorem [35] gives a lower bound for variance in terms that can be related to the LRF (Sect. S4). Our metric Θ is the proportion of time in the day that the LRF spends above the curve *C*(*t*|*T*) defined in Sect. S2.2) and is associated with the length of such a confidence interval (Sect. S4) and therefore Θ gives an assessment of this sort of dysfunction and higher Θ is associated with higher dysfunction.

However, our likelihood curves often contain structures that are relevant to assessing dysfunction but which are not covered by this aspect of statistical theory. For example, when *g* has significant dysfunction of type lowML, Θ, as defined by us (Sect. S2.2), can also contain a contribution from this. Moreover, the LC and LRF can contain a secondary peaks that have a lower likelihood than that at *T* and this can contribute to Θ. We discuss these aspects in Sect. 2.4 after discussing the values of Θ and ML in training data.

#### 2.3.4 Θ and ML for training data

To continue the evaluation of TimeTeller’s LCs, and the corresponding dysfunction metrics Θ and ML we firstly tested it on the Zhang *et al*. and Bjarnason *et al*. training datasets using the appropriate leave-one-out cross-validation approach. In particular, we asked if there was consistently low Θ values and relatively high maximum likelihoods associated with the good clock function (GCF) of typical WT, control and healthy tissue. The results showed such consistency across tissues for the genetically homogeneous mouse datasets and genetically inhomogeneous individuals for the human data (Figs. S10D,E S15 and S32). A boxplot of the Θ values found in the leave-one-out analysis of the Bjarnason *et al*. human data shows the relative uniformity of the Θ values which have low value similar to that found in other healthy human data studied by us (Figs. 3C,I, S15).

**Figure 3:**
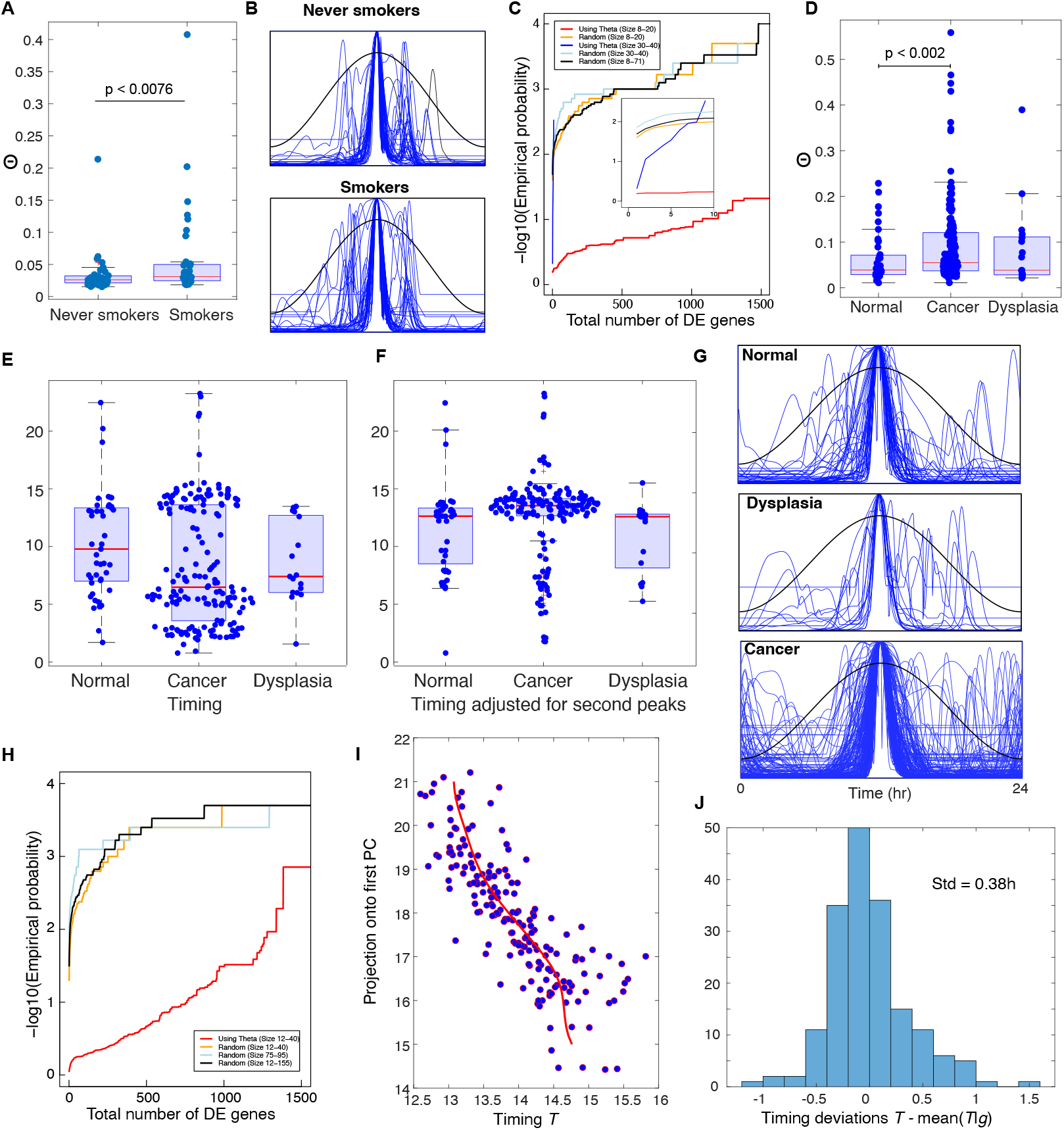
A-C. Analysis of the Boyle *et al*. data. **A**. Boxplots of the Θ values for the smoker and never smoker individuals showing a statistically significant difference in the distributions. There is no statistically significant (Wilcoxon test) difference for the maximum likelihoods (Fig. S21A). **B**. The centred likelihood curves for the smokers and never smokers. **C**. The black, orange and light blue curves are estimates of the probability *p*_rand_ for random choices of the bad clock group of different sizes as in the legend. The red curve is for *p*_Θ_(*m*). There were 5000 iterations for each curve shown. which gave a similar result to 10,000. The number of DE genes was decided using the BH adjustment method with p *<* 0.05 without any restriction on the minimum log fold change. The inset shows a blow up of these curves for *m ≤* 10. From the blue curve (*p*_Θ_(*m*) for 30 *≤ m ≤* 40) we see that for this range of *m* (unlike 8 *≤ m ≤* 20) it is very likely that only a very small number of DEGs are found. For the 66% of cases where a DEG is found there is a 99% chance that *PER3* is among them and a 68% chance of *NR1D2* being present. **D-I. Analysis of the Feng *et al*. data. D**. Boxplots of the Θ values for the samples from individuals in the normal, cancer and dysplasia subgroups. These show a statistically significant (Wilcoxon test) difference in the distributions between the normal and cancer groups and the cancer and combined normal and dysplasia subgroups. **E**. Boxplots showing the predicted timing of the samples. **F**. Boxplots showing the predicted timing when all samples timed as before 7am and with a second peak are given the timing of the second peak. Of the 108 such samples 93 have moved. This suggests that the mistimed samples are primarily so because the wrong peak has a higher likelihood. **G**. Centred LRFs for the three subgroups. **H**. A study of differential effects between between those *n* individuals with the worse clocks according to the Θ stratification and those with better clocks. The black, orange and light blue curves are estimates of the probability *p*_rand_ as in C above but for the Feng *et al*. data. The red curve is for *p*_Θ_(*m*). **I**.Scatter plot of the projection 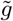 of each REV in the Feng *et al*. data with 12 *< T <* 16 (after using the second peaks if the first gives *T <* 7) against timing *T*. The red curve is a kernel smoothed estimate of the mean of 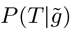. **J**. Distribution of the deviations in H. For each data point this is the horizontal difference between the data point and the red curve. A simple analysis shows that this is largely independent of 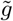 and hence its standard deviation can be used as an upper bound for that of *P*(*T* |*g*).

### 2.4 Multiple tools for assessing functionality in test data

Importantly, this consistency of good timing errors, high ML and low Θ for WT/control test data was also observed in the analysis of the various test data sets considered below and in the SI. For example, using the Zhang *et al*. microarray data for training and intergene normalisation our analysis of the microarray timecourse dataset created by LeMartelot *et al*. [36] (Sect. S1.2) produced a mean absolute error for time estimation of less than one hour and Θ values similar to those found in the training data (Fig. S18). Similar results were found for the Acosta-Rodríguez *et al*. data [37] for *ad libitum* fed mice using the Zhang *et al*. RNA-seq data for training and timecourse-matched normalisation (Fig. 5), and for liver microarray test data from Hughes *et al*. [38] after training with the Zhang *et al*. microarray data (Fig. S19). For the latter we used timecourse normalisation for the training and test data as the microarray platforms are different, demonstrating good results across different platforms.

To further test the use of TimeTeller across different transcriptomics platforms we carried out a cross-validation experiment where we trained TimeTeller on the Zhang *et al*. microarray data and used this to test the Zhang *et al*. RNA-seq data and vice-versa (Figs. 1F, S11 and S12. This not only tests the robustness of our approach but also examines the effectiveness of timecourse normalisation in allowing us to work across different transcriptomics technologies. The timing results are given in Table 1F,G with small mean and median errors of a size compatible with the within-dataset leave-one-out analysis. As well as the relatively small timing errors we observe informative visualisation and consistent Θ values.

Across the various datasets we consider, this good timimg, high ML and low Θ for WT/control test data almost always differed from the perturbed test data. For example, in Fig. 2H-O we consider the timing, ML and Θ diagnostics for the Fang *et al*. and Kinouch *et al*. test data discussed above and use them to illustrate how to gain more insight into dysfunction. In such an analysis one should start with an assessment of the MLs and an inspection of the LRFs for training, control and test data.

From this one can choose an initial value for the important parameter *l*_thresh_ using the approach described in Sect. S2.3. This parameter plays an important role in ensuring that the incorporation of the local likelihoods into a global one (see Sect. S2.2 is not wrecked by inaccurate and uninformative exceptionally low local likelihoods. Briefly, *l*_thresh_ should be chosen as large as possible subject to two conditions: firstly that very few training and control samples have ML smaller than 0.14 exp(*l*_thresh_) which is equivalent to the LRF not having flat regions that significantly intersect *C*(*t*|*T*) and thus contributing to Θ, and, secondly, such that as much as possible of the test data should have MLs above *exp*(*l*_thresh_). The reason for keeping it as large as possible is that taking it too small can remove any structure in the LRF that is not close enough to the peak at *T* and this looses information. This is explained in Sect. S2.3.

For the Fang *et al*. data and the Kinouch *et al*. skeletal muscle test data we see that the ML values for the perturbed test data are significantly lower than those for the control test data (Fig. 2J,N) suggesting that dysfunction of the lowML type is present in the perturbed systems. For the test data from Fang *et al*. we also observe significant differences between the WT and *Nr1d1*KO observations for the timing and Θ diagnostics (Fig. 2H-K). The timing *T* of the KO observations is significantly further from the true timing and they consistently overestimate the sample time (Fig. 2H). Analysis as in Sect. S2.3 suggests setting *l*_thresh_ around -5 but the results are very similar for any value between -4 and -7. However, the centred LRCs also indicate that there is a significant amount of highTvar dysfunction in the KO sample because of the second peaks and increased width of the LRCs near the maximum.

For the Kinouchi *et al*. skeletal muscle data the FED samples show uniformly small errors in timing *T* (MAE 0.44h) and uniformly low Θ values (Fig. 2L,M). On the other hand, for the FAST samples the timing *T* values are clustered around ZT 18-24 reflecting the tight clustering seen in the visualisation (Fig. 1E) and the MAE is significantly greater at 4.04h. So far as dysfunction is concerned, the situation is somewhat different from the Fang *et al*. data since, although there are also significant differences in ML and Θ values between FED and FAST (Fig. 2I-L), there are no second peaks in the centred LRFs contributing to Θ (Fig. 2L). Consequently, the primary difference between the FED and FAST samples is due to the significant difference in the MLs. Thus, only substantial lowML type dysfunction is present. The stratification by Θ in Fig. 2K reflects this.

The Kinouch *et al*. data provides a very informative example of how the choice of the parameter *l*_thresh_ works because the maximum likelihoods ML for both the FED and FAST liver data are significantly higher than that for the skeletal muscle data discussed above and this means that different values of *l*_thresh_ are appropriate. The discussion in Sects. S2.3 and S9.4 shows that the value for the liver data should be substantially larger at -6 or -7 rather than -12. When this value is chosen the results for the Kinouchi *et al*. liver data (Sect. S9.4) are similar to those above for the skeletal muscle data (Fig. S16).

In order to understand the effect of fasting on the amplitude of core clock components Kinouchi *et al*. [34] needed to treat the data as though the FAST samples belonged to a continuous time series even though each timepoint was proceeded by 24 hours of starvation. This underlines a significant extra advantage of TimeTeller because the FAST test samples can be considered independently from one another.

Koronowski et al [39] compared the liver transcriptomes of wild-type (WT) with whole body *Arntl* deficient mice (KO) or *Arntl* KO mice with liver-specific *Arntl* reconstitution (Liver-RE). Their data enables us to test TimeTeller’s sensitivity to not only the substantial KO perturbation but also the much subtler one of the Liver-RE. Our analysis shows statistically significant differences in timing between WT, KO and Liver-RE including the phase advancement noted in [39] of the Liver-RE clock relative to WT (Fig. 2M-P). The MLs (resp. Θs) for the KO data are significantly smaller (resp. larger) than for both the WT and Liver-RE data with no significant difference between WT and Liver-RE (Fig. 2N,O). However, the LRFs (Fig. 2P) suggest a clear difference between WT and Liver-RE data in that, unlike WT, about half of the Liver-RE samples have a significant extra peak suggesting a contribution of highTvar type dysruption and a hypothesis that this is causing the observed timing change in the Liver-RE data. The KO data also has about half with significant second peaks suggesting a combination of some highTvar dysfunction combined with the significant lowML dysfunction.

#### 2.4.1 Analysing stopped clocks

In the SI (Sect. S9.13) we discuss an analysis of two studies (Weger *et al*. [40] and Yeung *et al*. [41]) where the clock is disrupted when either the gene *Arntl* (*Bmal1*) or both genes *Cr1/Cry2* are deleted out in the mouse liver (see Fig. S33). As well as confirming the observation in Hughey *et al*. [16] that the data shows clustering to a narrow range of apparent times for the KO samples, the TimeTeller Θ and maximum likelihood values provide quantitative evidence about the dysfunction caused. This is similar for the two *Arntl* KO datasets but different to that of the *Cry1/Cry2* KO dataset. For the two *Arntl* datasets the KO samples have significantly reduced ML values and significantly increased Θ values and inspection of the centred LRFs show that almost all the contribution to Θ in the KO samples comes from flat regions in the LRFs. It follows that the dysfunction is primarily of lowML type with the KO data having moved away from the training clock in a way that gives consistently wrong times. On the other hand, the *Cry1/Cry2* KO samples though having radically wrong timing have similar high ML and low Θ values to the control data (Fig. S33I,L,O) and this confirms the visualisation that the KO data sits remarkably close to the mean trajectory of the training clock in a way that indicates that its dysfunction is just in the timing. We therefore hypothesise that the *Cry1/Cry2* KO clock is “frozen” in a particular state very close to a wild-type clock state because it has undergone a SNIC bifurcation (see below). This is an extreme example of where there is significant timing dysfunction where the clock reliably gives the same wrong time but no dysfunction of the lowML or highTvar types. We call this dysfunction type *reliable wrong timing* (relTwrong). We will see other examples of this below where the clock is not stopped.

In a deterministic dynamical system, when a parameter is changed slowly there are only two generic ways that oscillations are killed: the Hopf bifurcation where the amplitude declines to zero, and the saddle-node SNIC bifurcation where, until the bifurcation occurs, the amplitude of the oscillation is maintained but at the bifurcation the system stops at a point on the system’s limit cycle [42]. This insight and the quantification results from TimeTeller suggest our hypothesis that mice deficient in *Cr1/Cry2* have undergone a SNIC bifurcation in the liver clock.

### 2.5 The potential for the use of the Θ stratification to identify differential effects in patients

It is particularly interesting to apply TimeTeller to genetically heterogeneous human data because it allows us to test the idea that it can uncover corresponding heterogeneity in the “clock” phenotype or effects on individuals such as patients.

We firstly consider data from a study of the effects of cigarette smoke on the human oral mucosal transcriptome, In this study (Boyle *et al*. [43] & Sect. S1.5), transcriptomes from buccal biopsies of 39 current smokers (*≥* 15 pack-year exposure) and 40 age- and sex-matched never smokers (*<* 100 cigarettes per lifetime) were analysed and compared. The authors found that smoking altered the expression of numerous genes but none of those found were core clock genes nor did they consider the effect of smoking on the circadian clock. They found smokers had increased expression of genes involved in xenobiotic metabolism, oxidant stress, eicosanoid synthesis, nicotine signaling and cell adhesion and decreases were observed in the genes *CCL18, SOX9, IGF2BP3* and *LEPR*. On the other hand, it has been reported elsewhere that smoking has an impact on multiple sleep parameters and significantly lowers sleep quality [44, 45, 46] and this was confirmed in an experimental study which also correlates poor sleep to inflammation [47] while inflammation has been linked to clock disruption. Moreover, CS exposure has been shown to cause circadian disruption in the lungs of WT mice and this is exaggerated in the *Nr1d1* knockouts [48] and has a connection to *Arntl* [49].

Interestingly, when analysed by TimeTeller (Fig. 3A-C) we see a clear and statistically significant difference between the Θ values of the never smoked and smoking individuals (Fig. 3A) which is reflected in the 3D visualisation (Fig. S20). Inspection of the LRFs show that the variations in Θ come mainly from second peaks rather than low ML (fig. 3B). Indeed, the ML values for smokers and non-smokers were not significantly different although the smokers had more observations with a very small ML (Fig. S21A). The lowest values were around *e*^−11^ suggesting that a *l*_thresh_ of about -12 would be appropriate.

A significant proportion of the smokers had Θ values similar to those of the never-smokers but many had much higher values (Fig. 3A). Therefore, we asked if we could identify differential gene expression between the individuals with high Θ versus those with lower Θ. To do this we tested for differential gene expression between the *n* worst clocks (the bad clock group (BCG)) and the others (good clock group (GCG)) adjusting the p-value appropriately to allow for the multiple testing. For a fixed *l*_thresh_ in the range from 11 to 13 with *n* between 8 and 20 we found many differentially expressed genes (DEGs) at the appropriately adjusted *p* = 0.05 level including some clock genes (Fig. S23). However, the particular genes found were sensitive to changing the value of *l*_thresh_ among the suggested values of -11, -12 or -13 and changing the group size *n*.

On the other hand, we calculated that the probability of finding such numbers of DEGs by chance was extremely small and we noticed significant differences between the behaviour when the BCG size was in the range 8 to 20 from that when it was 30 to 40. Therefore, we estimated by simulation the probability *p*_rand_(*m*) of finding *m* or more DEGs by chance when we choose a random group of *n* individuals for our BCG and compared this to the probability *p*_Θ_(*m*) of finding *m* or more DEGs when the stratification by Θ is used to choose the BCG and *l*_thresh_ and *n* are chosen randomly in the ranges -11 to -13 and 8 to 20. We find that uniformly in *m, p*_Θ_(*m*)*/p*_rand_(*m*) *>* 100 (Fig. 3C). We get an interestingly different result if we instead let the group size *n* range between 30 and 40. The probability *p*_rand_(*m*) behaves in approximately the same way but *p*_Θ_(*m*) does not (Fig. 3C(inset)). For *m* very small *p*_Θ_(*m*) is high but as *m* increases *p*_Θ_(*m*) rapidly decreases to values much smaller than those for *p*_rand_(*m*). There is a 34.26% chance of getting no DEGs but when this is not the case there is a more than 99% chance of getting the gene *PER3* and a 68% chance of getting *NR1D2*. Thus, this analysis identifies two interesting groups of individuals with a nontrivial transcriptional phenotype that distinguishes them from the individuals with good clocks. One of these groups appears to be associated with differential expression of *PER3* and *NR1D2*, genes not identified in the original paper where all non-smokers and smokers were compared.

In conclusion, any link between smoking and clock dysfunction is likely to be complex but these results suggest that in a genetically heterogeneous population where the effects of a perturbation such as smoking are likely to be diverse, TimeTeller’s Θ stratification can help identify individuals or groups where the effect is significant.

As a final example of this section we consider the distribution of Θ values by disease state for the transcriptomic data of healthy or dysplastic oral mucosa and oral squamous cell carcinoma (OSCC) from Feng *et al*. [50]. Since the lowest ML values were around *e*^−11^ a *l*_thresh_ of -12 was used. The ML values for normals and cancer were not significantly different although the cancer group had more observations with a very small ML (Fig. S22). However, there is a highly significant difference in median Θ values between the cancer group (167 individuals) and the the normal mucosa group (45 individuals) (p < 0.002) (Fig. 3D). Moreover, there appears to be significant dysfunction in terms of timing estimation (Fig. 3E) that can be significantly ameliorated if the second peaks in the LRF is used for timing when the first peak is clearly misleading (Fig. 3F, details below). Inspection of the LRCs (Fig. 3G) shows that, as for the Boyle *et al*. data, the variations in Θ come mainly from second peaks rather than low ML.

As for the Boyle *et al*. data we asked if there are differentially expressed genes between the worst clocks in the cancer group (high Θ) and the best clocks within the same group and carried out a similar analysis. For genes in general and BCG sizes *n* between 12 and 40 we find similar results with *p*_Θ_(*m*)*/p*_rand_(*m*) *>* 100 for the number *m* of DEGs between 2 and 1200 (Fig. 3H). Many of these DEGs are associated with gene signatures such as DNA repair, E2F targets, G2M checkpoint and the mitotic spindle. However, we do not find any groups like that for the Boyle *et al*. data (with *n* between 30 and 40) that have very low numbers of specific DEGs.

A study of the estimated timing *T* for this data (Fig. 3E) was very informative. The estimates for the normal data are generally between 7 am and 3 pm. A large number of cancer samples have unlikely times well outside the normal working day and the median is clearly much too early. Interestingly, it appears that the mistimed samples are primarily so because the likelihood curve has a second peak (Fig. 3G) and the peak giving an unreasonable timing estimate is slightly higher than one giving the best estimate. In fact, there are 109 samples whose timing *T* is before 7am and 93 of these have a second peak. if we replace the timing by that given by the second highest peak, the great majority moved to a time firmly in the early afternoon between 12noon and 4pm (Fig. 3F). As a result 74% of all samples then fall in this time slot and only 7% remain before 7am. The analysis in the next section indicates that this corrected timing is likely to be the correct time of sampling to within approximately 0.4h.

### 2.6 TimeTeller’s precision on untimed cancer data

The only method currently utilised to estimate the precision of timing/phase algorithms is to use time stamped data and compare the algorithm’s predicted times *T* with the external time stamps *t* when the samples were acquired. However, such a measure of precision is problematic when the individuals, tissues or conditions have a nontrivial molecular chronotype as is the case with the human data considered here and can’t be done if the data is not time stamped. A related test which avoids these problems is instead to determine the variance or standard deviation of the distribution *P*(*T* |*g*) where *T* is the predicted time and *g* is the relevant gene expression vector (Sect. S10). This tells us how well the algorithm calculates the time given the gene state. Interestingly, we can calculate this precision measure even in some cases where we have no timing data and where there is dysfunction and the Feng *et al*. data gives a very informative example of this.

To illustrate this we study the 77% (176 samples) of that data for which the estimated timing *T* after adjustment by second peaks is between 12 noon and 4pm (Fig. 3F). We ask if within this data we can see coherent timing structure or not. A positive answer is suggested by the fact that during this time period the clock genes *NR1D2* and *ARNTL* are changing significantly with time, with *NR1D2* decreasing and *ARNTL* increasing, and if we plot the expression level of these genes against *T*, we see respectively a significant positive and negative slope. Moreover, we can estimate the required standard deviation by carrying out a principal component (PC) analysis of the expression data (see Sect. S10) and plotting the projection of these data onto the first PC against the predicted time *T* (Fig. 3I). The standard deviation of *P*(*T*|*g*) can be estimated from this (Sect. S10) and we obtain an estimate of less than 0.4 hours. If we consider all the deviations from the mean for the timings *T* (give by the horizontal deviation of the relevant data point from the red curve in Fig. 3I) across all of the REVs in this data we obtain the distribution shown in Fig. 3J. Remarkably, although the data is not timestamped and has significant dysfunction giving rise to significant second peaks, TimeTeller is able to accurately measure the time of sampling.

We carried out a similar analysis and found similar results but a bigger standard deviation of 0.83h for the large breast cancer dataset analysed in [51] (Sect. S10). In this case there is no need for adjustment for second peaks as 86% of the data has its predicted time *T* between 10am and 8pm (Fig. S34). We believe this approach gives a new simple method to assess timing performance.

### 2.7 Comparing clocks across individuals, conditions and tissues

Current analyses comparing the circadian clock across individuals, tissues and conditions such as the three studies we consider below proceed by analysing the behaviour of the individual interesting genes separately. Such analyses tend to focus on the level of expression and do not take into account correlations between related genes. We asked whether using Timeteller such an analysis could be done in a more integrated way treating the clock as a system (and hence using correlations) and whether such an approach uncovers some aspects that are hard to see when done gene by gene. The key results here are that it enables us to identify coherent differences in timing across individuals, conditions and tissues and that using these we are able to see in a quantifiable way if the timing differences come from a more or less coordinated change in gene phases.

#### 2.7.1 Using TimeTeller to identify a molecular chronotype

The human training data that we consider involves genetically heterogeneous individuals and therefore we also asked to what extent in this analysis of time-series data we could differentiate systematic variation of timing in an individual or tissue due, for example, to genetic and/or environmental factors, i.e., a molecular chronotype.

We observed above that for the Bjarnason human data, while the Θ and maximum likelihood values are reasonably consistent across individuals, the apparent timing error was not. For some individuals there were substantial timing displacements arising from consistent deviations of the estimated time from the sampling time (Fig. 2D). For example, the individuals labelled 1 and 6 in Fig. 2D have substantial statistically significant (p < 0.003) timing displacements in opposite directions. To further understand this, we hypothesised that the timing displacement of an individual might be largely a result of well-coordinated phase changes in the core clock genes.

If this is the case there should be a definite relation between TimeTeller’s timing error and the phase of the genes. Moreover, since this relationship is local in that the timing displacements are small compared to 24 hours, it is reasonable to suspect that it might be approximately linear. Therefore, we tested for a linear relation between the phase variation of the genes in our panel and timing displacement.

In this analysis we regressed the timing displacement against the phase of each of the genes in the REP (Fig. 4) using Cosinor [52] to measure gene expression phase. For all the probes used we observed an approximately linear relationship between apparent error and the variation in gene phase with a positive slope. For all genes the non-zero slope is statistically significant and the *r*^2^ value is greater than 0.7, and for many genes it is greater than 0.9. This measures the proportion of the variation in the gene phase that is predictable from the TimeTeller displacement using the linear relationship. Thus TimeTeller is able to clearly identify coherent and substantial phase variation in the clock genes for each individual across all genes in the rhythmic expression profile. It identifies a clear “chronotype” for each individual and a quantifiable phase difference. Moreover, the strong coherence between the time estimations and the gene phases is further validation of TimeTeller’s time estimation. These results suggest that if the real clock time of the sample is known, by combining the observation of a Θ suggesting good clock function with an advanced or retarded time prediction, TimeTeller can help identify substantial coherent phase variation in an individual’s clock genes from a single sample.

**Figure 4:**
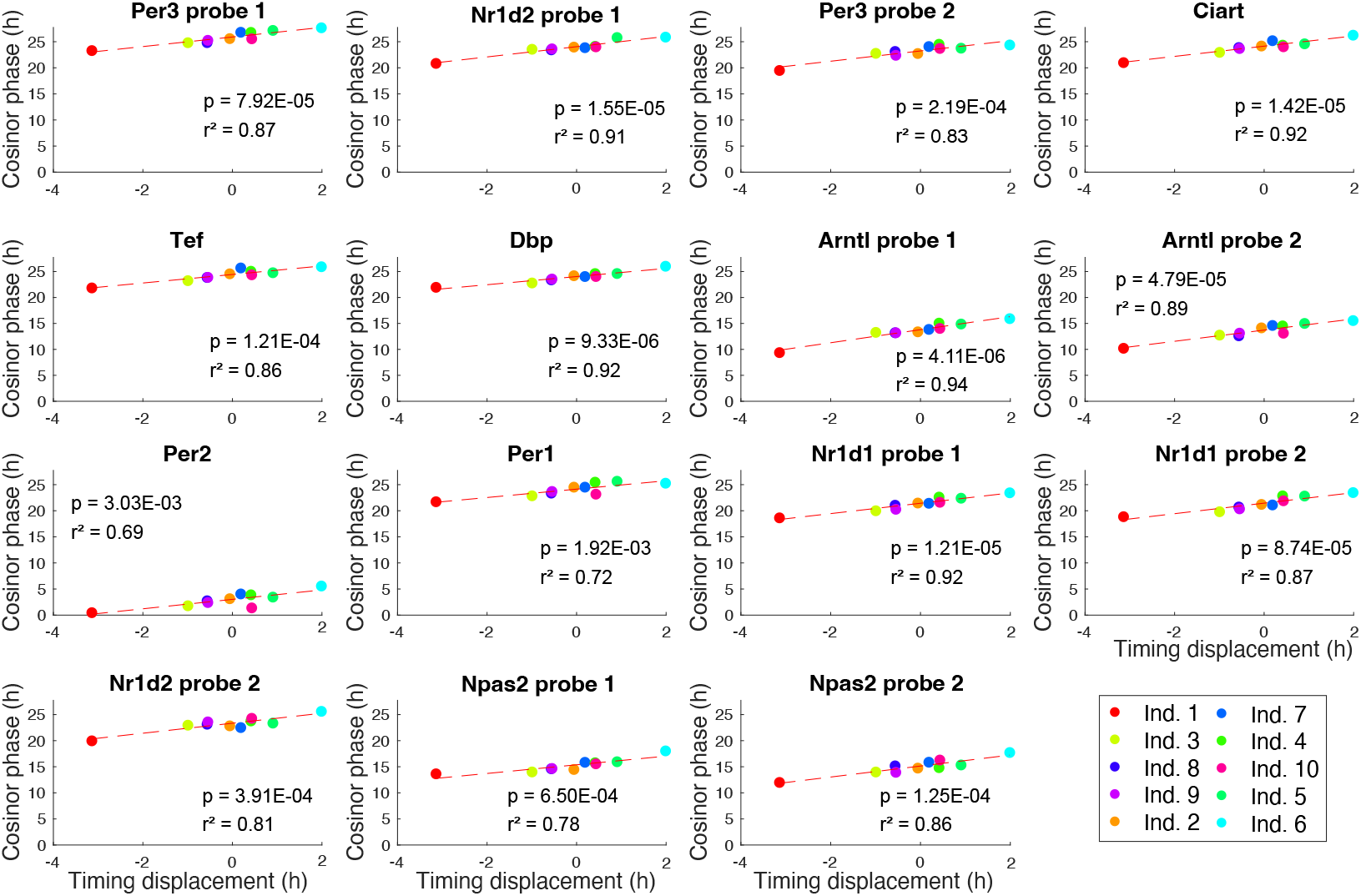
This shows the strong linear relationship between the gene phases and the timing deviation in the Bjarnason *et al*. data. Each point corresponds to an individual. The regression was carried out using Matlab’s fit function and Cosinor [52] was used to estimate the gene phases from the time series of each individual. The *p*-values test the hypothesis the the slope of the line is non-zero and are given by the F-test using the Matlab functions coefTest and fitlm. The *r*^2^ values measure of the proportion of total variation of gene phase explained by the linear model. They indicate the extent to which the linear model (corresponding to a simple phase change) explains the data with the values very close to 1 almost completely explaining it.

We will utilise such regression plots in the analyses below where we attempt to characterise the nature of the change in the clock caused by different conditions or in different tissues. We call such plots *phase/displacement plots* (PDPs).

#### 2.7.2 Timing divergences and clock comparisons for time-restricted feeding in ageing mice

Recently, Acosta-Rodríguez *et al*. [37] studied the synergistic effects of various time-restricted feeding protocols with caloric restriction (CR) on the prolongation of life span in mice focusing on the liver which is a major metabolic target of the circadian clock. After 6 weeks of baseline *ab libitum* (AL) food access, C57BL/6J male mice were subjected to 30% CR. Mice were fed nine to ten 300-mg food pellets containing 9.72 to 10.8 kcal every 24 h starting at the beginning of the day (CR-day-2h) or night (CR-night-2h) constrained to consume their food within 2h.

Two additional CR groups of mice were fed a single 300-mg pellet delivered every 90 min to distribute the food intake over a 12-h window either during the day (CR-day-12h) or during the night (CR-night-12h). A fifth CR group of mice was fed a single 300-mg pellet every 160 min continuously spread out over 24 h (CR-spread). Liver gene expression was profiled using RNA-seq in all six feeding conditions at 6 and 19 months of age. Livers were collected in constant darkness at 12 time points every 4 hours for 48 hours across two circadian cycles. We treat the data from time *t* and *t* + 24 as replicates of a 24h cycle.

Together with a young and old group where feeding was *ad libitum* (AL) this results in 12 feeding conditions. We used TimeTeller to analyse this data asking if it could identify the nature of systemic changes in the core clock between the different feeding*×*age conditions. We used the Zhang *et al*. RNA-seq data as training data. Thus, all feeding conditions of [37] are regarded as test data. We analysed this using the Zhang *et al*. RNA-seq data for training and using both time-course and timecourse-matched normalisation for the test data. The results are very similar and we give the timecourse-matched results here.

Visualisation showed that the test data fell nicely within the trained distribution close to the mean cycle (Fig. S24). Analysis as in Sect. S2.3 point to using a *l*_thresh_ of -8. The results on the predicted times *T* showed a substantial timing displacement (Fig. 5A) for eight of the conditions with CR-day-2h being the most extreme (see Table S7 for the p-values for the differences; only 12 of the 66 comparisons have p *≥* 0.05). Moreover, there is a striking apparent age-related difference for the CR-day-2h feeding conditions in that the timing displacements of the 6 month and 19 month mice differ by over 4 hours (*p <* 0.0001, Table S7).

**Figure 5:**
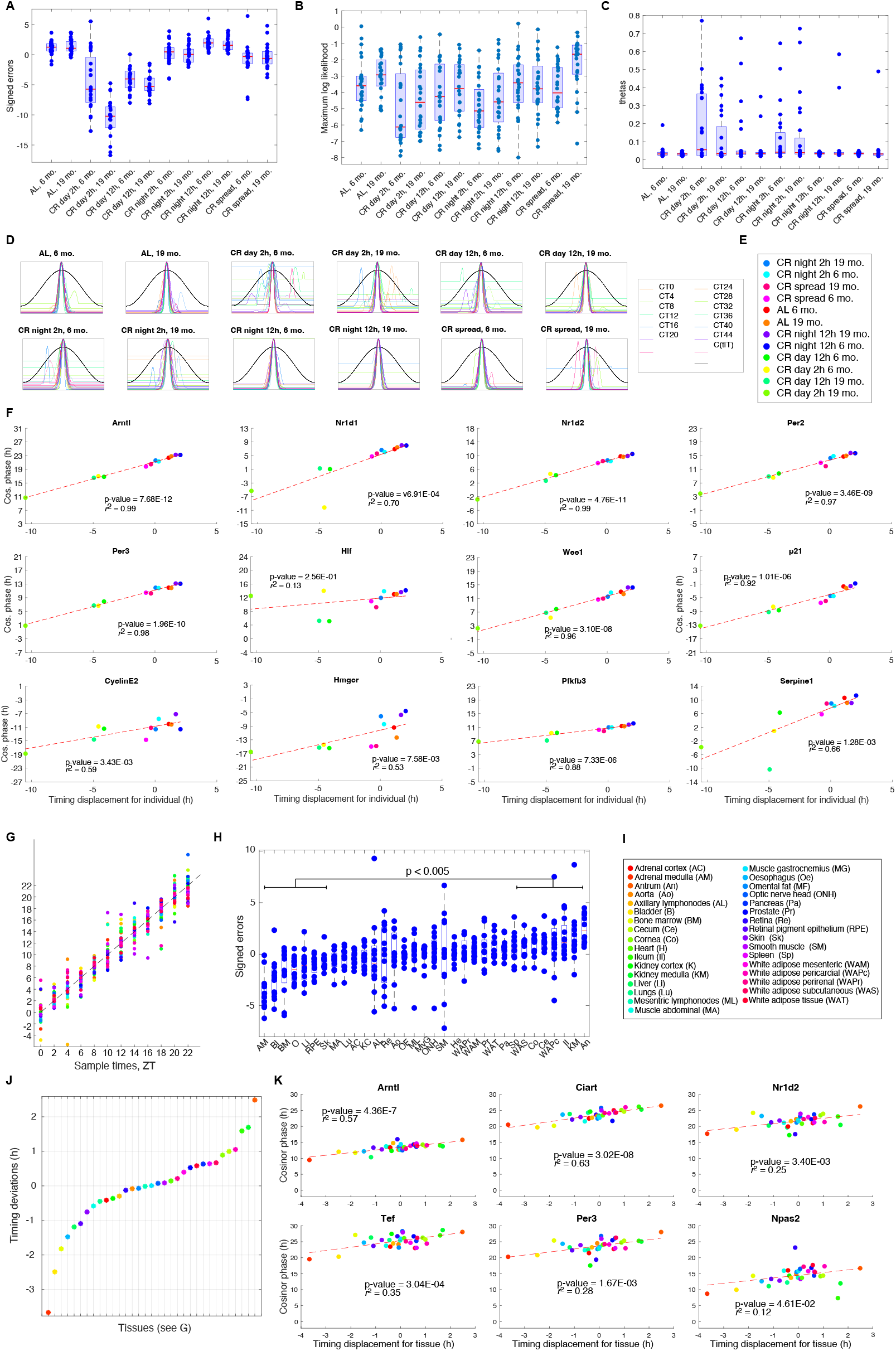
A-E. Analysis of the Acosta-Rodríguez *et al*. data [37]. The training data is the Zhang *et al*. RNA-seq data and this data is analysed as test data using timecourse-matched normalisation and *l*_thresh_ = −8. The timing, Θ and ML for each sample point shown is a suitably averaged value for the two replicates with the same feeding condition and age. **A**. Box plots of the timing error for the different conditions and ages. The mean value of each box plot gives the timing displacement for the condition and age. See Table S7 for statistical analysis of the differences. **B**. Box plots of the ML for the different conditions and ages. **C**. Box plots of the Θ value for each condition and age. See Table S9 for statistical analysis of the differences. **D**. Centred LRFs for each condition and age. **E**. Legend. **F**. PDP plots for the genes in the REP, some cell cycle genes and some of the genes highlighted in [37]. See Sect. S9.10 for more information. The gene phases were measured by Cosinor [52]. **G-K. Analysis of the Mure *et al*. data**. In each plot the color corresponds to the tissue as shown in I. The data from the central tissues is used for training. **G**. TimeTeller predicted time *T* vs the sample time using leave-one-out analysis for each sample from the 33 tissues. **H**. Box plots of the signed errors for the samples for each of the 33 tissues in order of increasing timing displacement. Each of the 7 leftmost boxplots is significantly different from each of the 7 rightmost boxplots at the *p* = 0.005 level. **I**. Legend for G-K. **J**. Timing displacements for each tissue. **K**. Some examples showing PDP plots of the gene phases and the TimeTeller timing displacement for all 33 tissues from Mure *et al*.. The p-value and *r*^2^ values for the other genes are in Table S12. Note that the *r*^2^s for core clock genes are much smaller than those for the Bjarnason *et al*. human data (Fig. 4) and the Acosta-Rodríguez *et al*. data in F.

There are some statistically significant differences between the Θ and ML values found for the different conditions (Fig. 5B,C). These are given in Tables S8 & S9. This is also as noticeable from the centred LRFs (Fig. 5D). For example CR-spread-19m has significantly higher ML values than all other conditions and lower Θ values than most, and CR-night-2h-6m has significantly lower ML values and higher Θ values than all but CR-night-2h-19m (Fig. 5 & Tables S8 & S9). However, overall the ML values are relatively high and therefore confirm the observation that, although the timing can be displaced, the test data is close in data space to the training clock. This is compatible with the hypothesis that the different feeding condition induce a a simple phase change in the clock.

Given these timing displacements, we carried out a comparison of the clocks under the different conditions by analysis using PDP plots where we regressed the phases of the genes against the timing displacements of the various conditions in order to try and quantify the extent to which the observed timing differences are the result of a coherent phase adjustment of each gene (Fig. 5F). For the individuals in the Bjarnason *et al*. oral mucosa data we saw that the differences between the individuals’ clock was primarily of this nature. For the feeding*×*age conditions the situation is even clearer for the core clock genes considered since the *r*^2^ values for them (Fig. 5F) are typically closer to 1 implying the linear model (corresponding to a simple phase change) almost completely explains the data. From this analysis we conclude that it is likely that the different feeding*×*age conditions cause a change in the core clock that is primarily a simple phase change and that for some of the conditions such as CR-day-2h this is substantial. In Fig. S31 we show comparable results for the timing deviation and PCPs where we have used timecourse normalisation for the test data, a different values for *l*_thresh_ and have kept the replicates separate so that the methods consistency can be checked.

In summary, for this data, TimeTeller has enabled the discovery of substantial and coherent differences of the core clock systems state associated to the feeding conditions and provided quantified evidence that the core clocks corresponding to the different conditions differ by a simple phase change. This benefitted from a systems approach. Finally, note that although there is time series data in this instance since we used time-course normalisation and analysed the test data independently having a time series is not necessary.

#### 2.7.3 Timing divergences and clock comparisons for the Mure *et al*. baboon data

We found a different result when we compared the clocks in the different tissues studied in Mure *et al*. [2]. In this paper, the transcriptomes of 64 tissues of the diurnal primate *Papio anubis* (baboon) were analysed from one animal every 2 hours for 24 hours. The results of [2] demonstrate that many ubiquitously expressed genes that participate in essential cellular functions show a tissue-specific rhythmic pattern, and confirmed a shifted temporal organization of central and peripheral tissues between diurnal and nocturnal mammals. Since this RNA-seq dataset involves a genetically heterogeneous population and multiple transcriptionally heterogeneous tissues, we were keen to assess how well TimeTeller was able to analyse it.

We studied 33 of the tissues leaving out those from the brain and some others with missing data. An initial leave-one-tissue-out analysis gave reasonably accurate timing (MdAE around 1h, Fig. 5G, Table reftable1) and indicated that many tissues had a substantial timing displacement (Fig. 5H,J) ranging from approximately -3.5h to +2.5h compared to the time the samples were taken. The standard deviation of the individual sample timing errors around the timing dispacement from a given tissue was generally much smaller than the 6h range of the timing displacements (Fig. 5H). Moreover, the null hypothesis that the *m*th most advanced tissue has the same timing displacement as the *m*th most retarded is rejected at the *p* = 0.01 level for all *m <* 7 (Wilcoxon-Mann-Whitney test).

Given many tissues had large absolute timing displacements, we used only the 18 tissues with the smallest for the training data, i.e., central tissues. This gives slightly better timing results than using all 33 tissues as can be seen in Table 1. Correcting the TimeTeller time predictions by adjusting them using the phase displacements of the tissues resulted in a substantial improvement of about half an hour in the timing accuracy (Fig. 5G & Table 1). Given the heterogeneities in the data this results in a very reasonable performance with a mean absolute error just over one hour.

The analysis of the variation of the core clock across the 64 tissues in Mure *et al*. [2] is mainly concerned with the overall transcript abundance and rhythmicity of expression of the individual core clock genes. The authors note that the heterogeneity of this implies different composition of core activators, repressors, and modulators in different tissues. They do not mention the timing divergences we find in the data using TimeTeller. Using these timing divergences, for the limited set of 33 tissues, we can study this in a different and more integrated way.

As above, we considered a comparison of the clocks in the different tissues by using a PDP plot (Fig. 5K). For this dataset we see that the observed differences between them are not due to a simple coherent phase adjustment in the genes but involves a more complex interaction. This is because the *r*^2^ values, which measure of the proportion of total variation of outcomes explained by the linear model, are very low and much lower than those for the Bjarnason *et al*. and Acosta-Rodríguez *et al*. data. This suggests that the adjustment of the clock from tissue to tissue is more complex than a simple phase shift in the core clock genes. On the other hand, the relatively low p-values suggest that there is a definite correlation between gene phase and timing displacement suggesting that a appreciable component of the changes in the genes is a phase change.

Again this analysis benefitted from a systems approach which enables us to identify coherent differences between tissues and relate this to changes in the core clock.

### 2.8 Probing the effect of changes in the core clock on downstream genes

Changes in the core clock will affect the regulation of rhythmic genes that are downstream of it. Current methods allow one to check whether these genes remain rhythmic when the clock is perturbed in some way but TimeTeller also allows one to check if they maintain their relationship with the clock in a coherent fashion. The way in which the different conditions of the Acosta-Rodríguez *et al*. mouse data [37] changed the phase of the core clock provides a very interesting example where we can demonstrate such an analysis.

The automatically determined REP discussed in Sect. 2.1.1 contained three genes that are not considered to be core clock genes, *Hlf* (Hepatic Leukemia Factor, a bZIP transcription factor), *Cys1* (cystin 1) and *Wee1* (cell cycle regulator of entry into mitosis). In our analysis of the phase change linearity in Fig. 5D we noted that while all the clock genes, *Hlf* and *Wee1* displayed approximately linear phase changes this was not the case for *Cys1* (Fig. 5D). We therefore decided to use this analysis to look at the effect of the clock phase changes observed on some other genes that are rhythmic in the liver of AL fed mice. In particular, we inspected the plots for some cell cycle genes and also a number of the genes identified in Acosta-Rodríguez *et al*. [37] as affected by the CR conditions or ageing.

Of the cell cycle genes *Wee1, p21, P53, Timeless, CyclinA, CHK2, CyclinB1, CyclinE2* and *ATM* it appears that only *Wee1, p21* and *CyclinE2* are rhythmic in the liver in the AL conditions. These three genes maintain coherence with the clock with *Wee1* doing so strongly (*r*^2^ = 0.96) followed closely by *p21* (*r*^2^ = 0.92). The coherence of *CyclinE2* seemed somewhat weaker (*r*^2^ = 0.59). All of the other genes had *r*^2^ *<* 0.4 and appeared incoherent (Fig. S25). This suggests a hypothesis that for the genes *Wee1* and *p21* the primary changes in their regulation induced by the different CR conditions concern only phase change of the core clock.

If we probe the rhythmicity of these cell cycle genes as in Fig. S11 we see that there is a strong correlation between the maintenance of coherence and rhythmicity and, in general in PDP plots, where we see divergence from the regression line we should expect absence of rhythmicity and therefore unreliable gene phase.

In their study [37] Acosta-Rodríguez *et al*. highlighted a number of genes that were affected by ageing or the CR conditions and sorted these into four categories: those susceptible to ageing-related changes under any condition tested, those related to fasting conditions, timing related genes and genes associated with effects on circadian cycling such as rhythmic damping. Our analysis using PDP plots for these genes clearly identifies which of them move coherently with the core clock under the different feeding conditions (Sect. S9.11). None of the timing related genes stayed coherent and, amongst the fasting genes, only *Hal1* (*r*^2^ = 0.66) was. Several ageing genes show some level of coherence (Figs. S26 & S27): *Serpine1* (*r*^2^ = 0.66) *Adora1* (*r*^2^ = 0.68) *Got1* (*r*^2^ = 0.65) *Lepr* (*r*^2^ = 0.68) *Pfkfb5* (*r*^2^ = 0.88). For the genes affecting circadian cycling. while *Gys1* (*r*^2^ = 0.15) and *Per1* were incoherent, the rest were coherent: *Arntl* (*r*^2^ = 0.99), *Nr1d1* (*r*^2^ = 0.70), *Per1* (*r*^2^ = 0.69), *Per2* (*r*^2^ = 0.97) and *Pck1* (*r*^2^ = 0.60). For a significant number of the genes affected by ageing or the CR conditions, while the gene is not coherent under all conditions it is coherent under a significant number of the conditions with the less extreme timing deviations. This can be seen from the PCPs and was the case for *Per1* which seems coherent under all conditions except the four CR-day conditions.

These results demonstrate that such an analysis can give a novel overview of gene response and whether a given gene maintains coherence with the clock when the clock timing changes. Such coherence is associated with genes that show good linearity with a significant slope in the PDP plots. Consequently, TimeTeller can be used to investigate function and dysfunction in genes controlled by the circadian clock when the clock is perturbed.

## 3 Discussion

What we hope stands out is the way we are able to use TimeTeller to study single samples of external test data in ways that reach beyond the information provided by current algorithms. The main aim of this study was to indicate the different ways that TimeTeller can be used to visualise and probe the circadian clock. The visualisation component allows one to get a global picture of clock data and see aspects of its variation and correlation structure as for example in our figures where the ellipsoids give an idea of the correlations as explained in Fig. 1. Although the data mixes intrinsic biological noise with measurement error the visualisation allows some idea of the size of the combined noise. Insights from this can be followed up quantitatively using the software that we will make available. When working with test data the visualisation allows rapid visual comparison of WT/control and perturbed test data to determine how well WT/control data agrees with training data and the extent to which the perturbed data differs. While the visualisations in this paper are static, when using the software the ability to rotate the visualisation is highly informative.

A fundamental assertion is that the TimeTeller likelihood curve contains more information about dysfunction than just timing. We believe that the examples we discuss bring this out. The algorithm’s output is not just limited to a timing estimate alone but comes with an estimate of Θ, ML and the likelihood curve. Thus, one has much more information with which to assess both dysfunction and the assessment’s quality. We can even do a precision assessment on datasets whose timing is unknown.

We give many examples where the dysfunction metrics Θ and ML that we introduce take statistically significant different values in perturbed conditions compared to WT/control. An important aspect of this analytical approach is that Θ can provide a stratification of individual transcriptomes by measured dysfunction. This is important because it enables the possibility of associating clock dysfunction with other aspects of disease on the level of the individual. This is illustrated most clearly by our analysis of the Boyle *et al*. data on the effects of smoking on the transcriptome of the human oral mucosa and that of Feng *et al*. data on oral squamous cell carcinoma. This analysis showed significant differences between the smokers and non-smokers in the Boyle *et al*. data and between normal and cancer for the Feng *et al*. data and in both cases enabled the identification of a “bad clock” group with a significant number of differentially expressed genes compared to other individuals of the same cohort (smoker or cancerous tissue).

When analysing the cancer data samples from Feng *et al*. and Cadenas *et al*. we were able to validate the quality of timing estimates without using any time stamps. This means that we were able to identify a large number of patients with significant dysfunction in the clock but still identify the sample time which for the Feng *et al*. data often involved the second peak in the LRF. Moreover, this method of analysis gives a new way to estimate the precision of timing/phase algorithms on large data sets even if they are not time stamped and even if they contain significant dysfunction as is the case with the Feng *et al*. data. In a future paper we expect to apply TimeTeller to study other cancer datasets.

TimeTeller offers other new possibilities for the analysis of clock data as shown by the analysis of the Bjarnason *et al*., Acosta-Rodríguez *et al*. and Mure *et al*. data. Firstly, TimeTeller allowed us to identify significant timing displacements for the individuals, conditions or tissues that had not been observed and it was not necessary for these data to be in time-series. Secondly, when these are in time series, by identifying the timing displacements and then regressing the gene phases against them, we were able to compare the clock in different individuals, conditions or tissues and attempt to assess whether the difference is largely a phase shift or a more complex adjustment. Moreover, we show how to analyse genes downstream of the clock in a similar way. For example, using the Acosta-Rodríguez *et al*. mouse data we were able to see which genes maintained their rhythmicity and coherence with the clock in all the temporally restricted feeding conditions and which did not.

An important insight of the study of Wittenbrink *et al*. [24] is the need to develope optimised high quality data that is cheap to collect. This will also be important for the use of TimeTeller. While it is clear that the sort of data we discuss in this paper will become increasingly abundant and much cheaper to generate, other data types such as Nanostring’s nCounter platform [53], which might be more suitable to clinical workflows and may be used to provide cheaper purpose-designed datasets that can be used with TimeTeller. This will also bring the opportunity to improve TimeTeller because timecourse normalisation will be less necessary and the training will be improved by having more training data at more timepoints around the day.

The algorithm is very customisable and flexible and relatively fast. The user is free to choose the genes employed by TimeTeller and experiment with the parameters *l*_thresh_, *η* and *ε*. Although we have experimented with changes, the effect of changing *η* and *ε* is clear from equation S2.2 and Fig. S3 and we have seen no reason for changing them from the values we have used here. Keeping them constant means that Θ values can be compared across datasets. On the other hand, *l*_thresh_ needs to be chosen using the data for the reasons explained in Sect. S2.3. To use TimeTeller across datasets and with individuals in a clinical context it will, of course, be crucial to settle on a choice of *l*_thresh_ for the particular type of data being considered. It is also easy to change the internal parameters such as the choice of method for interpolating the covariance matrices around the day and the dimension reducing projection method. We have a preferred choice for these and have used them throughout this paper. Other methods for interpolation may become useful as the TimeTeller is developed. The relatively fast speed of TimeTeller means that it easy for users to experiment with the choices available.

## Acknowledgements

We are especially grateful to Sylvie Giacchetti (Assistance Publique-Hopitaux de Paris) for discussions that were important for motivating the initial development of TimeTeller and for advice on cancer datasets. DAR and DV thank the Engineering & Physical Sciences Research Council (EPSRC) for a PhD studentship through the MOAC Doctoral Training Centre grant number EP/F500378/1. DAR was also supported by Biotechnology and Biological Sciences Research Council (BBSRC) Grant BB/K003097/1 and EPSRC Grant EP/P019811/1. MV, RD & DAR were supported by a grant from Cancer Research UK and EPSRC (C53561/A19933). GAB was supported by The Anna-Liisa Farquharson Chair in Renal Cell Cancer Research. LU and VV were funded by the UK Medical Research Council Doctoral Training Partnership (MR/N014294/1).

## Code and data availability

The software code and data underlying the results of this study are available for reviewers and will be made publicly available on publication.

## References

[1] Zhang R, Lahens NF, Ballance HI, Hughes ME, Hogenesch JB. A circadian gene expression atlas in mammals: implications for biology and medicine. Proceedings of the National Academy of Sciences of the United States of America. 2014;111(45):16219–24. doi:10.1073/pnas.1408886111.

[2] Mure LS, L. HD, Benegiamo G, Chang MW, Rios L, Jillani N, et al. Diurnal transcriptome atlas of a primate across major neural and peripheral tissues. Science. 2018;359(6381):eaao0318. doi:10.1126/science.aao0318.

[3] Anafi RC, Francey LJ, Hogenesch JB, Kim J. CYCLOPS reveals human transcriptional rhythms in health and disease. Proceedings of the National Academy of Sciences. 2017;114(20):5312–5317.

[4] Levi F, Schibler U. Circadian Rhythms: Mechanisms and Therapeutic Implications. Annual Review of Pharmacology and Toxicology. 2007;47(1):593–628.

[5] Lévi F, Okyar A, Dulong S, Innominato PF, Clairambault J. Circadian timing in cancer treatments. Annual Review of Pharmacology and Toxicology. 2010;50:377–421.

[6] Kobuchi S, Yazaki Y, Ito Y, Sakaeda T. Circadian variations in the pharmacokinetics of capecitabine and its metabolites in rats. Eur J Pharm Sci. 2018;112:152–158.

[7] Squire T, Buchanan G, Rangiah D, Davis I, Yip D, Chua Y, et al. Does chronomodulated radiotherapy improve pathological response in locally advanced rectal cancer. Chronobiol Int. 2017;34(4):492–503.

[8] Cordina-Duverger E, Menegaux F, Popa A, Rabstein S, Harth V, Pesch B, et al. Night shift work and breast cancer: a pooled analysis of population-based case-control studies with complete work history. European Journal of Epidemiology. 2018;33(4):369–379.

[9] Shan Z, Li Y, Zong G, Guo Y, Li J, Manson J, et al. Rotating night shift work and adherence to unhealthy lifestyle in predicting risk of type 2 diabetes: results from two large US cohorts of female nurses. BMJ. 2018;363:k4641.

[10] Kettner N, Voicu H, Finegold M, Coarfa C, Sreekumar A, Putluri N, et al. Circadian Homeostasis of Liver Metabolism Suppresses Hepatocarcinogenesis. Cancer Cell. 2016;30(6):909–924.

[11] Cappuccio F, Miller MA, Lockley SW. Sleep, health, and society: From aetiology to public health. Oxford University Press, USA; 2010.

[12] Leger D, Bayon V, de Sanctis A. The role of sleep in the regulation of body weight. Mol Cell Endocrinol. 2015;418 Pt 2:101–107.

[13] Cappuccio F, Miller M. Sleep and Cardio-Metabolic Disease. Curr Cardiol Rep. 2017;19(11):110.

[14] Jike M, Itani O, Watanabe N, Buysse DJ, Kaneita Y. Long sleep duration and health outcomes: A systematic review, meta-analysis and meta-regression. Sleep Medicine Reviews. 2018;39:25–36.

[15] Ueda HR, Chen W, Minami Y, Honma S, Honma K, Iino M, et al. Molecular-timetable methods for detection of body time and rhythm disorders from single-time-point genome-wide expression profiles. Proceedings of the National Academy of Sciences. 2004;101(31):11227–11232.

[16] Hughey JJ, Hastie T, Butte AJ. ZeitZeiger: supervised learning for high-dimensional data from an oscillatory system. Nucleic acids research. 2016;44(8):e80–e80.

[17] Agostinelli F, Ceglia N, Shahbaba B, Sassone-Corsi P, Baldi P. What time is it? Deep learning approaches for circadian rhythms. Bioinformatics. 2016;32(12):i8–i17.

[18] Laing EE, Möller-Levet CS, Poh N, Santhi N, Archer SN, Dijk DJ. Blood transcriptome based biomarkers for human circadian phase. Elife. 2017;6:e20214.

[19] Braun R, Kath WL, Iwanaszko M, Kula-Eversole E, Abbott SM, Reid KJ, et al. Universal method for robust detection of circadian state from gene expression. Proceedings of the National Academy of Sciences. 2018;115(39):E9247–E9256.

[20] Ruben M, Wu G, Smith D, Schmidt R, Francey L, Lee Y, et al. A database of tissue-specific rhythmically expressed human genes has potential applications in circadian medicine. Sci Transl Med. 2018;10(458).

[21] del Olmo M, Spörl F, Korge S, Jürchott K, Felten M, Grudziecki A, et al. Inter-layer and inter-subject variability of circadian gene expression in human skin. bioRxiv. 2022;.

[22] Talamanca L, Gobet C, Naef F. Sex-dimorphic and age-dependent organization of 24 hour gene expression rhythms in human. bioRxiv. 2022;.

[23] Talamanca L, Gobet C, Naef F. Sex-dimorphic and age-dependent organization of 24-hour gene expression rhythms in humans. Science. 2023;379(6631):478–483.

[24] Wittenbrink N, Ananthasubramaniam B, Münch M, Koller B, Maier B, Weschke C, et al. Highaccuracy determination of internal circadian time from a single blood sample. J Clin Invest. 2018;128(9):3826,3839.

[25] Takahashi JS. Transcriptional architecture of the mammalian circadian clock. Nature Reviews Genetics. 2017;18(3):164–179.

[26] Hughey JJ. Machine learning identifies a compact gene set for monitoring the circadian clock in human blood. Genome medicine. 2017;9(1):1–11.

[27] Wu G, Ruben MD, Schmidt RE, Francey LJ, Smith DF, Anafi RC, et al. Population-level rhythms in human skin with implications for circadian medicine. Proceedings of the National Academy of Sciences. 2018;115(48):12313–12318.

[28] Shilts J, Chen G, Hughey JJ. Evidence for widespread dysregulation of circadian clock progression in human cancer. PeerJ. 2018;6:e4327.

[29] Bjarnason G, Seth A, Wang Z, Blanas N, Straume M, Martino T. Diurnal rhythms (DR) in gene expression in human oral mucosa: Implications for gender differences in toxicity, response and survival and optimal timing of targeted therapy (Rx). Journal of Clinical Oncology. 2007;25(18_suppl):2507–2507.

[30] Relógio A, Westermark PO, Wallach T, Schellenberg K, Kramer A, Herzel H. Tuning the mammalian circadian clock: robust synergy of two loops. PLoS computational biology. 2011;7(12):e1002309.

[31] Minas G, Rand DA. Long-time analytic approximation of large stochastic oscillators: Simulation, analysis and inference. PLoS Computational Biology. 2017;13(7):e1005676.

[32] Fang B, Everett LJ, Jager J, Briggs E, Armour SM, Feng D, et al. Circadian enhancers coordinate multiple phases of rhythmic gene transcription in vivo. Cell. 2014;159(5):1140–1152.

[33] Cho H, Zhao X, Hatori M, Yu RT, Barish GD, Lam MT, et al. Regulation of circadian behaviour and metabolism by REV-ERB-α and REV-ERB-β. Nature. 2012;485(7396):123–127.

[34] Kinouchi K, Magnan C, Ceglia N, Liu Y, Cervantes M, Pastore N, et al. Fasting imparts a switch to alternative daily pathways in liver and muscle. Cell reports. 2018;25(12):3299–3314.

[35] Casella G, Berger RL. Statistical inference. vol. 2. Duxbury Pacific Grove, CA; 2002.

[36] Le Martelot G, Canella D, Symul L, Migliavacca E, Gilardi F, Liechti R, et al. Genomewide RNA polymerase II profiles and RNA accumulation reveal kinetics of transcription and associated epigenetic changes during diurnal cycles. PLoS biology. 2012;10(11):e1001442.

[37] Acosta-Rodriguez V, Rijo-Ferreira F, Izumo M, Xu P, Wight-Carter M, Green C, et al. Circadian alignment of early onset caloric restriction promotes longevity in male C57BL/6J mice. Science. 2022;376(6598):1192–1202.

[38] Hughes ME, DiTacchio L, Hayes KR, Vollmers C, Pulivarthy S, Baggs JE, et al. Harmonics of circadian gene transcription in mammals. PLoS genetics. 2009;5(4):e1000442.

[39] Koronowski KB, Kinouchi K, Welz PS, Smith JG, Zinna VM, Shi J, et al. Defining the independence of the liver circadian clock. Cell. 2019;177(6):1448–1462.

[40] Weger BD, Gobet C, David FP, Atger F, Martin E, Phillips NE, et al. Systematic analysis of differential rhythmic liver gene expression mediated by the circadian clock and feeding rhythms. Proceedings of the National Academy of Sciences. 2021;118(3).

[41] Yeung J, Mermet J, Jouffe C, Marquis J, Charpagne A, Gachon F, et al. Transcription factor activity rhythms and tissue-specific chromatin interactions explain circadian gene expression across organs. Genome research. 2018;28(2):182–191.

[42] Guckenheimer J, Holmes P. Nonlinear oscillations, dynamical systems, and bifurcations of vector fields. vol. 42. Springer Science & Business Media; 2013.

[43] Boyle JO, h Z, Kacker A, Choksi VL, Bocker JM, Zhou XK, et al. Effects of cigarette smoke on the human oral mucosal transcriptome. Cancer prevention research. 2010;3(3):266–278.

[44] Liao Y, Xie L, Chen X, Kelly BC, Qi C, Pan C, et al. Sleep quality in cigarette smokers and nonsmokers: findings from the general population in central China. BMC Public Health. 2019;19(1):1–9.

[45] Lee YY, Lau JH, Vaingankar JA, Sambasivam R, Shafie S, Chua BY, et al. Sleep quality of Singapore residents: findings from the 2016 Singapore mental health study. Sleep medicine: X. 2022;4:100043.

[46] Witek A, Lipowicz A. The impact of cigarette smoking on the quality of sleep in Polish men. Anthropological Review. 2021;84(4):369–382.

[47] Liu Y, Li H, Li G, Kang Y, Shi J, Kong T, et al. Active smoking, sleep quality and cerebrospinal fluid biomarkers of neuroinflammation. Brain, Behavior, and Immunity. 2020;89:623–627.

[48] Wang Q, Sundar IK, Lucas JH, Muthumalage T, Rahman I. Molecular clock REV-ERBα regulates cigarette smoke induced pulmonary inflammation and epithelial-mesenchymal transition. JCI Insight. 2021;6(12). doi:10.1172/jci.insight.145200.

[49] Hwang JW, Sundar IK, Yao H, Sellix MT, Rahman I. Circadian clock function is disrupted by environmental tobacco/cigarette smoke, leading to lung inflammation and injury via a SIRT1-BMAL1 pathway. The FASEB Journal. 2014;28(1):176.

[50] Feng L, Houck JR, Lohavanichbutr P, Chen C. Transcriptome analysis reveals differentially expressed lncRNAs between oral squamous cell carcinoma and healthy oral mucosa. Oncotarget. 2017;8(19):31521.

[51] Cadenas C, van de Sandt L, Edlund K, Lohr M, Hellwig B, Marchan R, et al. Loss of circadian clock gene expression is associated with tumor progression in breast cancer. Cell Cycle. 2014;13(20):3282Ð3291.

[52] Cornelissen G. Cosinor-based rhythmometry. Theoretical Biology and Medical Modelling. 2014;11(1):1–24.

[53] Veldman-Jones MH, Brant R, Rooney C, Geh C, Emery H, Harbron CG, et al. Evaluating Robustness and Sensitivity of the NanoString Technologies nCounter Platform to Enable Multiplexed Gene Expression Analysis of Clinical SamplesEvaluation of NanoString Technologies nCounter Platform. Cancer research. 2015;75(13):2587–2593.

