## Supplementary Information for "TimeTeller: a tool to probe the circadian clock as a multigene dynamical system"

### Supplementary Material for the paper "TimeTeller: a tool to probe the circadian clock as a multigene dynamical system"

[1]Mathematics Institute & Zeeman Institute for Systems Biology.
and Infectious Epidemiology Research, University of Warwick, Coventry CV4 7AL, UK.
[2]Odette Cancer Centre, Sunnybrook Health Sciences Centre, 2075 Bayview Ave.,
Toronto, ON, M4N 3M5, Canada.
[3]European Associated Laboratory "Personalizing Cancer Chronotherapy through Systems Medicine",
Institut National de la Santé et de la Recherche Médicale, UMRS 935,
Campus CNRS, 7 rue Guy Moquet, 9480- Villejuif, France.
[4]Assistance Publique-Hopitaux de Paris, Saint-Louis Hospital, Breast Disease Unit,
University Paris Diderot, 75475 Paris, France.
[5] Division of Biomedical Sciences, Warwick Medical School, University of Warwick, Coventry CV4 7AL, UK.

*To whom correspondence should be addressed;
.

[a]Current address: GSK Research, Gunnels Wood Road, Stevenage, Herts, SG1 2NY,UK
[b]Current address: CAMS Oxford Institute, Nuffield Department of Medicine, University of Oxford, Old Road Campus, Oxford, OX3 7BN
[c]Current address: Hit Discovery, Discovery Sciences, R&D, AstraZeneca, Cambridge, CB2 0AA, UK.

### Contents


S1 Data sets . . . S3
- S1.1 Mouse training datasets . . . S3
- S1.2 Mouse test datasets . . . S3
- S1.3 Baboon training dataset . . . S5
- S1.4 Human training dataset . . . S5
- S1.5 Human test datasets . . . S5
- S1.6 Preparatory RNAseq methods. . . . S6
- S1.7 Batch Effects . . . S6

S2 Probability Model & Likelihood curves $L_g(t)$ . . . S7
- S2.1 Probability model constructed from training data . . . S7
- S2.2 The likelihood curve $L_g(t)$ and the log threshold $l_{\text{thresh}}$ . . . S8
- S2.3 Choice of $l_{\text{thresh}}$ by inspecting the LRFs and the maximum likelihoods . . . S10
- S2.4 Parameters $\varepsilon$ and $\eta$ . . . S13

S3 Outline pseudocode of algorithm . . . S14

S4 Role of gene-to-gene correlations in the likelihood function . . . S14
- S4.1 $\Theta$ contains an estimate of a direct measure of clock precision . . . S14
- S4.2 $P(t|g)$ is estimated using $P(g|t)$ . . . S15
- S4.3 $\Theta$ depends crucially on the covariance structure of $P(g|t)$ . . . S15

S5 Does the rhythmic expression profile (REP) provide a faithful representation of clock dynamics? . . . S15

S6 TimeTeller normalisations . . . S16
- S6.1 Timecourse normalisation . . . S16
- S6.2 Intergene normalisation . . . S17
- S6.3 Timecourse then intergene normalisation . . . S17
- S6.4 Showing where the normalisations are used in the main paper . . . S17

S7 Singular Value Decomposition . . . S17
- S7.1 Optimal projections via SVD and the projection $U_{d,i}$ . . . S18

S8 Analysis of synchronicity and rhythmicity . . . S18
- S8.1 Zhang *et al.* microarray data . . . S19
- S8.2 Zhang *et al.* (2014) RNA-seq dataset . . . S20
- S8.3 Bjarnason *et al.* data . . . S21

S9 Extra information on the analysis of datasets . . . S22
- S9.1 Zhang *et al.* microarray data . . . S22
- S9.2 Testing Zhang *et al.* RNA-seq data when trained on Zhang *et al.* microarray data and vice-versa . . . S24
- S9.3 Bjarnason *et al.* data . . . S25
- S9.4 Kinouchi *et al.* liver data . . . S28
- S9.5 Le Martelot *et al.* data . . . S29
- S9.6 Hughes *et al.* data (*1*) . . . S30
- S9.7 Boyle *et al.* data (*2*) . . . S31
- S9.8 Feng *et al.* data (*3*) . . . S32

S9.9 Differentially expressed genes in the Boyle *et al.* data (*2*) and the Feng *et al.* data (*3*) . . . S33
S9.10 Acosta-Rodríguez *et al.* data . . . S34
S9.11 Assessing the coherence of downstream genes in the Acosta-Rodríguez *et al.* data . . . S37
S9.12 Mure *et al.* data . . . S44
S9.13 Analysing stopped clocks . . . S45
S10 Precision assessment without time stamps: Feng *et al.* and Cadanas *et al.* . . . S47
S11 Supplementary information about synthetic datasets: 1. Local PCs versus global PCs, 2. Θ dependence on the efficiency of a *Bmal1* knockdown . . . S48
S12 Stochastic modelling . . . S50
S12.1 The stochastic Relogio model . . . S50

Throughout these notes the main paper is referred to as **I**.

### S1 Data sets

#### S1.1 Mouse training datasets

| **Mouse training datasets** | |
|---|---|
| **Zhang et al. 2014 Microarray (*4*)** | |
| Technology | Microarray (Affymetrix MoGene 1.0 ST arrays) |
| GEO | GSE54652 |
| Tissue(s) | Adrenal, aorta, brown fat, heart, kidney, liver, lung, skeletal muscle, white fat |
| Entrainment conditions | 1 week 12 hr light/ 12 hr dark. Sample collection commenced after 18 hrs constant darkness. |
| Experimental conditions | Wild type. |
| Timepoints | CT18 - CT64, every 2 hrs. |
| Number of replicates | 1 sample per timepoint consisting of 3 pooled mice |
| Male or female? | Male |
| Age | 6 weeks |
| Strain | C57BL/6 |
| **Zhang et al. 2014 RNA-seq (*4*)** | |
| Technology | RNA-seq |
| GEO | GSE54652 |
| Tissue(s) | Adrenal, aorta, brown fat, heart, kidney, liver, lung, skeletal muscle, white fat |
| Entrainment conditions | 1 week 12 hr light/ 12 hr dark. |
| Experimental conditions | Sample collection commenced after 18 hrs constant darkness. Wild type |
| Timepoints | CT22 - CT64, every 6 hrs. |
| Number of replicates | 1 sample per timepoint consisting of 3 pooled mice |
| Male or female? | Male |
| Age | 6 weeks |
| Strain | C57BL/6 |

#### S1.2 Mouse test datasets

| **Mouse test datasets** | |
|---|---|
| **Kinouchi et al., 2018 RNA-seq** (*5*) | |
| Technology | RNA-seq |
| GEO | GSE107787 |
| Tissue(s) | Liver, skeletal muscle |
| Entrainment conditions | 12hrlight/12hrdark |
| Experimental conditions | Ad libitum fed vs 24 hr starved |
| Timepoints | ZT0 -ZT20, every 4 hrs. |
| Number of replicates | 3 samples per timepoint |
| Male or female? | Male |
| Age | 8 weeks |
| Strain | C57BL/6 |
| **Weger et al., 2021** (*6*) | |
| Technology | RNA-seq |
| GEO | GSE135898 |
| Tissue(s) | Liver |
| Entrainment conditions | 12hrlight/12hrdark |
| Experimental conditions | *Bmal* KO with WT controls and Cry1/2 double KO with WT |
| Timepoints | controls ZT0 - ZT20, every 4 hrs. |
| Number of replicates | 2 samples per timepoint |
| Male or female? | Male |
| Age | 12-16 weeks |
| Strain | C57BL/6 |
| **Yeung et al., 2018 (*4*)** | |
| Technology | RNA-seq |
| GEO | GSE100457 |

| Mouse test datasets (cont'd) | |
|---|---|
| Tissue(s) | Kidney |
| Entrainment conditions | 12 hr light / 12 hr dark |
| Experimental conditions | *Bmal* KO with WT controls. Night restricted feeding protocol for all mice. |
| Timepoints | ZT0 - ZT20, every 4 hrs. |
| Number of replicates | 2 samples per timepoint |
| Male or female? | Male |
| Age | 8-12 weeks |
| Strain | C57BL/6 |
| **Fang *et al.*** (*7*) | |
| Technology | Microarray (Affymetrix MoGene 1.0 ST arrays) |
| GEO | GSE59460 |
| Tissue(s) | Liver |
| Entrainment conditions | 12 hr light / 12 hr dark |
| Experimental conditions | 5 Wild Type and 5 *NR1D1* KO mice |
| Timepoints | ZT10 |
| Number of replicates | 1 |
| Male or female? | Male |
| Age | 8-12 weeks |
| Strain | C57BL/6 |
| **Barclay *et al.*** (*8*) | |
| Technology | Microarray (Affymetrix MoGene 1.0 ST arrays) |
| GEO | GSE33381 |
| Tissue(s) | Liver and adipose samples |
| Entrainment conditions | 12 hr light / 12 hr dark |
| Experimental conditions | Half of mice were were kept awake during ZT 0-6 on days 1 - 5 and days 8 - 12. Fed *ad libitum* |
| Timepoints | ZT1, ZT7, ZT13 and ZT19 |
| Number of replicates | 3 |
| Male or female? | Male |
| Age | 8-12 weeks |
| Strain | C57BL/6 |
| **LeMartelot *et al.*** (*9*) | |
| Technology | Microarray (Affymetrix MoGene 1.0 ST arrays) |
| GEO | GSE35789 |
| Tissue(s) | Liver, pooled from 5 mice |
| Entrainment conditions | 12 hr light / 12 hr dark |
| Experimental conditions | WT |
| Timepoints | ZT2, ZT6, ZT10, ZT14, ZT18, ZT22, ZT2(+24) |
| Number of replicates | 1 pooled from 5 mice |
| Male or female? | Male |
| Age | 8-12 weeks |
| **Acosta-Rodríguez *et al.*** (*10*) | |
| Technology | RNA-seq |
| GEO | GSE190939 |
| Tissue(s) | Liver |
| Entrainment conditions | Constant darkness |
| Experimental conditions | WT mice in 6 feeding conditions |
| Timepoints | 12 time points |
| Number of replicates | 2 (these are the observations at t and t+24) |
| Male or female? | Male |
| Age | 6 months & 19 months |
| Strain | C57BL/6J |
| **Koronowski *et al.*** (*11*) | |
| Technology | RNA-seq |
| GEO | GSE117134 |
| Tissue(s) | Liver |
| Entrainment conditions | 12hr light/ 12hr dark schedule |
| Experimental conditions | wild type (WT), *Arntl* knockout (KO), Liver-RE-Arntl-stop-FL (reconstructed) |
| Timepoints | ZT00, ZT04, ZT08, ZT12, ZT16, ZT20 |
| Number of replicates | 3 |
| Male or female? | Female |
| Age | 8-12 weeks |
| Strain | C57BL/6J |

#### S1.3 Baboon training dataset

| Baboon training dataset | |
|---|---|
| **Mure et al. 2014 RNA-seq (*12*)** | |
| Technology | RNA-seq |
| GEO | GSE98965 |
| Tissue(s) | Adrenal cortex, Adrenal medulla, Antrum, aorta, axillary, bladder, bone marrow, cecum, cornea, heart, ileum, kidney cortex, kidney medulla, liver, lungs, mesentric lymphonodes muscle abdominal, muscle gastrocnemius, oesophagus, omental fat, optic nerve head, pancreas, prostate, retina, retinal pigment epithelium, skin, smooth muscle, spleen, white adipose mesenteric, white adipose pericardial, white adipose perirenal, white adipose subcutaneous, and white adipose tissue. |
| Entrainment conditions | 12 hr light/ 12 hr dark. |
| Experimental conditions | Sample collection commenced after 18 hrs constant darkness. Wild type |
| Timepoints | ZT00 - ZT22, every 2 hrs. where ZT00 is time light is switched ON, ZT12 is time light is switched OFF (and no food intake) |
| Number of replicates | 1, all tissues from a baboon sampled per timepoint |
| Male or female? | Male |
| Age | young |
| Strain | C57BL/6 |

#### S1.4 Human training dataset

##### Human: Oral Mucosa Timecourse microarray data (Bjarnason *et al.* )

For this study Bjarnason *et al.* (*13*) recruited ten healthy human volunteers, five female and five male. Mucosa tissue was collected at six time points: 8 am, noon, 4 pm, 8 pm, 12 midnight, and 4 am. Subjects were selected after screening by clinical history, physical examination, routine blood work (complete blood count, electrolytes, creatinine) and actigraphy to confirm regular sleep-wake patterns. Mucosa samples were collected by a dental surgeon, using a tissue punch biopsy. Subjects went to sleep in a darkroom at their usual bedtime and were awoken for the midnight and the 4 am samples[a].

After collection, tissue mucosa samples were immediately frozen in liquid nitrogen and stored at $-80°C$ until use. Total RNA was prepared by Trizol Reagent (Invitrogen) in accordance with the manufacturer's specifications. RNA samples were quantified by optical density measurements at A260nm and A280nm. All samples were determined to be of high quality with A260:A280 ratios $>$ $1.9$. Total RNA (5$\mu$g) of each sample was used for microarray analysis on Affymetrix HG_U133_Plus2 chips. Cumulatively this chip represents 54,679 gene transcripts for analysis. Biotinylated cRNA was prepared according to the standard Affymetrix protocol (Expression Analysis Technical Manual, 2004, Affymetrix). Following fragmentation, 15 $\mu$g of cRNA were hybridized for 16 hrs at $45°C$ on GeneChip Human Genome U133 Plus 2.0 arrays. GeneChips were washed and stained in the Affymetrix Fluidics Station 450 and were scanned using the Affymetrix GeneChip Scanner 3000.

Although there were 16 probes identified in Note S8 as rhythmic and synchronised, we only use 15 probes going forward. The reason is that the *Per1* probe 244677_at was found to have significant signal issues in many of the independent human datasets, i.e. the signals values were very low. As there is another *Per1* probe in this dataset that does not have this problem, we can conclude that this is a probe issue, and not an issue with the *Per1* gene expression.

#### S1.5 Human test datasets

| Human test datasets |
|---|

[a]The Research Ethics Board at Sunnybrook Health Science Centre approved the clinical protocol for this study.

| Human test datasets (cont'd) | |
|---|---|
| **Boyle *et al.* (*2*)** | |
| Technology | GeneChip Human Genome U133 Plus 2.0 arrays |
| GEO | GSE59460 |
| Tissue(s) | Oral mucosa |
| Samples | 40 current smokers and 40 age and gender matched never-smokers underwent buccal biopsies. One smoker sample was excluded from the study based on a quality measure. |
| Study | Effects of smoking on the oral mucosal transcriptome |
| Timepoints | Unknown |
| Number of replicates | 1 |
| Gender | Mixed, gender matched |
| **Feng *et al.* (*3*)** | |
| Technology | GeneChip Human Genome U133 Plus 2.0 arrays |
| GEO | GSE59460 |
| Tissue(s) | Oral mucosa |
| Samples | 229 samples in total, 167 of OSCCs, 45 of normal oral mucosa, and 17 samples are of dysplasic oral mucosa tissue |
| Sudy | a comparative analysis of healthy oral mucosa transcriptome and oral squamous cell carcinoma (OSCC) transcriptome |
| Timepoints | Unknown |
| Number of replicates | 1 |
| Gender | Mixed |
| Note | Dysplasic tissue is abnormal tissue that could signify early signs of cancer. |
| **Cadenas *et al.* (*14*)** | |
| Technology | Affymetrix HG-U133A arrays |
| GEO | GSE11121, GSE2034, GSE6532 and GSE7390 |
| Tissue(s) | Breast cancer tumour |
| Samples | Mainz n=200, Rotterdam n=286, Transbig n=280. Total 766. |
| Study | a comparative analysis of healthy oral mucosa transcriptome and oral squamous cell carcinoma (OSCC) transcriptome |
| Timepoints | Unknown |
| Number of replicates | 1 |
| Note | Metadata, details and analysis is in Schmidt *et al.* . Cancer Res 2008; 68:5405-13; PMID:18593943. |

#### S1.6 Preparatory RNAseq methods.

All RNA-seq datasets were downloaded as .sra files from NCBIU's SRA database (accessed via the GEO database). SRA files were converted to fastq files using "fasterq-dump" from NCBIU's SRA Toolkit. FASTQ files were aligned to the mouse genome (GRC release m38.84) and converted to SAM files using HISAT2 v2.2.0 (Kim et al., 2019). SAM files were compressed to BAM using Samtools v1.10 (Heng Li et al., 2009). Transcript read counts were determined from the BAM files and the mouse transcriptome (GRCm38.84 .gtf file) using LiBiNorm v2.4, an in-house software package, in HTSeq-count mode (Anders et al., 2015; Dyer et al., 2019). Raw read counts were concatenated for all samples and exported as one text file for all subsequent analysis on a Mac OS. For dataset specific arguments see Table 4.2. Raw count normalisation was carried out in R Studio. The edgeR package was used to normalise the raw counts to log2 counts per million (logCPM) or log2 trimmed mean of M-values (logTMM). Data was also inspected for quality by checking library sizes, replicate plots and PCA plots.

#### S1.7 Batch Effects

In high-throughput studies batch effects occur because measurements are affected by variations in experimental conditions such as laboratory conditions, reagent lots, and personnel differences. We used the R packages BatchQC, ComBat and sva to analyse batch effects.

For the training data we are only concerned with the $G = 9 - 16$ selected clock-associated genes. We inspected the embedding of the data into $G$-dimensional space and the subsequent projections into

3-dimensional space using SVD for any sign of batch effects especially with respect to timing of the samples (see Figs. S1A and S35). Following analysis, the timing, $\Theta$ and ML values, and the likelihood ratio curves are inspected for signs of batch effects where this is relevant. A similar analysis is carried out after TimeTeller normalisation, a process that also reduces batch effects. We want to avoid batch correction as this would arbitrarily destroy the statistical distribution of the data (e.g., the covariance information) which is a crucial ingredient of our algorithm.

For the test data there is no question of a batch effect affecting the determination of $\Theta$ in a test data set as each sample is treated individually i.e. it is treated as an independent single sample in a way that does not depend upon any other test samples under both TimeTeller normalisation and the initial fRMA normalisation and then the probability model (which only depends upon the training data) is applied to the data from this sample independently. Our analysis also found no evidence of batch effects in this data when projected into $d = 3$-dimensional space using PCA and when analysed as above.

Using algorithms such as ComBat to remove batch effects between different tissue types has several disadvantages. Firstly, batches will typically not be evenly balanced as the training data will only contain WT/healthy data, whilst test data batches will contain both WT/healthy and perturbed data (e.g., genetically manipulated knock-out (KO) samples). The inappropriate application of batch correction methods to datasets with unbalanced batches is reviewed elsewhere (*15*). A second disadvantage to the use of batch correction is that it necessitates that the model be re-trained for every new training and test dataset combination post-batch correction, therefore it is difficult to directly compare findings from different test datasets. Ultimately, one of the primary aims is to apply TimeTeller to human biopsies - both healthy and unhealthy. Batch correction of human data would require *a priori* knowledge of covariates, which may not even be known. In other words, 'real world' data contains many variables which would confound batch correction such as patient age, sex, time of sampling, health status etc. Therefore, a method for time prediction should ideally be applicable without the need for batch correction.

#### S2 Probability Model & Likelihood curves $L_g(t)$

An outline pseudocode description of the algorithm is in Note S3.

##### S2.1 Probability model constructed from training data

For a given training dataset we firstly choose the panel of rhythmic genes that TimeTeller will use. This is called the *rhythmic expression profile* (REP) and $G$ denotes the number of genes in the REP. For a given transcriptomics sample the expression levels $g_k$, $k = 1, \ldots, G$, of these genes are collected into a vector $(g_1, \ldots, g_G)$ which we will call the *rhythmic expression vector* (REV). The training data will have been collected at sample times $t_i$, $i = 1, \ldots, N_t$. In the training data used here the number $N_s$ of samples at each time point is the same. Therefore, the $G$-dimensional REVs with sample time $t_i$ can be labelled by the $i$ and $j$ and denoted $\bar{g}_{ij}$.

As discussed in **I** and below, we use four training datasets. In three of these the $j$s correspond to different tissues and in the other to different individuals. We call the $j$s *instances*. Each $g_{ij}$ is then normalised using timecourse and/or intergene normalisation as described in **I** and Sect. S6 resulting in vectors $g_{ij}^{\mathrm{norm}}$. As explained in Sect. S8 the REP is chosen so as to try and minimise the effect of the data coming from different instances and the normalisation is used to remove what is left. These $g_{ij}^{\mathrm{norm}}$ are the vectors that will be used to train TimeTeller. The issue of batch effects is considered in Sect. S1.7.

To construct the probability model we firstly construct one for each timepoint $t_i$ in the training data by using the local statistical structure of the data at that timepoint and then we combine these. Associated with this time $t_i$ is the set of $N_s$ points $g_{ij}^{\mathrm{norm}}$, $j = 1, \ldots, N_s$, in $G$-dimensional space. The projection operator described in Sect. S7.1 gives a way to linearly project these points into $d$-dimensional space for all $d < G$ with well-known optimality properties (see Sect. S7). This produces a corresponding set $\mathrm{Q}_{t_i}$ of $N_s$ $d$-dimensional points $Q_{ij}$, $j = 1, \ldots, N$. We then fit a multivariate normal distribution (MVN) to the points $Q_{ij}$. The dimensionality $d$ is chosen so that there are enough vectors $Q_{ij}$ to fit a $d$-dimensional multivariate Gaussian (using the MATLAB function fitgmdist) while ensuring that most of the variance in the data is captured by the $d$-dimensional projection (e.g. see Fig. S2). In our case we take $d = 3$. A MVN distribution is defined by its $d$-dimensional mean and $d \times d$ covariance matrix which we denote by $\mu_i(t_i)$ and $\Sigma_i(t_i)$ respectively.

We fit a periodical piecewise cubic hermite interpolating polynomial spline through the mean vectors $\mu_i(t_i)$ and each of the six entries that determine the $3 \times 3$ symmetric matrix $\Sigma_i(t_i)$ so as to extend $\mu_i(t_i)$ and $\Sigma_i(t_i)$ to all times $t$ between the time points, thus obtaining $\mu_i(t)$ and $\Sigma_i(t)$. For this we use the MATLAB function *perpchip*. Our software offers some alternatives to *perpchip* such as the MATLAB function *pchip* which is shape preserving, i.e. continuity of the second derivative is not obligatory. With the latter, for example, if two covariance matrix entries were identical for two consecutive time Gaussians, the Hermite spline allows the value of the joining spline to stay the same in the space between, while a standard spline would enforce some change. For consistency across all data says discussed in **I** and this SI we only use perpchip. Using this approach, for each value of the time index $i$, we determine a family of $d$-dimensional MVN distributions $\mathcal{P}_{i,t}$ for all times $t$ between the first and last data times. These have mean $\mu_i(t)$ and covariance $\Sigma_i(t)$. This family of MVN distributions indexed by time is what we refer to as the *probability model*.

#### S2.2 The likelihood curve $L_g(t)$ and the log threshold $l_{\text{thresh}}$

Now we define the likelihood curve $L_g(t)$ where $g$ is a REV from either training or test data. We assume that we have calculated the probability model.

For each of the time indices $i$ we define the likelihood curve associated with the $i$th timepoint using the probability given by the MVNs $\mathcal{P}_{i,t}$ i.e.

$$L_{g,i}(t) = \frac{1}{(2\pi)^{d/2}|\Sigma_i(t)|^{1/2}} \exp\left(-\frac{1}{2}(U_{d,i}g^{\text{norm}} - \mu_i(t))^T \Sigma_i(t)^{-1}(U_{d,i}g^{\text{norm}} - \mu_i(t))\right)$$

where $g^{\text{norm}}$ is the vector obtained after normalising $g$ with the relevant normalisation (see Sect. S6). The idea is to obtain $\log L_g(t)$ by averaging these individual log likelihoods $\log L_{g,i}$, $i = 1, \ldots, s$ but some modification is needed. To ensure that this product is not wrecked by inaccurate exceptionally low values of one $L_{g,i}$ affecting robust high values of another at the same $t$ we truncate the $L_{g,i}$. A curve $L_{g,i}$ will take on very low values away from its maximum and these may well be inaccurate. If this happens at a $t$ value for which another such curve $L_{g,j}$ has a high accurate value then this may badly affect the estimate of $L_g(t)$. To overcome this we fix a lower threshold $l_{\text{thresh}} < 0$ and replace each $\log L_{g,i}$ by $\max\{\log L_{g,i}, l_{\text{thresh}}\}$ in the sum so that $L_g(t)$ is defined by

$$\log L_g(t) = \frac{1}{s}\sum_{i=1}^{s} \max\{\log L_{g,i}, l_{\text{thresh}}\}.$$

The way to choose the value of $l_{\text{thresh}}$ is discussed below in Sect. S2.3.

A common characteristic of likelihood curves corresponding to poor clock function is the emergence of a second peak often roughly 8-12h from the dominant peak as shown in many of the centered likelihood ratio curves especially for those test samples showing dysfunction. Often this is because the curve in $d$-dimensional space tracing out the means of $P(g|t)$ (as in our visualisations) has a (distorted) elliptical shapes. Because of this shape we often have the situation where training data points at CT$t$ are close to those of CT$(t + t_1)$ with $t_1 \approx 8h - 16h$ and this can give rise to two peaks in the likelihood curve $L_g(t)$. Another way this can affect our analysis is shown in Fig. 2H of **I**. We see there that if we plot the real times against estimated times for those samples in the Feng *et al.* cancer group many have unreasonable timing $T$ which becomes reasonable when the second peak is instead chosen to give $T$.

If there is such a second peak then we want to penalise it if it has a height whose ratio to the height of the primary peak is above a threshold that depends upon the time $\delta t$ between these two peaks.

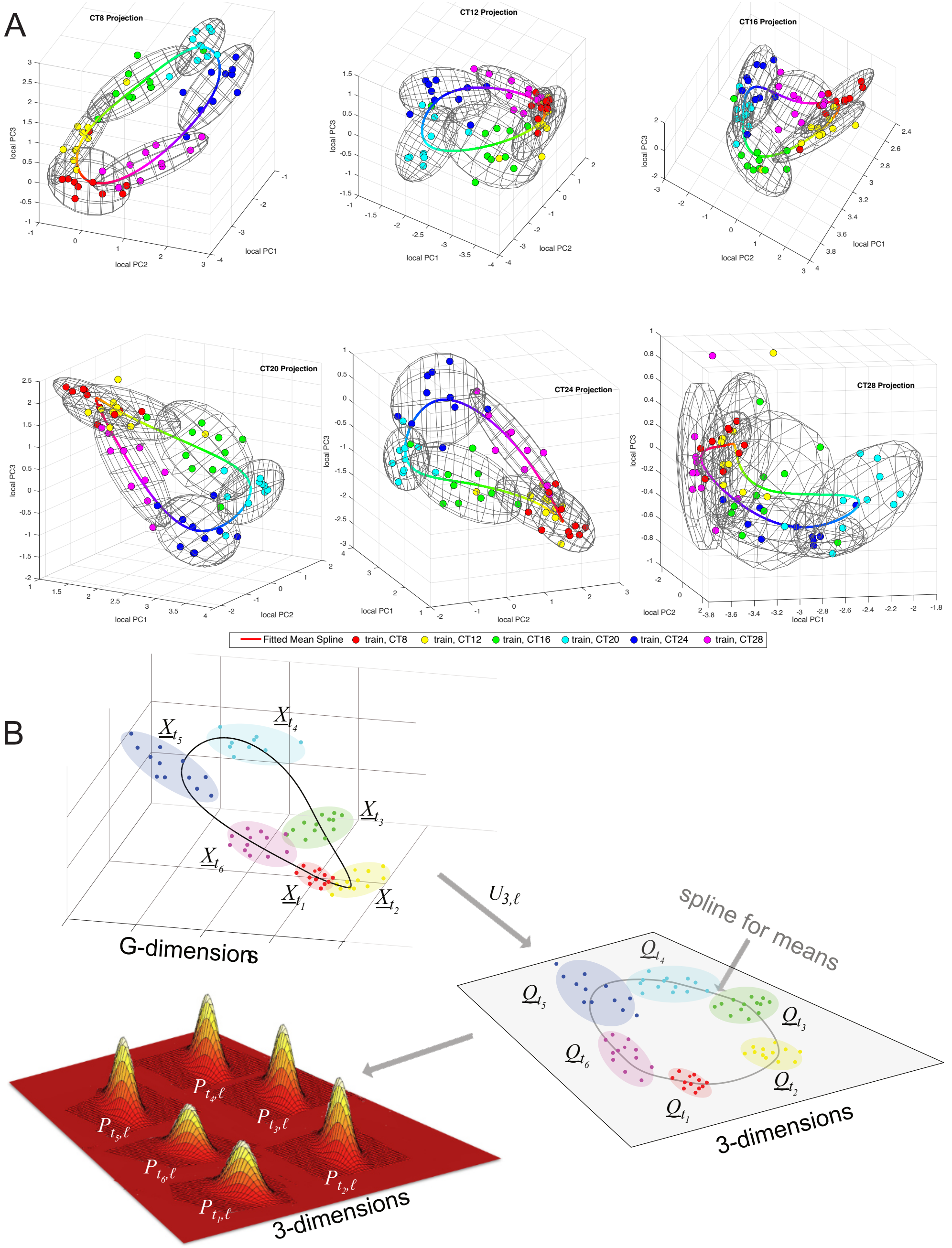


**Figure S1: A. All 6 local projected spaces for human data.** The data is projected as described in Material and Methods using the projections $U_{d,i}$ where $d = 3$ and $i$ indexes the six times. Each time point is coloured according to its time. Splines through the means $\mu_i(t_j)$ of each set of 10 time data points show (distorted) elliptical shapes. Because of this shape we often have the situation where training data points at CT$t$ are close to those of CT$(t + 12)$ and this can give rise to two peaks in the likelihood curve $L_g(t)$ if the data point corresponding to $g$ lies between the training data points at CT$t$ and those of CT$(t + 12)$. **B. This schematic outlines the construction of the likelihoods** $L_{g,i}(t)$. For each $t_i$ the set of vectors $X_{t_i} = \{g_{ij}, j = 1\ldots, N_s\}$ are projected into $d = 3$ dimensions using $U_{d,i}$ to get $\mathrm{Q}_{t_i} = \{Q_{ij}, j = 1\ldots, N\}$. A MVN distribution is estimated for each $\mathrm{Q}_{t_i}$ and then these distributions are interpolated using splines to all times $t$ of the day. The projections $U_{,i}$ for the Bjarnason *et al.* data and $d = 3$ is shown above in A.

Moreover, this threshold should be minimal when $\delta t = 12$h. To do this we use the curve $C(t|T)$ below.

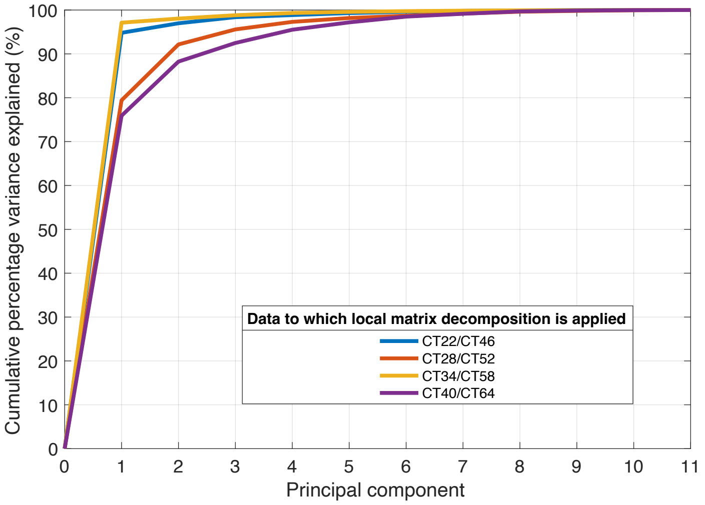


**Figure S2:** Cumulative percentage of variance explained by each of the principal components found by SVD for the Zhang *et al.* data. Data for day 1 and day 2 (e.g., CT22 and CT46) is combined to build a 24 hr model. 3 dimensions are sufficient to explain more than 90% of the variation in the dataset for each local projection. Such a situation follows when the eigenvalues of the covariance matrices of the $P(g|t_i)$ decay rapidly.

The *clock dysfunction metric* $\Theta$ is defined to be the proportion of time $t$ which satisfies

$$\Lambda(Z,t) = \frac{\mathcal{L}(Z|t)}{\mathcal{L}(Z|T))} \geq C(t|T) \tag{1}$$

i. e. the proportion of time in the day when the given likelihood ratio curve is above the curve $C(t|T)$. Here $C(t|T)$ is defined by

$$C(t|T) = \eta\left(1 + \varepsilon + cos\left(\frac{t-T}{24}2\pi\right)\right) \tag{2}$$

e.g., the black curve in Fig. S3. This is a simple cosine curve transformed so that

$$\varepsilon \leq C(t|T)/\eta \leq 2+\varepsilon, \quad C(T|T) = (2+\varepsilon)\eta \quad \text{and} \quad C(T+12|T)) = \varepsilon\eta.$$

The choice of the parameters $\varepsilon$ and $\eta$ is discussed below in Sect. S2.4.

#### S2.3 Choice of $l_{\text{thresh}}$ by inspecting the LRFs and the maximum likelihoods

We discuss how the cut-off threshold $l_{\text{thresh}}$ is chosen using the example of the two Kinouchi *et al.* datasets, one for skeletal muscle and the other for and liver.

The key considerations underlying the choice of $l_{\text{thresh}}$ are that it should be as large as possible subject to the conditions that

(i) very few training and control samples have flat regions that significantly intersect $C(t|T)$ and thus contributing to $\Theta$, and

(ii) as many as possible of the test data samples should have MLs above $exp(l_{\text{thresh}})$.

There are two reasons we do not want $l_{\text{thresh}}$ to be decreased further than necessary. Firstly, because as we decrease $l_{\text{thresh}}$ further the LC starts to depend on very small values in the $L_{g,i}$ that may well be inaccurate (see Sect. S2.2). Secondly, because if $l_{\text{thresh}}$ is reduced too far you are likely to remove structure in the LRF that is at times that are away from the time $T$ of maximum likelihood. This happens because, if the $L_{g,i}$ have their maxima not too far from $T$ then decreasing $l_{\text{thresh}}$ causes a much bigger decrease in the likelihood $L_g(t)$ away from $T$ than near $T$ and therefore decreases

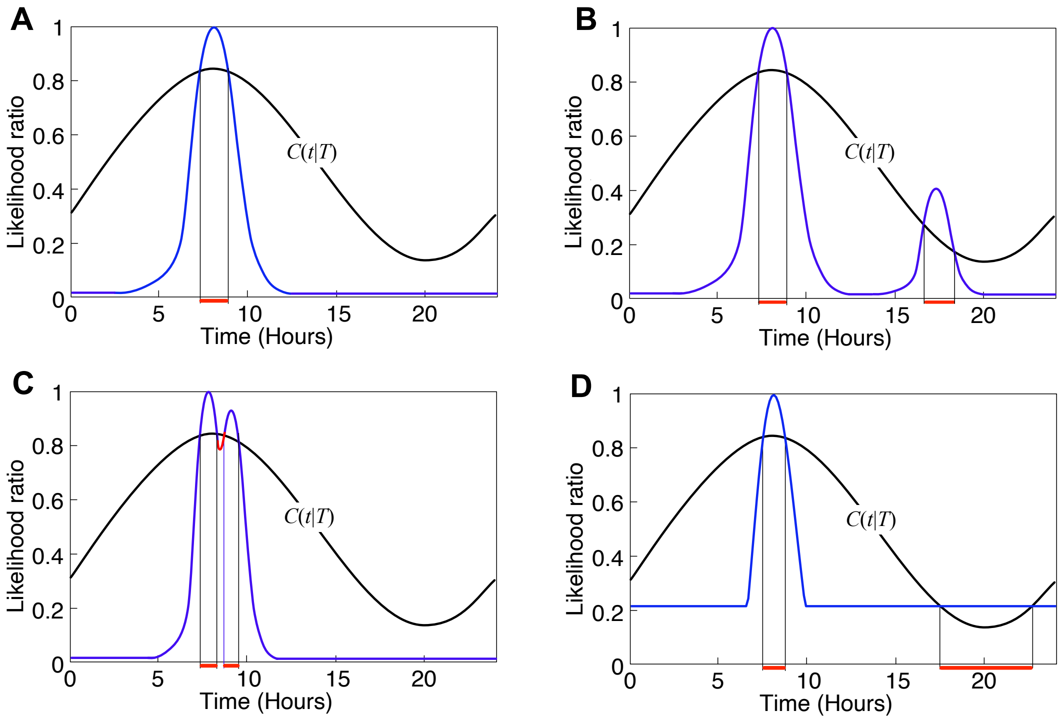


**Figure S3: A-C. Showing how the modification $C(t|T)$ affects the definition of $\Theta$.** See Sect. S4.1 for a further discussion. The graph of $C(t|T)$ is shown in black. Many, LRFs contain a combination of the various ways in which the widths of the peaks and the flat regions discussed in **A-D** contribute to increasing $\Theta$. **A.** This shows how a single peak in a likelihood curve would not be affected by the penalised cut-off. **B.** This shows how if there is a secondary peak approximately 8-12h from the MLE, then if the second peak rises above $C(t|T)$ the $\Theta$ will be penalised. **C.** This shows how if there is a secondary peak near to the MLE peak, then $\Theta$ will also be penalised. textbfD. This shows how a flat region in the LRC can make a contributing to $\Theta$ if it intersects $C(t|T)$. Such an intersection occurs if ML is not much greater than $\exp(l_{\text{thresh}})$, precisely if ML is smaller than $\alpha\beta\exp(l_{\text{thresh}})$ ($\alpha$ & $\beta$ as in the definition of $C(t|T)$). This is the way a low ML makes a contribution increasing $\Theta$. As one can see from the graph decreasing the ML increases the contribution from this flat region. The values of $\Theta$ in these plots is $l/24$ where $l$ is the sum of the lengths in hours of the red segments at the base of the graph.

the LRF away from $T$ while maintaining the peak structure near $T$. If the dysfunction is mainly manifested by low ML then decreasing $l_{\text{thresh}}$ by too much moves all the flat regions in the LRFs down below $C(t|T)$ while if it manifested by structures such as second peaks the these are also decreased below $C(t|T)$. In both cases this results in a decrease in $\Theta$. This phenomenon occurs clearly in the Kinouchi *et al.* liver data discussed below when the value of $l_{\text{thresh}}$ is taken too low (see Figs. S16 and S17).

If the maximum value $\log \text{ML}$ of $\log L_g(t)$ is only just above $l_{\text{thresh}}$, then it and the corresponding likelihood ratio curve will have intervals on which they are flat. If this is the case then the length of these flat intervals above the minimum of the curve $C(t|T)$ defined in Note S2 can contribute to $\Theta$. This contribution has interesting information in it because it is related to how low the maximum value ML of $L_g(t)$ is.

If the criterion (ii) results in too small a value so that too much structure has been removed, it is then generally acceptable to set $l_{\text{thresh}}$ at a higher value provided that the number of training samples violating (i) does not get too large. It so also desireable that the the number of test samples violating (ii) is not too large as otherwise many samples have $\Theta = 1$ meaninf that these samples do not have a non-trivial stratification even though they will be distinguished as having higher dysfunction than other samples.

Fig. S4 concerns the Kinouchi *et al.* skeletal muscle data. The first step is to get an estimate of the

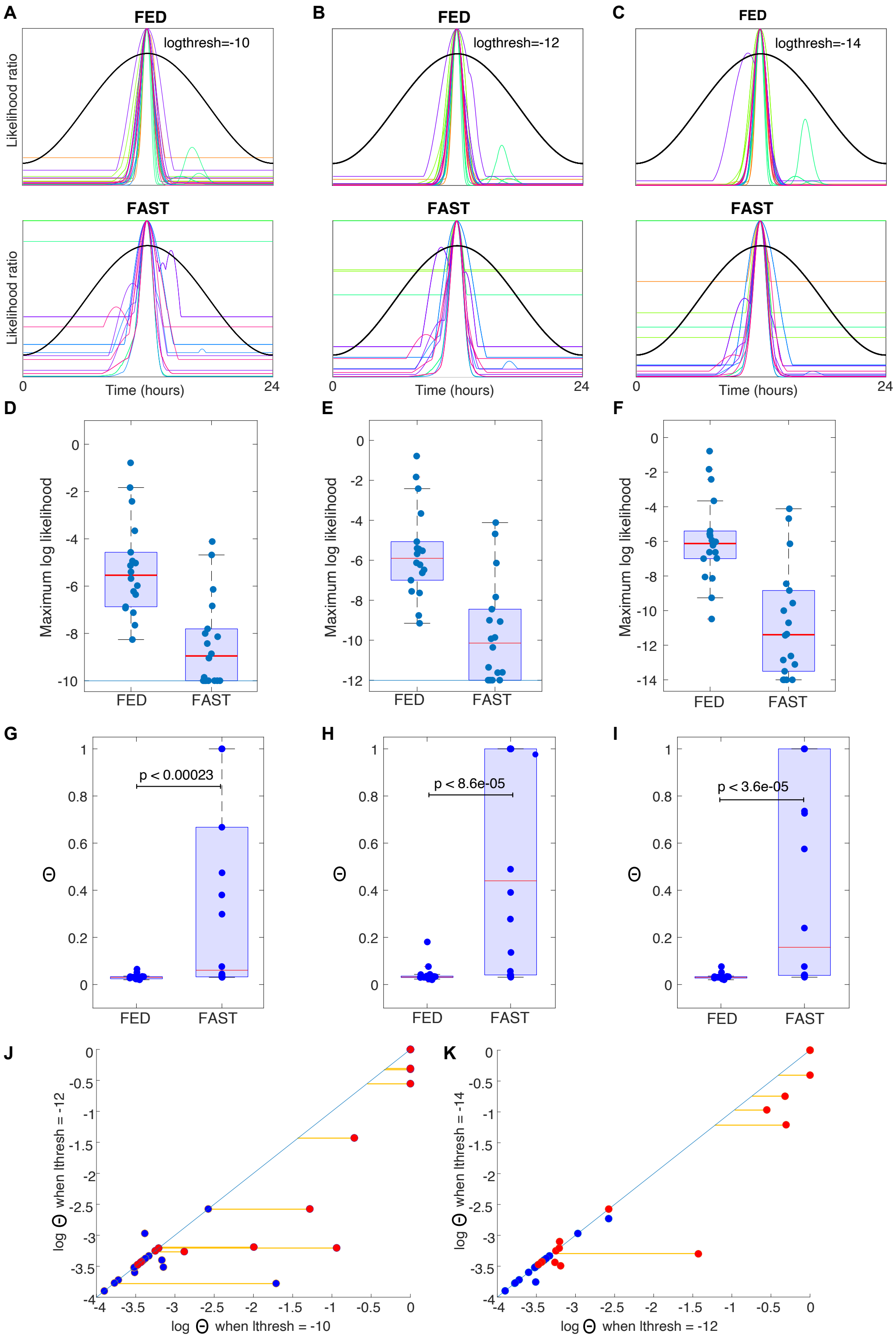


**Figure S4: Choosing $l_{\text{thresh}}$.** See Sect. S2.3 for more details. The data shown is for the Kinouchi *et al.* skeletal muscle data. **A-C.** Centre LRFs for $l_{\text{thresh}} = -10, -12$ and $-14$. **D-I.** Box plots of the MLs and $\Theta$s for the three values of $l_{\text{thresh}}$. In D-F the blue horizontal line shows the value of the threshold $\exp(l_{\text{thresh}})$. p-values shown are from the Wilcoxon rank sum test (Matlab function ranksum). **J.** Scatter plot showing how $\Theta$ and the ordering of $\Theta$s change when $l_{\text{thresh}}$ is decreased. Blue points FED, red points FAST. Orange lines guide the eye when checking whether a sample changes its position in the $\Theta$ stratification. Just one point does this in K.

the values of ML. From Fig. S4D,E,F we see the median of these for FAST samples from this data is a bit below $10^{-5}$ which is approximately $\exp(-11.5)$ which suggests a value of -10 or -12 for $l_{\text{thresh}}$.

The three columns in Fig. S4 correspond to respectively $l_{\text{thresh}} = -10$, $-12$ and $-14$. The setting $l_{\text{thresh}} = -10$ is seemingly an acceptable value because then only one of the FED control data samples has a flat region that intersects $C(t|T)$. However, we could also choose $l_{\text{thresh}} = -12$ because then none of the FED samples intersects the curve $C(t|T)$ and this setting is what we have used in the main paper. D, E and F in Fig. S4 show the corresponding maximum likelihood values.

The MLs hardly change with changing $l_{\text{thresh}}$ the key difference just concerns the change in the number of samples with their ML value at the value of the threshold $\exp(l_{\text{thresh}})$. This change occurs because there are FAST samples where ML is below the cut-off threshold and the number of these below-threshold samples decreases as we decrease $l_{\text{thresh}}$.

The $\Theta$ values that are obtained with these thresholds are shown in Fig. S4G,H,I. The way that the values change is shown in figure Fig. S4J,K. We see that when we change from -12 to -14 there is hardly any change in the ordering of the $\Theta$ values and hence the stratification, and this further justifies a choice of $l_{\text{thresh}} = -12$.

This choice is for the skeletal data. However, when we analyse the Kinouchi *et al.* liver data using $l_{\text{thresh}} = -12$ we obtain the results shown in Fig. S17. These suggest that the substantially worse timing for the FAST data is due to type $lowML$ dysfunction. However, all the structure in the FAST LRFs except for the central peak has been removed as discussed above and consequently the observed $\Theta$ values do not pick this $lowML$ dysfunction up. A quick inspection of the MLs suggests that $l_{\text{thresh}} = -12$ gives too low a threshold because only two sample have a ML below $\exp(-5.5)$ and these are much lower at approximately $\exp(-10)$. This suggests using a threshold with $l_{\text{thresh}} = -6$ or $-7$. In Fig. S16 we used $l_{\text{thresh}} = -6$ and we see that then the $\Theta$ spectrum nicely picks up the $lowML$ dysfunction. One might ask why we did not just use the ML to stratify things in this case and, indeed, this would give essentially the same answer for this data. However, this is not true when our samples have significant structures like second peaks as, for example seen in the Fang *et al.* , Boyle *et al.* and Feng *et al.* datasets and in other cancer datasets that we have studied. for these $\Theta$ integrates the different types of dysfunction.

#### S2.4 Parameters $\varepsilon$ and $\eta$

The values of $\varepsilon$ and $\eta$ used in the definition of $\Theta$ should be appropriately chosen. We must choose $\varepsilon > 0$ so that $C > 0$. The larger $\varepsilon$ is, the less anti-phase peaks impact the final confidence metric. Once chosen, where possible the values used should be maintained in order to enable comparisons of $\Theta$ values across datasets.

From (1), it is clear that to ensure the positivity of $\Theta$ we must enforce that

$$C(t|T) < 1 \tag{3}$$

As $\max(C) = \eta(2+\varepsilon)$ and $\min(C) = \eta\varepsilon$, we must have,

$$0 < \eta\varepsilon < \eta(2+\varepsilon) < 1 \tag{4}$$

and, note that the last part of this implies $0 < \eta < 1/(2+\varepsilon) < 0.5$.

Also note that $\eta\varepsilon$ defines the minimum value of the threshold for which the graph of the likelihood ratio $\Lambda$ can be intersected, and $\eta(2+\varepsilon)$ defines the maximum. We set $\varepsilon = 0.4$ and $\eta = 0.35$, which sets the maximum threshold value (at the MLE) to $\eta(2+\varepsilon) = 0.84$, and the minimum threshold (anti-phase to the MLE) to $\eta\varepsilon = 0.14$. This means that if there is a peak in the likelihood exactly anti-phase to the MLE that is more than 14% of the maximum, then it penalises $\Theta$.

The hyperparameters $\varepsilon = 0.4$ and $\eta = 0.35$ are used throughout. However, we note that, since the likelihood curves are continuous (and smooth away from the ends of flat regions) the same is true of the dependence of $\Theta$ upon the parameters $\eta$ and $\varepsilon$. Consequently, the precise choice of $\varepsilon$ and $\eta$ is not crucial.

As $\varepsilon \to \infty$ the method is the same as a non penalised approach and as $\varepsilon \to 0$ any anti-phase secondary peak above $m_t$ will penalise $\Theta$.

In Fig. S3 we show a selection of centered likelihood ratio curves for the indicated data sets together with the graph of $C(t|T)$ in black. In each case the likelihood ratio curve has been centered so that the maximum is at $t = 12$. This enables easier comparison of the curves and the associated $\Theta$ value.

### S3 Outline pseudocode of algorithm

**Algorithm 1:** TimeTeller

Training data consists of a set $G_{t_i}$ of $j = 1, \dots, N$ gene expression vectors $g_{ij}$ at s timepoints $t_i$

Normalise gene expression vectors $g_{ij}$ using timecourse or intergene normalisation.

**for** *projection time index* $p = 1$ **to** $s$ **do**

  calculate SVD projector $U_p$ using $G_{t_p}$ (Note S7);

  **foreach** $i \in \{1, 2, \dots, s\}$ **do**

    use $U_p$ to project $G_{t_i}$ into $\mathbb{R}^d$ to get projected data $G_{t_i}^p$

    fit multivariate normal to projected $G_{t_i}^p$, gives mean $\mu_p(t_i)$ and covariance matrix$\Sigma_p(t_i)$

    fit splines to extend $\mu_p$ and $\Sigma_p$ from $t_i$ to all times of day

Test data consists of a gene expression vector $g$ usually using the same expression technology as the training data.

**for** *projection time index* $p = 1$ **to** $s$ **do**

  Calculate $\ell_p(t) = \log P(g|t)$ for the MVN distributions $P(g|t)$ given by $\mu_p(t)$ and $\Sigma_p(t)$.

Combine these to form $\ell(t) = N^{-1} \sum \min(\ell_p(t), \tau)$ after applying a threshold $\tau$ to $\ell_p(t)$.

Likelihood function is $L(t) = \exp \ell(t)$ and likelihood ratio function (LRF) is $\Lambda(t) = L(t)/L(T)$ where $T$ is the time that $L(t)$ is maximal.

Use $\Lambda(t)$ to calculate $\Theta$ (Note S2).

### S4 Role of gene-to-gene correlations in the likelihood function

#### S4.1 $\Theta$ contains an estimate of a direct measure of clock precision

In what follows $g = (g_1, \dots, g_G)$ denotes a vector consisting of the possibly normalised levels of $G$ clock-associated genes. We call these *normalised rhythmic expression vectors* (NREVs).

We consider the probability $P(t, g)$ that $g$ is observed at time $t$ in WT conditions and the associated conditional probability distributions $P(g|t)$ and $P(t|g)$. Then if we want to assess how well the clock is working in an independent sample with expression vector $g^* \in \mathbb{R}^G$ we would want to estimate the conditional distribution $P(t|g^*)$. The time $T$ would be estimated to be the value of $t$ maximising $P(t|g^*)$ i.e. the maximum likelihood estimate (MLE).

The corresponding likelihood ratio is given by

$$\Lambda(t) = P(t|g^*)/P(T|g^*).$$

As is well-known (e.g., Chaps. 8 & 9 of (*16*)) the likelihood ratio confidence interval gives a good estimate of the confidence intervals for the MLE when the log likelihood function is approximately quadratic on some scale as is the case with our likelihood functions. If $\alpha$ is the required sensitivity (i.e. 1 minus the confidence level) then the confidence interval is given by the set of times $t$ that satisfy

$$\Lambda(t) \geq \exp -\chi^2_{\alpha,1}/2.$$

We can relate this to $\Theta$ (see Note S2) when the likelihood curve only has a single peak (or, more generally, only twice meets the curve $C$ in Note S2). Since the likelihood ratio curve is quadratic near its maximum we deduce that the contribution to $\Theta$ from the region around the maximum is

$$\Theta = \sqrt{2\eta/\chi^2_{\alpha,1}}C$$

where $C$ is the length of the confidence interval and $\eta$ is the parameter in Note S2.4. Since the confidence intervals define the precision of the clock we see that, in this case, $\Theta$ is a direct measure of clock precision.

##### S4.2 $P(t|g)$ is estimated using $P(g|t)$

To measure $P(t|g)$ we will use the fact that, by Bayes theorem, so far as dependence upon $t$ is concerned,

$$P(t|g) = \frac{P(g|T)P(t)}{P(g)} \propto P(g|t)$$

provided we assume that $P(t)$ is reasonably independent of $t$. Therefore, we can use $P(g|t)/P(g|T)$ to estimate $\Lambda(t) = P(t|g)/P(T|g)$. Consequently, TimeTeller tries to estimate $P(g|t)$ directly from WT data. For the estimation it is assumed that $P(g|t)$ is multivariate normal.

##### S4.3 $\Theta$ depends crucially on the covariance structure of $P(g|t)$

To see what $\Theta$ depends upon consider the case where $P(t|g^*)$ is relatively sharply peaked around $T$. Expanding around this estimate, the distribution is approximately Gaussian with

$$P(t|g^*) \approx \frac{1}{\sqrt{2\pi\sigma_t}} \exp\left[\frac{(t-T)^2}{2\sigma_t^2}\right]$$

and the variance $\sigma_t$ is given by

$$\frac{1}{\sigma_t^2} = \delta g^T \cdot \Sigma^{-1} \cdot \delta g$$

where $\Sigma$ is the covariance matrix of $P(g|t)$ and $\delta g$ is the derivative with respect to $t$ of the mean of $P(g|t)$ at $g = g^*, t = T$. Indeed, if we drop the tightness assumption we can use the Cramer-Rao theorem to deduce that the term on the righthand side is a lower bound for the variance because this term is the dominant part of the Fisher Information matrix of $P(g|t)$ with respect to $t$.

Therefore, we see that it is crucial in estimating our clock dysfunction metric that we take account of the covariance structure of $P(g|t)$.

#### S5 Does the rhythmic expression profile (REP) provide a faithful representation of clock dynamics?

It is necessary to try and ensure that the choice of the the REP and rhythmic properties of the genes included provide a faithful representation of the clock state i.e. so that the mapping from clock state to NREV $(g_1, \ldots, g_n)$ is an embedding. This means that the NREV is a smooth function of the clock state and that each state of the NREV arises from at most one clock state. Conditions for this to be the case are discussed in the area of applied dynamical systems known as *embedology* (*17–19*). This addresses the situation where one has a dynamical system whose state depends upon many variables $y_1, \ldots, y_N$ and one seeks a faithful representation of the dynamics in terms of a much reduced number of variables $x_1, \ldots, x_n$ and shows that for oscillating non-chaotic systems like ours it is reasonable to find such a representation by the sort of projections we use. In our case the $y_i$ represent the expression levels of all the molecular components of the GRN underlying the clock and the $x_i$ are the expression levels of the genes in the REP. There is no way to prove that such a REP does fully represent the dynamics but for a tightly coupled dynamical system like the circadian clock, one can validate this to a great extent by checking that increasing the size of the REP does not alter results and, more specifically, checking the singular values of the local projections introduced in Sect. S2 as in Fig. S2 to ensure that the dynamics of the $n$-dimensional REP can be reduced to an even smaller dimension $d$ as explained in Sect. S2. We deal with it by the use of a REP with significantly more genes that are likely to be needed. Note that we are not suggesting that the genes in the REP are the only genes regulating the clock but are hypothesising that they are enough to provide a full representation of the clock dynamics for the data sets considered.

### S6 TimeTeller normalisations

See Table S5.

#### S6.1 Timecourse normalisation

We observed that the timeseries for clock genes in RNA-seq data tended to have much greater tissue to tissue variation than that when using the Affymetrix MoGene 1.0 ST and GeneChip Human Genome U133 Plus 2.0 microarray platforms. This is illustrated for example in Fig. S5 below.

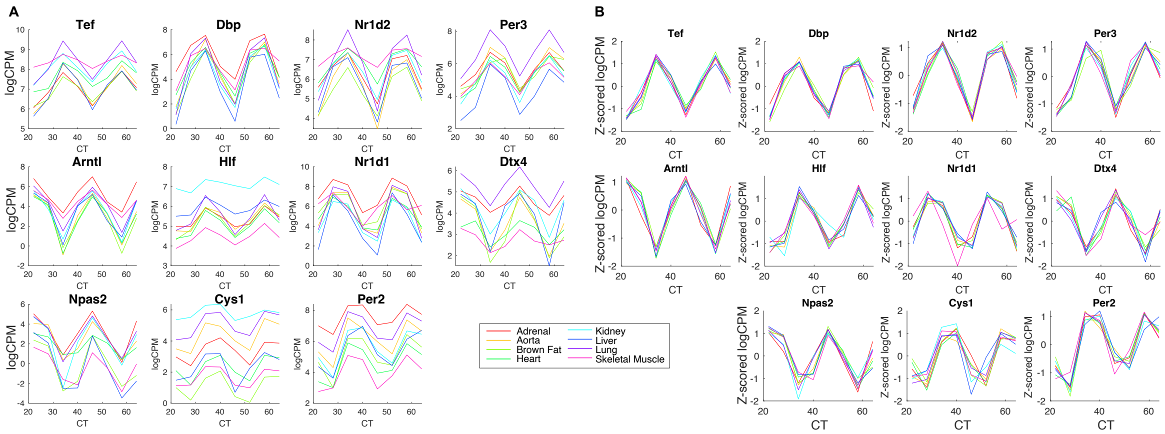


**Figure S5:** Time-dependent expression of the RNA-seq transcripts in the Zhang *et al.* RNA-seq training data. **A.** Before timecourse normalisation different tissues express the oscillatory genes at quite different amplitudes and magnitudes. **B.** After normalisation amplitudes and magnitudes are better aligned.

In *timecourse normalisation*, for each gene we normalise its timeseries $g_{ij}(t_k)$ by replacing it by $(g_{ij}(t_k) - \bar{g}_{ij})/\sigma_{ij}$ where $\bar{g}_{ij}$ is the mean of the expression levels $g_{ij}(t_k)$ over the time points $t_k$ and $\sigma_{ij}^2$ is their variance. Fig. S5 shows the result of this for the Zhang *et al.* RNA-seq data.

When doing this it is necessary to define what timeseries are to be used and choose these so that each sample in the training data is in exactly one timeseries. For example if at each timepoint each sample has two replicates and there is no distinction between the replicates then for one timeseries one can take replicate 1 and for another replicate 2, but there are many other possible choices such as alternating replicate 1 and 2 in each of the two timeseries. In all the cases we consider there is a natural way to do this allocation. In cases like the one with two replicates there is also a possible choice of normalising the timeseries separately or jointly. We always choose the separate approach.

When timecourse normalisation has been used on the training data, it is necessary to normalise any test data. We use *timecourse-matched* normalisation where, if the test data is $g_{ij}(t_k)$ (perhaps with unknown $t_k$), then we use the measured values of $\bar{g}_{ij}$ and $\sigma_{ij}^2$ from the training data normalisation to normalise it. This can only be done if the test data tissue is one of the ones in the training data. If not there are two possible ways to proceed. If the test data consists of timeseries then it can be normalised by timecourse normalisation. If not then the only way to proceed is via intergene normalisation for both training and test data. Similar considerations apply when the instances are individuals or conditions.

When using timecourse-matched normalisation, it is also crucial that the training data are produced by the same transcriptomics platform (e.g., RNA-seq). However, if you have appropriate training data using one platform and also a single training timeseries and test data using another platform then, even though there is not enough timeseries data in the second platform to estimate a training clock, one can assess the test data using the trained clock by timecourse normalising the single training timeseries from the second platform and using the $\bar{g}_{ij}$ and $\sigma_{ij}^2$ values from this when timecourse-matching the test data.

##### S6.2 Intergene normalisation

When timecourse normalisation is unnecesary or impracticable the data is normalised using *intergene normalisation* where if $g = (g_i)$ is a expression vector the normalised levels are given by $\hat{g}_i = (g_i - \mu)/\sigma$ where $\mu$ and $\sigma^2$ are the mean and variance of the entries $g_i$. This is more likely to be effective for microarray data than RNA-seq data.

##### S6.3 Timecourse then intergene normalisation

It is also possible to usefully combine timecourse and intergene normalisation.

##### S6.4 Showing where the normalisations are used in the main paper

| **Use of normalisation** | | | |
|---|---|---|---|
| Dataset | Training normalisation | Test normalisation | Figure |
| Acosta-Rodeguez *et al.* | timecourse | timecourse-matched | 4 |
| Acosta-Rodeguez *et al.* | timecourse | timecourse | S31 |
| Bjarnason *et al.* | intergene | intergene | 1,2,4 |
| Boyle *et al.* | intergene | intergene | 3 |
| Fang *et al.* | intergene | intergene | 1,2 |
| Feng *et al.* | intergene | intergene | 3 |
| Hughes *et al.* | timecourse | timecourse-matched | S19 |
| Kinouchi *et al.* skel. mus. | timecourse | timecourse-matched | 2 |
| Kinouchi *et al.* liver | timecourse | timecourse-matched | S16 |
| Koronowski *et al. et al.* | timecourse | timecourse-matched | 2 |
| Le Martelot *et al.* | intergene | intergene | S18 |
| Mure *et al.* | timecourse | timecourse-matched | 4 |
| Weger *et al.* | timecourse | timecourse-matched | S33 |
| Yeung *et al.* | timecourse | timecourse-matched | S33 |
| Zhang *et al.* microarray | intergene | intergene | 1, 2 |
| Zhang *et al.* microarray | timecourse | timecourse-matched | 1, 2, 3, 4 |
| Zhang *et al.* RNA-seq | timecourse then intergene | timecourse then intergene | 1 |
| Zhang *et al.* RNA-seq | timecourse | timecourse-matched | 2,4, S16, S33 |

**Table S5:** This shows where the different normalisations are used in the data presented. Where test data normalisation is given for a training dataset this refers to how the test data comes from a leave one out analysis.

#### S7 Singular Value Decomposition

We will use Singular Value Decomposition (SVD) in a number of places and therefore we outline it here. SVD gives a decomposition of any $m \times k$ matrix $A$ into a product of the form $A = UDV^*$ where $U$ is a $m \times k$ column-orthonormal matrix ($UU^* = I_m$ and $U^*U = I_k$), $V$ is a $k \times k$ orthonormal matrix and $D = \text{diag}(\sigma_1, \dots, \sigma_k)$ is a diagonal matrix. (Throughout these notes $^*$ denotes the transpose of a matrix.) This is the version of SVD that is often called thin SVD (see (*20*) for more details). The ordered elements $\sigma_1 \geq \cdots \geq \sigma_k$ are the *singular values* of $A$. We note that the columns $V_j$,

$j = 1, \dots k$, of $V$ form an orthonormal basis for $\mathbb{R}^k$. If the last $n$ of the singular values are zero then the last $n$ columns are an orthonormal basis for the kernel of $A$. If $m < k$ then $n \geq k - m$.

In what follows we will refer to the columns $V_j$ (resp. $U_j$) of $V$ (resp. $U$) as the right (resp. left) singular vectors. Note that $AV_j = \sigma_j U_j$ and $U_j^* A = \sigma_j V_j^*$..

##### S7.1 Optimal projections via SVD and the projection $U_{d,i}$

Suppose $\mathbf{v} = v_1, v_2, \dots, v_N$ is a set of $N$ $n$-dimensional data vectors. Let $\mathbf{b} = b_1, \dots, b_n$ be an orthogonal basis of unit vectors for $\mathbb{R}^n$. Given $\mathbf{b}$ consider the $k$-dimensional projection of a vector $v$ in $\mathbf{v}$ given by

$$v_{i,k} = \sum_{j=1}^{k} \langle v_i, b_j \rangle b_j.$$

If the mean error $e_k(\mathbf{b})$ of this projection is defined by $e_k(\mathbf{b})^2 = N^{-1} \sum_i \|v_i - v_{i,k}\|^2$ we seek a basis which minimizes this error for all $k \leq n$.

Define $\sigma_i(\mathbf{b})$ by

$$\sigma_i(\mathbf{b})^2 = N^{-1} \sum_{j=1}^{N} \langle v_j, b_i \rangle^2.$$

then, by orthogonality of the $b_i$, $e_k(\mathbf{b})^2 = \sum_{i=k+1}^{n} \sigma_i(\mathbf{b})^2$.

As is well-known in (e.g., (*20*) (*21*)), the solution to this problem is provided by SVD. If $M$ is the $N \times n$ matrix whose columns are the vectors $v_i$, let $M = UDV^*$ be the SVD of $M$ ($*$ denoting transpose). Then $U$ is $N \times n$ and the solution to the above problem is $b_i = U_i$, the $i$th column of $U$, and the $\sigma_i(\mathbf{b})$ are the singular values of $M$ (i.e. the diagonal elements of $D$). Moreover, projecting into $\mathbb{R}^k$ using this basis gives

$$v_{i,k} = U^{[k]} v_i$$

where $U^{[k]}$ is the transpose of the matrix whose $k$ columns are $U_1, \dots, U_k$.

Thus for all $k = 1, \dots, n$ the projection of the vectors into $k$ dimensions using $U^{[k]}$ is the optimal representation of the vectors in $\mathbb{R}^k$ in the sense that the mean $L^2$ error is minimal.

The projection $U^{[d]}$ defined above is defined in terms of the vectors $\mathbf{v} = v_1, v_2, \dots, v_N$. If $\mathbf{v}$ consists of the vectors $X_{ij}$ for $j = 1, \dots, N$ (i.e. $v_j = X_{i,j}$), where $X_{ij}$ is as defined in Note S2 then we denote the projection $U^{[d]}$ by $U_{d,i}$.

#### S8 Analysis of synchronicity and rhythmicity

To measure synchronicity we used the approach using Singular Value Decomposition (SVD) as explained in section S7. We describe this in the context of the Zhang *et al.* microarray data. From this it is clear how to apply it to other training data.

After fRMA processing, the data contained 35,556 probe values for $m = 8$ organs at $T$ time points which was structured into 35,556 $T \times m$ matrices $M_g$, and normalised so that each $T$-dimensional column has mean 0 and standard deviation 1. The columns thus correspond to normalised timeseries and we have one for each organ. Then if the SVD of these matrices has the form $M_g = U_g D_g V_g^*$ as in Note S7, the optimal projection, as in Note S7.1 is given by using as basis the columns $U_{g,i}$ of $U_g$ which correspond to timeseries.

Thus, the first principal component $U_{g,1}$ describes the dominant temporal shape found across the various organs for gene $g$ and the other principal components $U_{g,i}$, $i > 1$, describe how this is modified across organs in a graded way with the strength given by the singular values $\sigma_{g,k}$. To characterise the relative strength of the first principal component we use

$$S_g^2 = \frac{\sigma_{g,1}^2}{\sum_k \sigma_{g,k}^2}$$

and this is our *synchronicity score*.

Alongside the synchronicity analysis we ranked genes by rhythmicity. Time-course normalised genes were ranked for goodness of 24 hr period cosine fit using cosinor analysis. Cosinor is a least squares regression approach, originally developed for use with short/sparse datasets (Cornelissen, 2014). The cosinor function written for MATLAB by Casey Cox (2008) was used to determine the genes with the best 24 hr cosine fits. One small change was made to the code, such that the p-value of the zero-amplitude f-test was calculated correctly using the fcdf function rather than the fpdf function. The p-values for the null hypothesis that that the gene was not rhythmic were calculated for each gene in each tissue. The mean p-value of all tissues was retained for each gene. This was compared for consistency with analysis by JTK using the package at https://github.com/mfcovington/jtk-cycle.

Using this we classified which genes optimise both rhythmicity and synchronicity using the scatter plots of synchronicity score against rhythmicity score. A combined ranking can be obtained by summing the two rankings and then the REP can be chosen from the top genes in this ranking. See Figs. S6, S7 and S8 for the Zhang *et al.* mouse microarray training data, the Zhang *et al.* mouse RNA-seq training data, and the Bjarnason *et al.* human training data.

#### S8.1 Zhang *et al.* microarray data

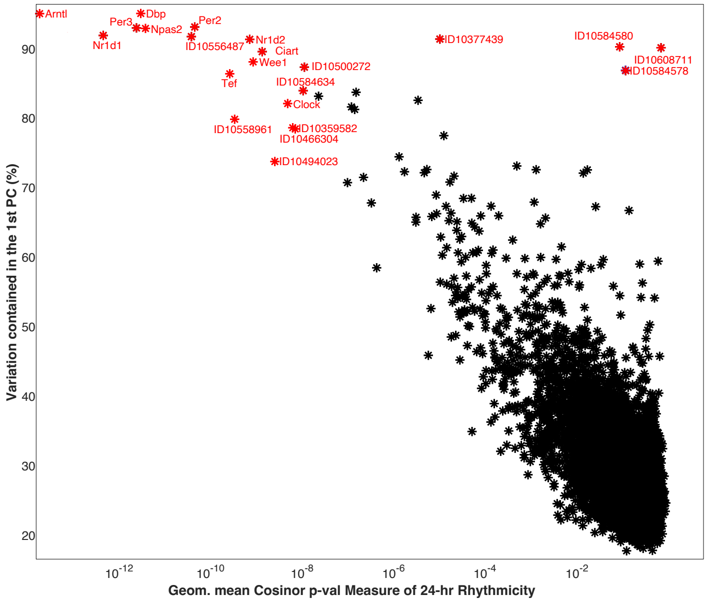


**Figure S6:** Scatter plot of each gene's score for rhythmicity (using the geometric mean of each tissue's Cosinor p-value) versus percentage of variation explained by the first principal component for the timeseries from the Zhang *et al.* microarray data. Genes highlighted in red are picked out for their extreme scores. Some are repeated because of the presence of multiple probes for that gene. The REP suggested for this data contains probes associated with the following genes (repeats correspond to multiple probes associated with that gene): *Arntl, Npas2, Per3, Dbp, Per2, Nr1d2, Tef, Wee1, Ciart ,Clock, Nr1d1*.

#### S8.2 Zhang *et al.* (2014) RNA-seq dataset

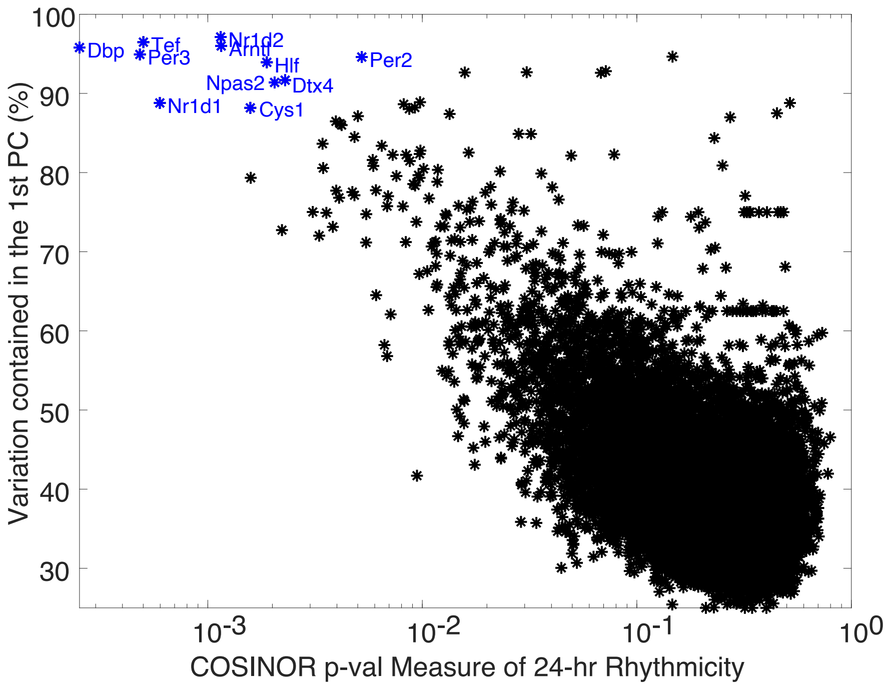


**Figure S7:** Scatter plot of genes expressed in 8 tissues from the Zhang *et al.* (2014) RNA-seq dataset. Genes are scored for rhythmicity (using the geometric mean of each tissue's Cosinor p-value) and percentage of variation explained by the first principal component. The REP suggested for this data contains probes associated with the following genes (repeats correspond to multiple probes associated with that gene): *Tef, Dbp, Nr1d2, Per3, Arntl, Hlf, Nr1d1, Dtx4, Npas2, Cys1, Per.*

#### S8.3 Bjarnason *et al.* data

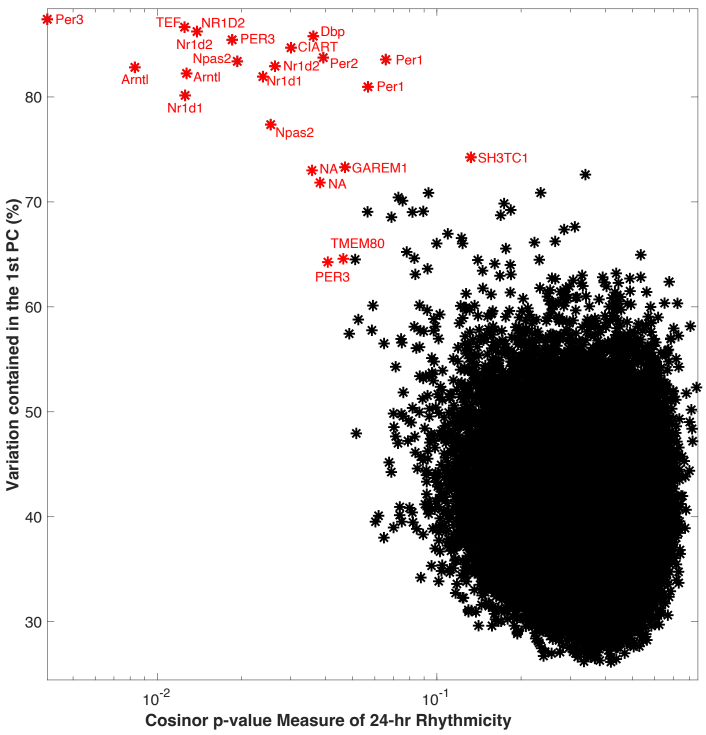


**Figure S8:** Scatter plot of each gene's score for rhythmicity (using the geometric mean of each tissue's Cosinor p-value) versus percentage of variation explained by the first principal component for the timeseries from the Bjarnason *et al.* data. Genes highlighted in red are picked out for their extreme scores. Some are repeated because of the presence of multiple probes for that gene. The REP suggested for the Bjarnason data contains 15 probes associated with the following genes: *Per3, Nr1d2, Ciart, Tef, Dbp, Arntl, Per2, Per1, Nr1d1, Nr1d2* and *Npas2*.

### S9 Extra information on the analysis of datasets

#### S9.1 Zhang *et al.* microarray data

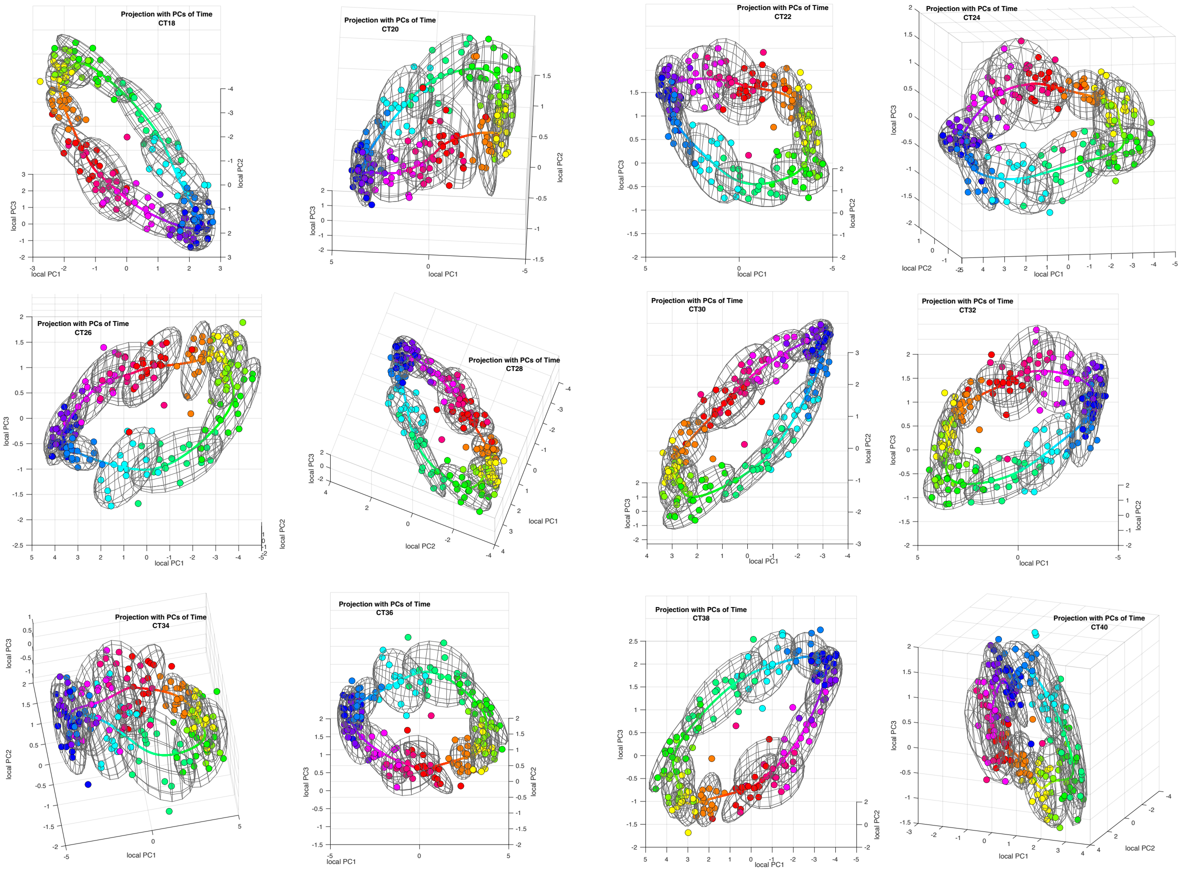


**Figure S9:** This shows all the twelve projections of the Zhang *et al.* microarray training data. There is one plot for each of the training times. When using the software they can be rotated to improve understanding of their geometry.

We now discuss treating the samples as test data using a leave-one-tissue out cross-validation approach. The choice of $l_{\text{thresh}}$ to use depends upon the test data. Since the test data here is from the training data we do not expect to see very many samples with very low likelihoods. Therefore, the choice of $l_{\text{thresh}}$ is likely to be much higher than we will choose when using this dataset for training and analyse test data with much lower likelihoods.

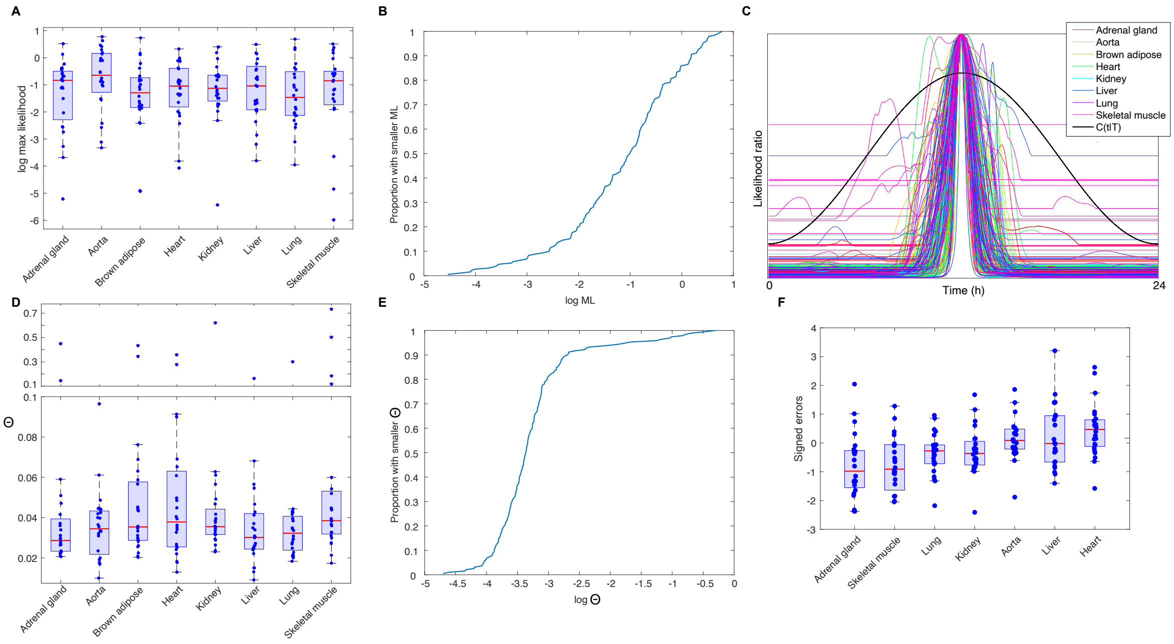


**Figure S10: Analysis of the Zhang *et al.* microarray data using a leave-one-tissue out cross-validation approach with $l_{\text{thresh}} = -5$. A.** Box plots showing the log maximum likelihood values $\log \text{ML}$ for each sample according to tissue. **B.** CDF of $\log \text{ML}$ values found. Since only a few percent are less than -4 we take $l_{\text{thresh}} = -5$. $l_{\text{thresh}} = -4$ would also have been appropriate. See Sect. S2.3 for details of how to choose $l_{\text{thresh}}$. **C.** Centered likelihood ratio curves (CLRFs). Note that although there are a number of samples with flat regions intersecting $C(t|T)$, by B, these are only a small proportion of all samples. About 15% of the CLRFs have two peaks. **D.** Box plots showing the $\Theta$ values for each tissue. **E.** CDF of $\log \Theta$ values found. Note that only about 5% have $\log \Theta > -2$. **E.** Box plots showing the signed timing errors for the different tissues. The means give the timing displacements.

| | Adrenal | Aorta | Brown adipose | Heart | Kidney | Liver | Lung | Skeletal muscle |
|---|---|---|---|---|---|---|---|---|
| ZeitZeiger | 1.08 | 0.53 | 0.60 | 0.68 | 0.66 | 0.66 | 0.62 | 1.50 |
| TimeTeller | 1.14 | 0.50 | 0.83 | 0.76 | 0.67 | 0.85 | 0.61 | 1.75 |
| TimeTeller corrected | 0.88 | 0.50 | 0.58 | 0.68 | 0.57 | 0.88 | 0.53 | 1.55 |

**Table S6:** This compares the mean absolute timing errors for the eight tissues used in the microarray training data from Zhang *et al.* . The top row are for ZeiZeiger and are extracted from Fig. S3 of (*22*). The 2nd and 3rd row are for TimeTeller using timecourse normalisation and the timing errors in the 3rd row have been corrected using the timing deviations for the tissues.

#### S9.2 Testing Zhang *et al.* RNA-seq data when trained on Zhang *et al.* microarray data and vice-versa

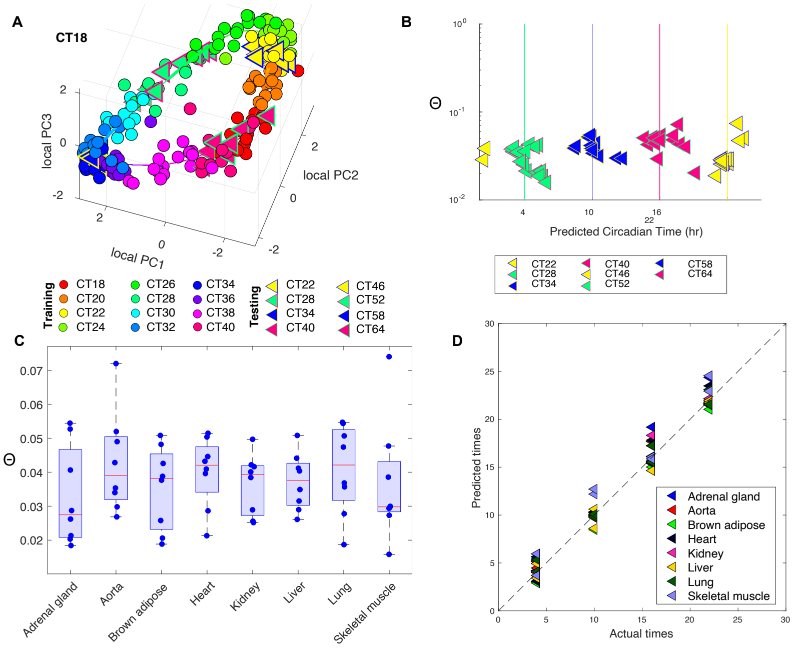

**Figure S11:** Results when testing Zhang *et al.* RNA-seq data when trained on Zhang *et al.* microarray data

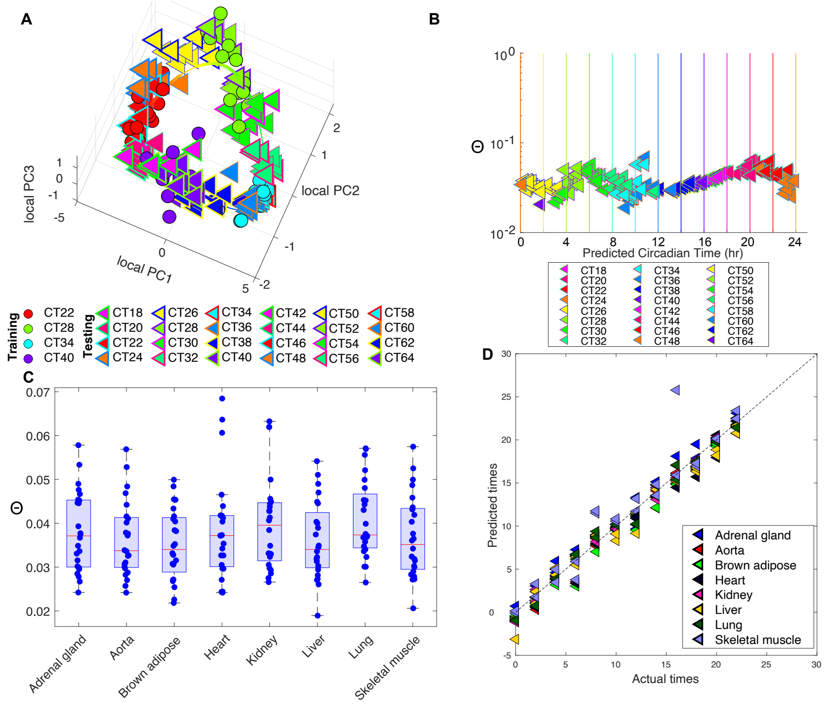

**Figure S12:** Results when testing Zhang *et al.* microarray data when trained on Zhang *et al.* RNA-seq data

#### S9.3 Bjarnason *et al.* data

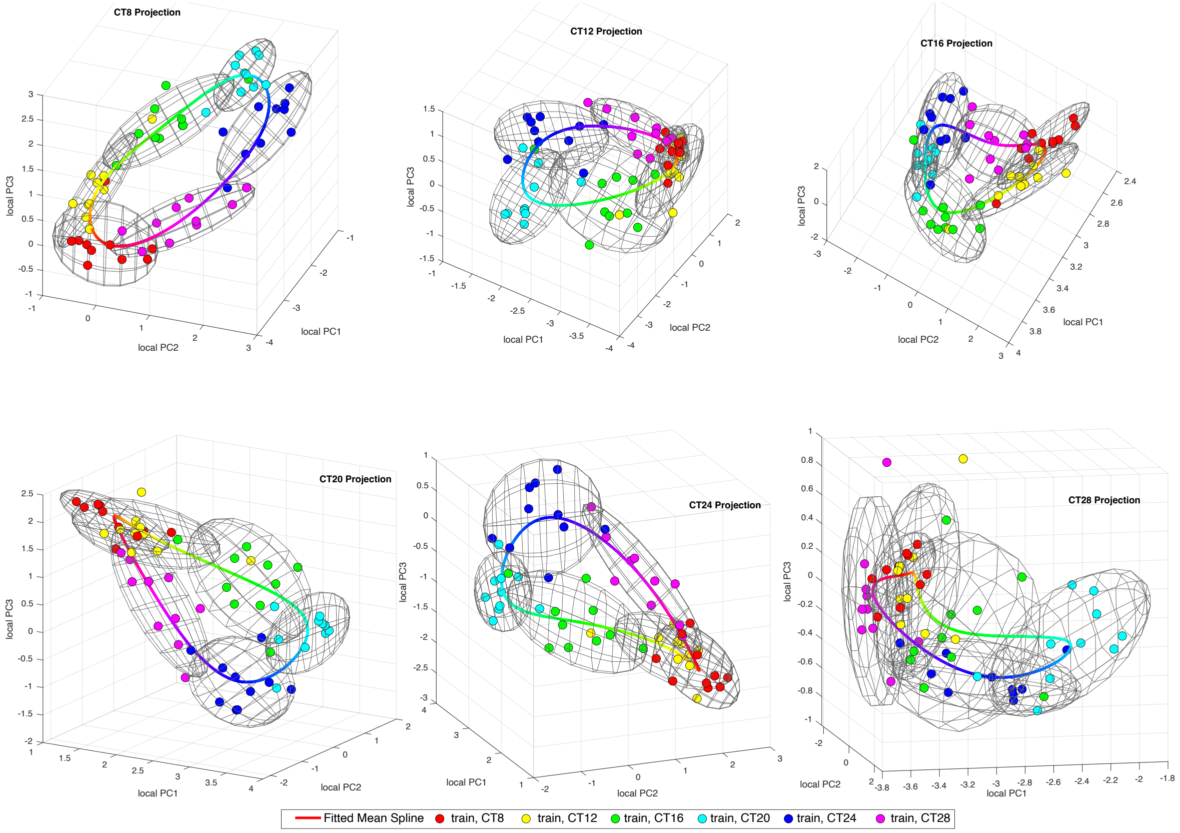


**Figure S13:** This shows all the six projections of the Bjarnason *et al.* training data. There one for each of the training times. When using the software they can be rotated to improve understanding of their geometry.

##### Leave-one-individual-out analysis of Bjarnason *et al.* data

For the same reasons that we mentioned above when discussing the Zhang *et al.* microarray data, the choice of $l_{\text{thresh}}$ for a leave-one-individual-out analysis of the Bjarnason *et al.* data is likely to be much higher than what we will choose when using this dataset for training and analysing test data with much lower likelihoods. We choose $l_{\text{thresh}} = -5$ here but for some test sets we analyse we use $l_{\text{thresh}}$ as low as -12. Therefore, we include in Fig. an analysis of how the $\Theta$ stratification changes as we lower $l_{\text{thresh}}$ from -5 to -12. The reason we choose $l_{\text{thresh}} = -5$ here is because (i) higher values have too many samples in the training data with flat regions significantly intersecting $C(t|T)$ and (ii) lower values are too far below the minimum $\mathrm{ML}$ of the test data.

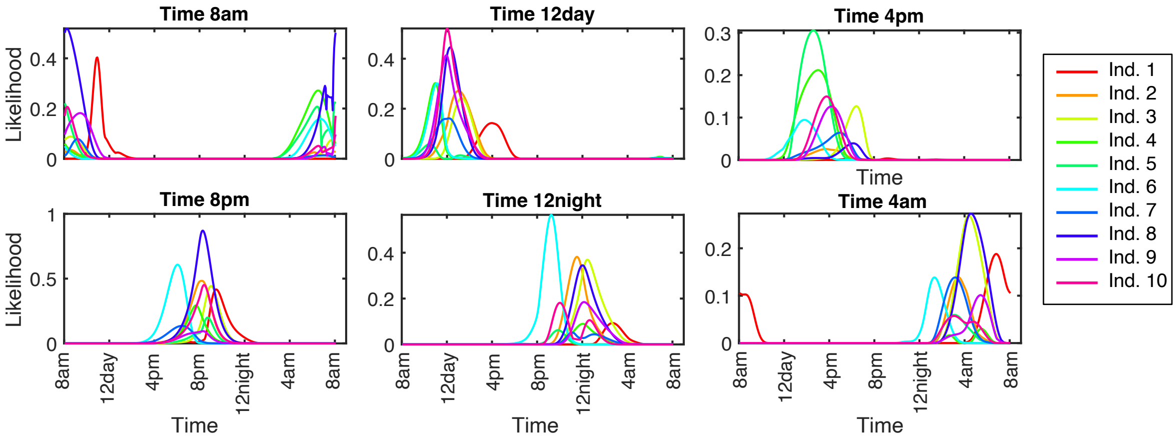


**Figure S14:** The likelihood curves for the leave-one-out analysis of the Bjarnason *et al.* data. The sample time is shown above each subplot.

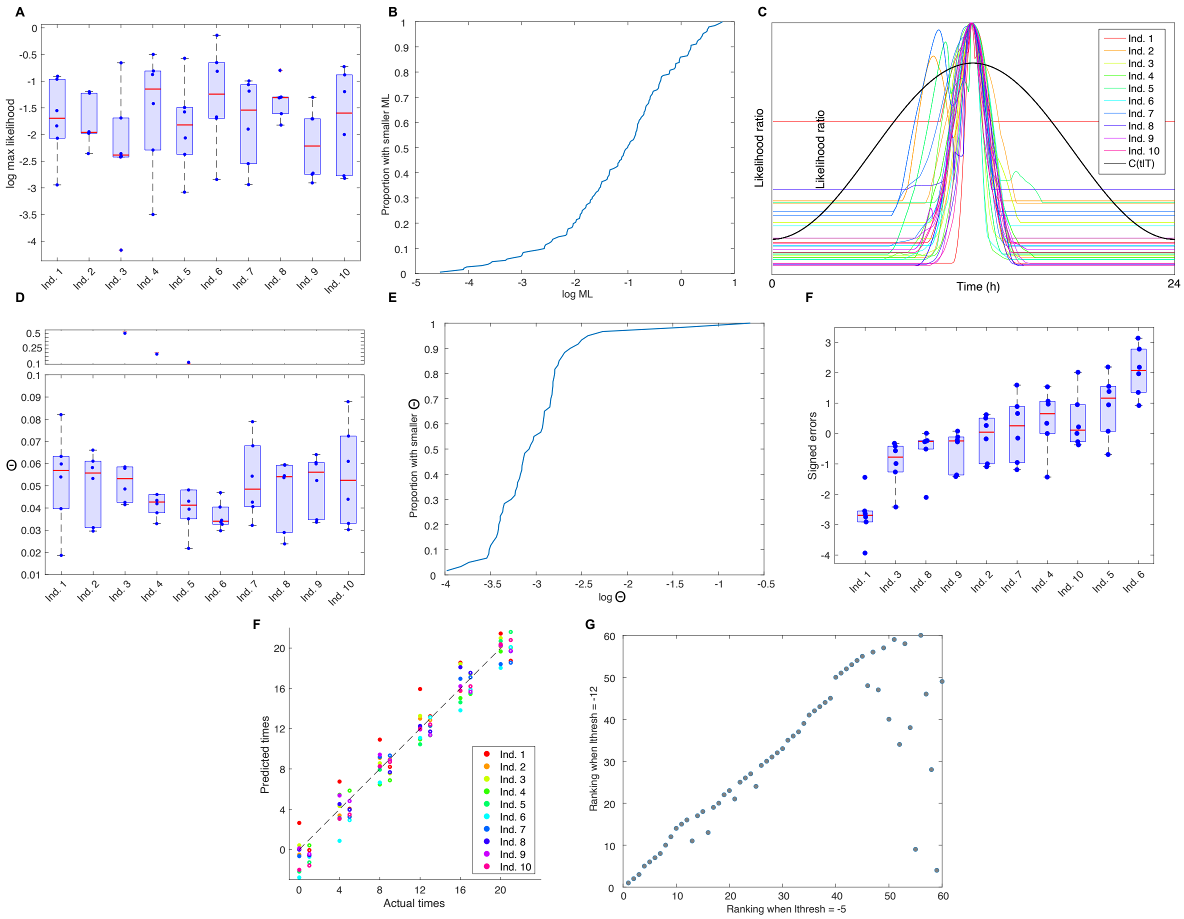

**Figure S15: Leave-one-individual-out analysis of Bjarnason *et al.* training data.** Intergene normalisation is used with $l_{\text{thresh}} = -5$. **A.** Boxplots showing the distribution of the log maximum likelihood values $\log \mathrm{ML}$ by individual. **B.** CDF of the $\log \mathrm{ML}$ values. **C.** Centred likelihood ratio functions for all individuals at all timepoints. **D.** Boxplots showing the distribution of $\Theta$ values by individual. **E.** CDF of the $\Theta$ values. **F.** Boxplots showing the distribution of the signed errors for each individual. The means increase from right to left. **G.** Analysis of the timing for the Bjarnason *et al.* human microarray data with each data point assigned the color corresponding to the individual. For each time the points over the time are for uncorrected timing and the points moved slightly to the right show the corrected timings (i.e. the predicted timing when corrected by the timing displacement for the individual. **H.** Scatter plot showing the change in the $\Theta$ stratification of samples when $l_{\text{thresh}}$ is changed from -5 to -12. While 14 points change their ranking for almost all the change preserves whether they are high or low. Only two samples change from high to low.

#### S9.4 Kinouchi *et al.* liver data

For the liver data an inspection of the ML values suggests using $l_{\text{thresh}} = -6$. In first figure below (Fig. S16) this value is used. However, in a second figure (Fig. S17) we apply the inappropriate value of -12 in order to demonstrate what happens if $l_{\text{thresh}}$ is taken too small. This forms part of the discussion started in Sect. S2.3 about how to choose $l_{\text{thresh}}$.

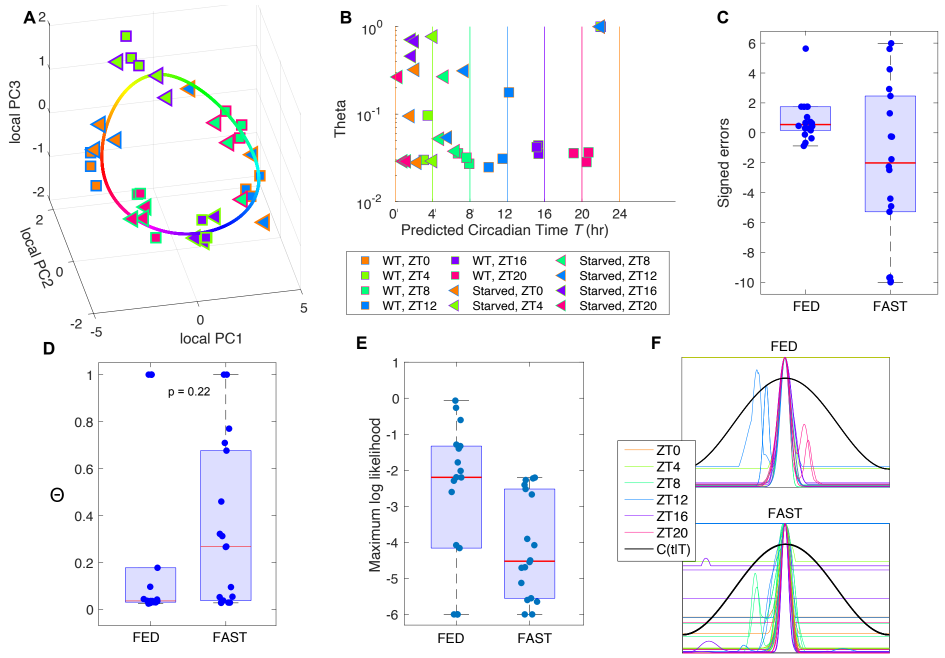


**Figure S16: Analysis of Kinouchi *et al.* liver data.** This uses $l_{\text{thresh}} = -6$. **A.** Visualisation of the Kinouchi *et al.* liver data plotted against the curve given by the means of $P(g|t)$ for the Zhang *et al.* RNA-seq training data. **B-E.** These use $l_{\text{thresh}} = -6$. **B.** Plot of the $\Theta$ values against the estimated time $T$ for each test sample. The vertical lines show the true time with colours indicating the sampling time. **C.** Boxplots of the signed errors showing that the absolute errors of the errors for FAST data are significantly higher than those for FED data. The mean values for FED and FAST are respectively 1.23h and 4.70h and the medians 0.78h and 4.46h. The difference is significant at the $p < 0.0008$ level. **C & D.** Boxplots of the $\Theta$ and maximum likelihood values. **E.** The likelihood curves for the FED and FAST samples. These have been been translated in time so that its highest peak is at 12noon as this makes comparison of the shapes easier. **F.** Centred LRFs for the FED and FAST mice.

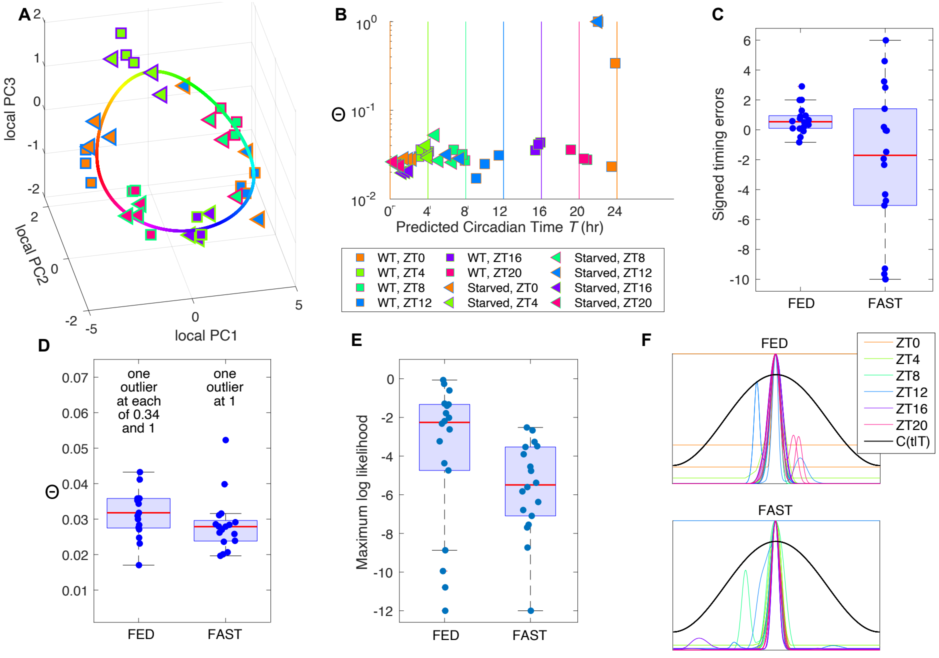

**Figure S17: Analysis of Kinouchi *et al.* liver data when $l_{\mathbf{thresh}} = -12$. This threshold is too low** and this figure is included as part of the discussion in Sect. S2.3 about how to choose $l_{\text{thresh}}$. **A.-F.** As Fig. S16 except that in **C.** the mean values for FED and FAST are respectively 0.83h and 4.29h and the medians 0.62h and 3.78h. The difference is significant at the $p < 0.0015$ level.

#### S9.5 Le Martelot *et al.* data

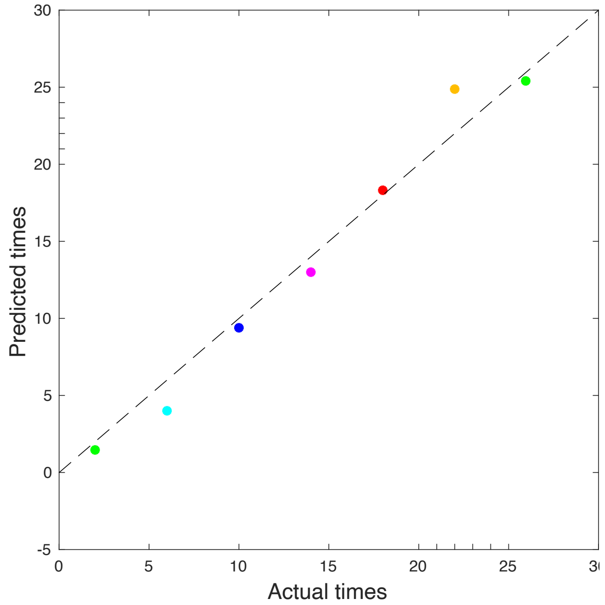


**Figure S18:** This shows the small timing error found in the independent LeMartelot *et al.* data (*9*).

#### S9.6 Hughes *et al.* data (*1*)

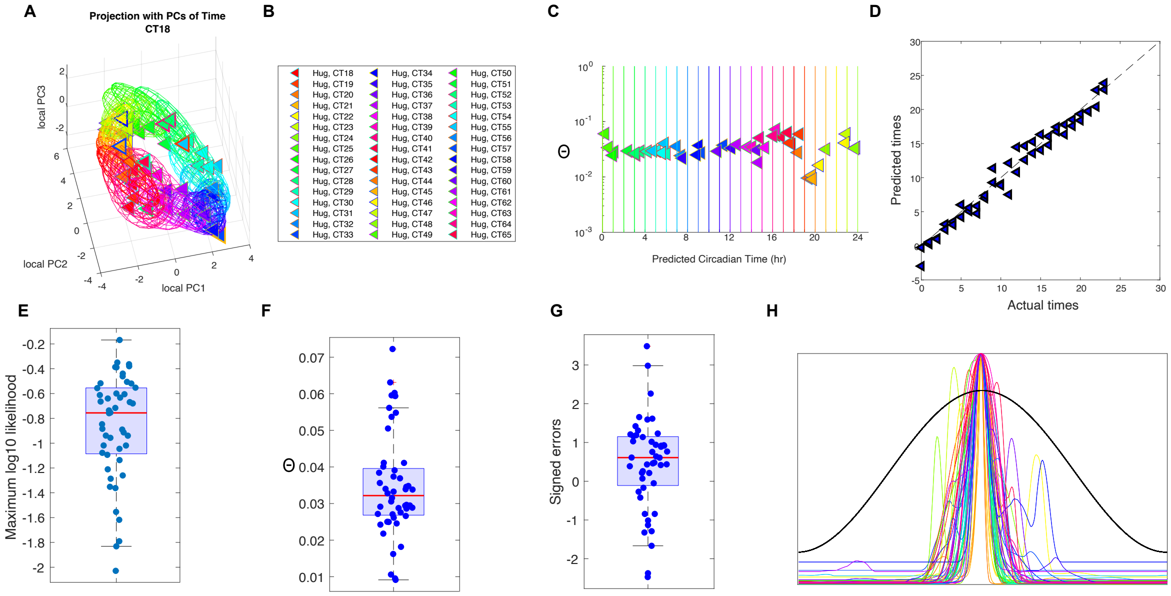


**Figure S19: Analysis of Hughes *et al.* liver data (*1*) when $l_{\textbf{thresh}} = -7$.** Timecourse normalisation is used for both training and test data. **A.** Visualisation of the Hughes *et al.* liver data plotted against the curve given by the means of $P(g|t)$ for the Zhang *et al.* RNA-seq training data. **B.** Legend. Colors are consistent across all plots. **C.** Plot of the $\Theta$ values against the estimated time $T$ for each test sample. The vertical lines show the true time with colours indicating the sampling time. **D.** Predicted $T$ times plotted against actual times. **E.** Boxplot of the maximum liklihoods. **F.** Boxplot of the $\Theta$ values. **G.** Boxplot of the signed errors. **H.** Centred likelihood curves for each sample.

#### S9.7 Boyle *et al.* data (2)

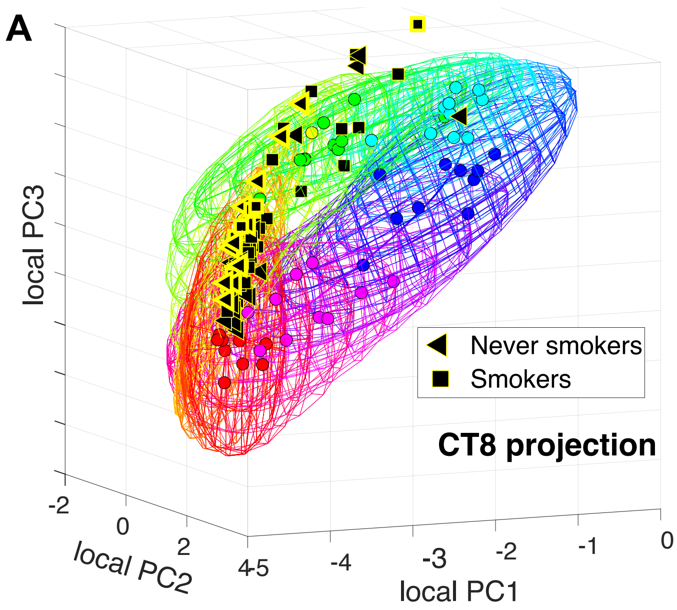


**Figure S20: Analysis of Boyle *et al.* smoking data (2). A.** Visualisation of the Boyle *et al.* data showing how this data sits in the Bjarnason *et al.* microarray training data.

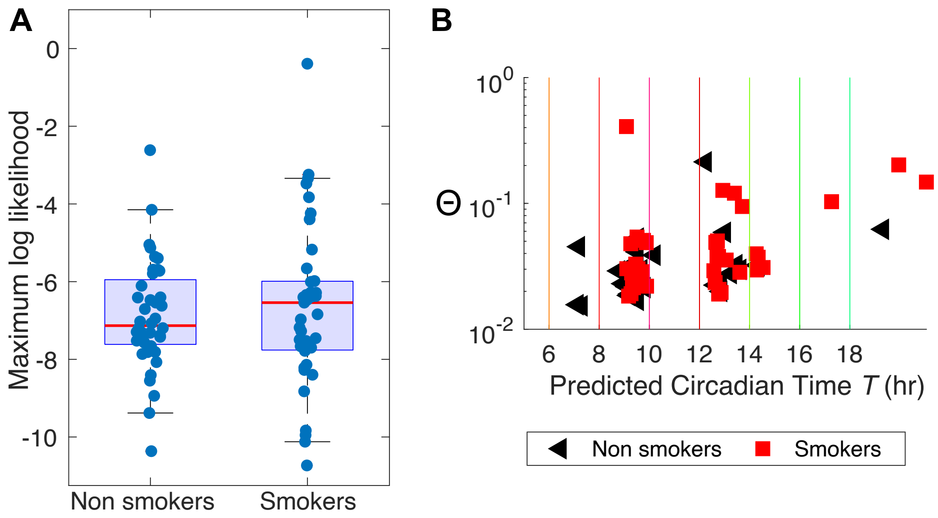


**Figure S21: Analysis of Boyle *et al.* smoking data (2).** This uses intergene normalisation and $l_{\text{thresh}} = -12$. **A.** ML values for smokers and non-smokers. **B.** Scatter plot of $\Theta$ values against estimated timing $T$.

#### S9.8 Feng *et al.* data (*3*)

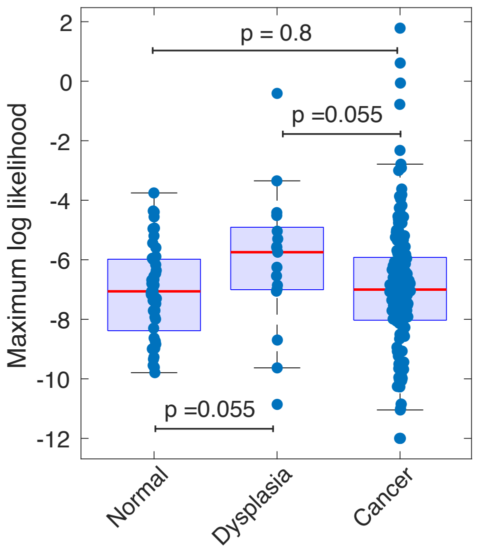


**Figure S22: Analysis of Feng *et al.* data (*3*).** ML values for the normal, dysplasic and cancer data. This uses intergene normalisation and $l_{\text{thresh}} = -12$.

#### S9.9 Differentially expressed genes in the Boyle *et al.* data (*2*) and the Feng *et al.* data (*3*)

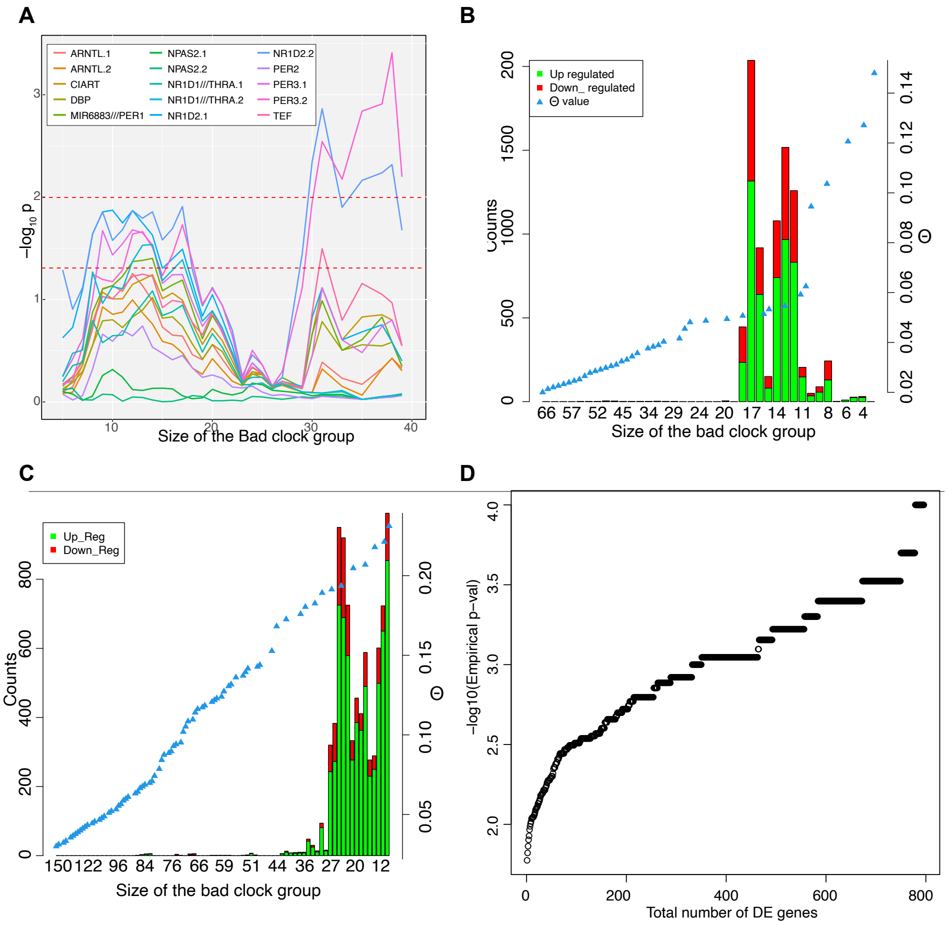


**Figure S23: A,B. Analysis of Boyle *et al.* smoking data (*2*).** This uses $l_{\text{thresh}} = -12$. **A.** The adjusted p-values for clock genes which are differentially expressed between those $n$ individuals with the worse clocks (according to the $\Theta$ stratification) and those with better clocks. The p-values were calculated using the limma package (v3.48.3) and have been adjusted to account for the multiple testing with the BH method used for adjustment. Some of the genes had two probes which are distinguished by a point followed by either 1 or 2. The broken red lines show where $p = 0.05$ and $p = 0.01$. **B.** The number of statistically significant differentially expressed genes as a function of $n$, the size of the bad clock group. The threshold for differential expression is p $< 0.05$ for the adjusted p-value (limma using BH). The blue triangles indicate the corresponding $\Theta$ values. **C,D. Analysis of Feng *et al.* data (*3*).** This uses $l_{\text{thresh}} = -12$. **C.** As B but for the Feng *et al.* data. **D.** Empirical probability of observing $m$ or more differentially expressed genes (DEGs) 10,000 simulations were run in which the number of DEGs were calculated after the bad clock group size was randomised to between 15 and 40 and individuals were randomly allocated to the bad and good clock groups. The empirical distribution plot shows the results of observing $m$ or more DE genes as per the distribution obtained. Plotted on -log10 scale.

#### S9.10 Acosta-Rodríguez *et al.* data

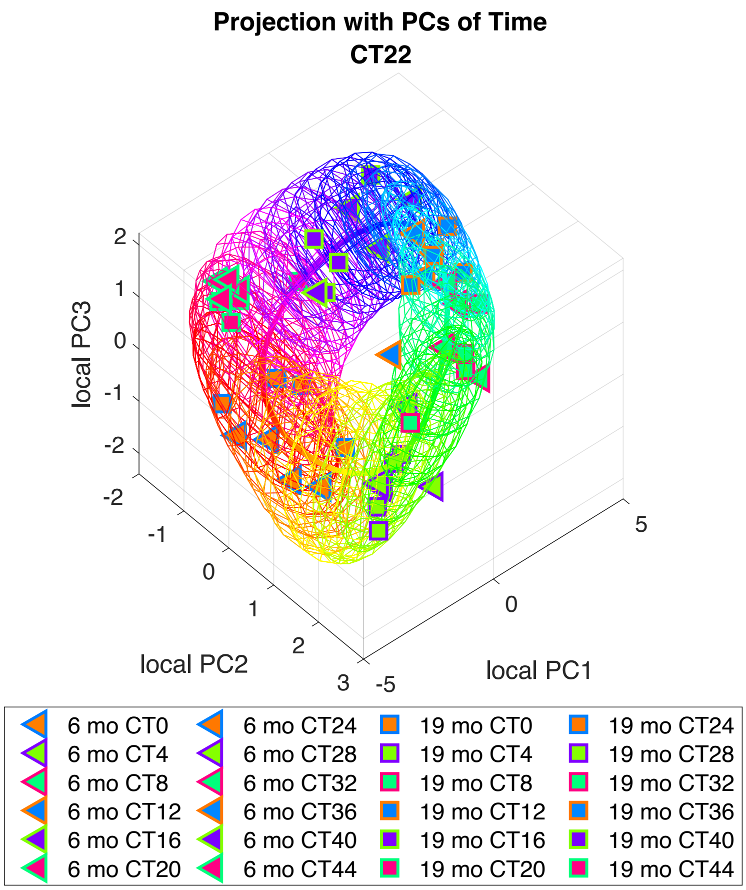


**Figure S24:** Visualisation of the 3 standard deviation ellipsoids for the probability model for the Zhang *et al.* RNA-seq training data together with the Acosta-Rodríguez *et al.* data (triangles and squares) treated as test data and normalised using timecourse matching.

| | AL6m | AL19m | CRday2h6m | CRday2h19m | CRday12h6m | CRday12h19m | CRnight2h6m | CRnight2h19m | CRnight12h6m | CRnight12h19m | CRspread6m | CRspread19m |
|---|---|---|---|---|---|---|---|---|---|---|---|---|
| AL6m | | | 3.90E-05 | 3.06E-09 | 3.93E-09 | 3.47E-09 | 4.66E-02 | 8.06E-03 | 4.88E-03 | | 2.26E-05 | 3.75E-04 |
| AL19m | | | 4.26E-05 | 3.06E-09 | 5.05E-09 | 3.06E-09 | 2.81E-02 | 2.05E-03 | 2.27E-02 | | 1.71E-05 | 1.55E-04 |
| CRday2h6m | 3.90E-05 | 4.26E-05 | | 4.26E-05 | | | 6.44E-04 | 5.53E-04 | 1.33E-06 | 8.85E-06 | 9.10E-03 | 2.05E-03 |
| CRday2h19m | 3.06E-09 | 3.06E-09 | 4.26E-05 | | 2.17E-08 | 1.10E-07 | 3.06E-09 | 3.06E-09 | 3.06E-09 | 3.06E-09 | 6.46E-09 | 3.06E-09 |
| CRday12h6m | 3.93E-09 | 5.05E-09 | | 2.17E-08 | | | 1.38E-07 | 1.51E-08 | 3.06E-09 | 3.06E-09 | 2.72E-06 | 2.15E-07 |
| CRday12h19m | 3.47E-09 | 3.06E-09 | | 1.10E-07 | | | 1.92E-08 | 4.46E-09 | 3.06E-09 | 3.06E-09 | 9.72E-07 | 1.71E-08 |
| CRnight2h6m | 4.66E-02 | 2.81E-02 | 6.44E-04 | 3.06E-09 | 1.38E-07 | 1.92E-08 | | | 3.21E-04 | 1.78E-03 | 3.64E-02 | |
| CCRnight2h19m | 8.06E-03 | 2.05E-03 | 5.53E-04 | 3.06E-09 | 1.51E-08 | 4.46E-09 | | | 2.26E-05 | 1.98E-04 | | |
| CRnight12h6m | 4.88E-03 | 2.27E-02 | 1.33E-06 | 3.06E-09 | 3.06E-09 | 3.06E-09 | 3.21E-04 | 2.26E-05 | | | 1.72E-07 | 3.32E-06 |
| CRnight12h19m | | | 8.85E-06 | 3.06E-09 | 3.06E-09 | 3.06E-09 | 1.78E-03 | 1.98E-04 | | | 7.88E-07 | 1.18E-05 |
| CRspread6m | 2.26E-05 | 1.71E-05 | 9.10E-03 | 6.46E-09 | 2.72E-06 | 9.72E-07 | 3.64E-02 | | 1.72E-07 | 7.88E-07 | | |
| CRspread19m | 3.75E-04 | 1.55E-04 | 2.05E-03 | 3.06E-09 | 2.15E-07 | 1.71E-08 | | | 3.32E-06 | 1.18E-05 | | |

**Table S7:** This table shows p-values for the Wilcoxon rank sum test (Matlab function ranksum) testing the null hypothesis that the timing errors in the indicated pair of boxes arise from distributions with equal medians. Only those with $p < 0.05$ are shown.

| | AL6m | AL19m | CRday2h6m | CRday2h19m | CRday12h6m | CRday12h19m | CRnight2h6m | CRnight2h19m | CRnight12h6m | CRnight12h19m | CRspread6m | CRspread19m |
|---|---|---|---|---|---|---|---|---|---|---|---|---|
| AL6m | | | 2.04E-02 | | | | 9.10E-03 | | | | | 5.55E-03 |
| AL19m | | | 5.21E-03 | 1.30E-02 | | | 1.68E-04 | 1.30E-02 | | | | 1.30E-02 |
| CRday2h6m | 2.04E-02 | 5.21E-03 | | | | | | | 3.12E-02 | | 4.02E-02 | 2.96E-04 |
| CRday2h19m | | 1.30E-02 | | | | | | | | | | 4.39E-04 |
| CRday12h6m | | | | | | | | | | | | 5.97E-04 |
| CRday12h19m | | | | | | | 2.96E-02 | | | | | 1.55E-03 |
| CRnight2h6m | 9.10E-03 | 1.68E-04 | | | | 2.96E-02 | | | 7.13E-03 | 2.40E-02 | 1.22E-02 | 8.04E-06 |
| CRnight2h19m | | 1.30E-02 | | | | | | | | | | 6.94E-04 |
| CRnight12h6m | | | 3.12E-02 | | | | 7.13E-03 | | | | | 3.77E-03 |
| CRnight12h19m | | | | | | | 2.40E-02 | | | | | 3.09E-03 |
| CRspread6m | | | 4.02E-02 | | | | 1.22E-02 | | | | | 5.97E-04 |
| CRspread19m | 5.55E-03 | 1.30E-02 | 2.96E-04 | 4.39E-04 | 5.97E-04 | 1.55E-03 | 8.04E-06 | 6.94E-04 | 3.77E-03 | 3.09E-03 | 5.97E-04 | |

**Table S8:** This table shows p-values for the Wilcoxon rank sum test (Matlab function ranksum) testing the null hypothesis that the maximum likelihoods in the indicated pair of boxes arise from distributions with equal medians. Only those with $p < 0.05$ are shown.

| | AL6m | AL19m | CRday2h6m | CRday2h19m | CRday12h6m | CRday12h19m | CRnight2h6m | CRnight2h19m | CRnight12h6m | CRnight12h19m | CRspread6m | CRspread19m |
|---|---|---|---|---|---|---|---|---|---|---|---|---|
| AL6m | | | | | | | 1.30E-02 | 3.82E-02 | | | | |
| AL19m | | | | | | | 2.20E-03 | 7.13E-03 | | | | |
| CRday2h6m | | | | | | | | | | | | |
| CRday2h19m | | | | | | | | | | | | |
| CRday12h6m | | | | | | | | | | | | |
| CRday12h19m | | | | | | | 4.89E-02 | | | | | |
| CRnight2h6m | 1.30E-02 | 2.20E-03 | | | | 4.89E-02 | | | 4.66E-02 | 4.89E-02 | 1.22E-02 | 5.91E-03 |
| CRnight2h19m | 3.82E-02 | 7.13E-03 | | | | | | | | | | 2.15E-02 |
| CRnight12h6m | | | | | | | 4.66E-02 | | | | | |
| CRnight12h19m | | | | | | | 4.89E-02 | | | | | |
| CRspread6m | | | | | | | 1.22E-02 | | | | | |
| CRspread19m | | | | | | 5.91E-03 | 2.15E-02 | | | | | |

**Table S9:** This table shows p-values for the Wilcoxon rank sum test (Matlab function ranksum) testing the null hypothesis that the $\Theta$ values in the indicated pair of boxes arise from distributions with equal medians. Only those with $p < 0.05$ are shown.

| | AL6mo | AL-19m | Cr-d-2h-6m | Cr-d-2h-19m | Cr-d-12h-6m | Cr-d-12h-19m |
|---|---|---|---|---|---|---|
| Tef | 4.10E-05 | 2.57E-07 | 2.91E-01 | 1.64E-03 | 5.76E-03 | 3.12E-03 |
| Dbp | 2.60E-05 | 4.92E-08 | 2.13E-02 | 1.61E-02 | 4.29E-04 | 7.86E-04 |
| Nr1d2 | 6.21E-05 | 7.23E-06 | 7.77E-01 | 7.38E-03 | 2.33E-03 | 3.75E-04 |
| Per3 | 9.01E-05 | 5.41E-07 | 2.53E-02 | 1.69E-02 | 3.52E-03 | 1.34E-03 |
| Arntl | 7.69E-06 | 7.15E-05 | 1.59E-01 | 1.69E-03 | 5.12E-04 | 3.14E-04 |
| Hlf | 1.58E-03 | 3.05E-03 | 1.47E-01 | 1.90E-03 | 2.03E-02 | 4.89E-02 |
| Nr1d1 | 1.12E-04 | 2.77E-07 | 9.71E-02 | 7.07E-04 | 3.58E-04 | 1.55E-03 |
| Dtx4 | 1.02E-05 | 6.51E-06 | 4.64E-01 | 7.24E-02 | 2.07E-03 | 1.73E-03 |
| Npas2 | 2.26E-06 | 4.59E-07 | 2.81E-01 | 4.24E-02 | 2.95E-04 | 1.18E-03 |
| Cys1 | 4.55E-02 | 1.79E-01 | 5.12E-01 | 8.37E-01 | 4.69E-01 | 2.79E-01 |
| Per2 | 7.80E-04 | 1.50E-05 | 1.00E-03 | 1.45E-02 | 3.40E-04 | 4.25E-04 |

| | Cr-nt-2h-6m | Cr-nt-2h-19m | Cr-nt-12h-6m | Cr-nt-12h-19m | CR-sp-6m | CR-sp-19m |
|---|---|---|---|---|---|---|
| Tef | 1.40E-03 | 1.89E-05 | 4.82E-05 | 7.55E-05 | 5.15E-02 | 2.20E-04 |
| Dbp | 5.39E-04 | 1.91E-05 | 1.90E-06 | 2.79E-05 | 1.27E-02 | 3.81E-05 |
| Nr1d2 | 1.24E-04 | 6.64E-06 | 2.76E-06 | 5.20E-05 | 2.00E-02 | 7.79E-05 |
| Per3 | 1.28E-03 | 1.05E-04 | 1.16E-04 | 2.13E-05 | 4.27E-02 | 3.95E-04 |
| Arntl | 2.12E-03 | 1.42E-03 | 9.51E-05 | 3.74E-04 | 1.43E-02 | 3.06E-04 |
| Hlf | 9.16E-02 | 1.31E-04 | 2.56E-03 | 1.88E-03 | 9.37E-02 | 1.92E-02 |
| Nr1d1 | 9.90E-05 | 2.09E-06 | 5.87E-06 | 1.31E-05 | 3.29E-05 | 1.97E-04 |
| Dtx4 | 9.83E-03 | 2.33E-03 | 9.78E-06 | 8.80E-04 | 5.50E-02 | 7.83E-05 |
| Npas2 | 3.65E-05 | 3.96E-07 | 5.78E-06 | 2.94E-04 | 4.80E-02 | 4.60E-06 |
| Cys1 | 1.96E-01 | 7.37E-02 | 4.97E-03 | 9.82E-04 | 8.45E-02 | 6.46E-02 |
| Per2 | 2.01E-04 | 2.64E-05 | 6.51E-05 | 8.74E-06 | 1.57E-02 | 4.76E-05 |

**Table S10:** These two tables give the Cosinor p-values for the genes used in the REP for each of the feeding conditions in the Acosta-Rodríguez *et al.* data.

#### S9.11 Assessing the coherence of downstream genes in the Acosta-Rodríguez *et al.* data

##### Cell cycle genes

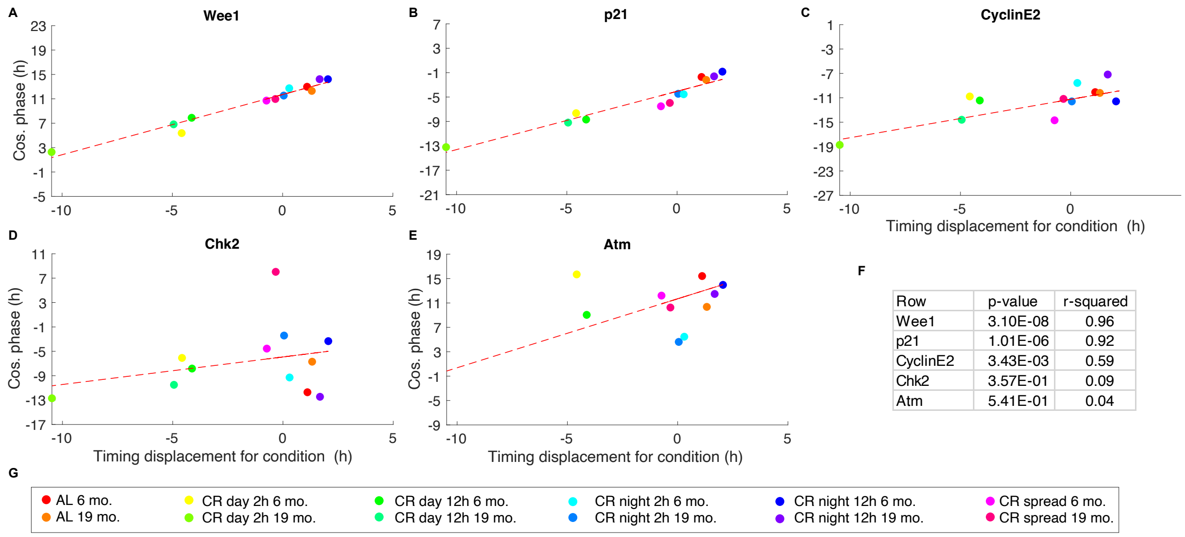


**Figure S25: Cell cycle genes. A-E.** PDP plots in which the timing displacement for the conditions is linearly regressed against the gene phase of each gene named in the plots. The Zhang *et al.* data was used for training and $l_{\text{thresh}}$ = -8. Of these genes *Wee1*, *p21* and *CyclinE2* appeared to be oscillating in all AL conditions and they appear to be changing phase in coordination with the clock. For the other genes, all of which appear to change in an uncoordinated way, it should be noted that the shown gene phase is not really meaningful as they are unlikely to be rhythmic for most or all the conditions. **F.** p-values and $r^2$ values for A-E. **G.** Legend showing which condition the points in A-E correspond to.

| | Wee1 | Cys1 | Timeless | CyclinA | CyclinB1 | CyclinE2 | CHK2 | ATM | P53 | p21 |
|---|---|---|---|---|---|---|---|---|---|---|
| AL 6mo. r1 | 1.83E-03 | 5.18E-02 | 6.13E-01 | 2.55E-01 | 6.22E-01 | 1.90E-02 | 7.42E-01 | 5.15E-01 | 2.47E-03 | 2.57E-03 |
| AL 6mo. r2 | 1.00E-03 | 4.00E-02 | 3.61E-01 | 2.43E-01 | 9.71E-01 | 5.84E-04 | 4.01E-01 | 8.01E-01 | 5.22E-02 | 3.88E-04 |
| AL 19mo. r1 | 8.82E-08 | 4.26E-02 | 4.14E-03 | 3.02E-01 | 4.65E-01 | 3.42E-03 | 2.21E-02 | 6.04E-03 | 7.80E-01 | 3.90E-02 |
| AL 19mo. r2 | 1.58E-04 | 7.52E-01 | 3.45E-02 | 1.09E-01 | 1.51E-01 | 2.54E-02 | 9.96E-01 | 9.30E-01 | 7.97E-02 | 4.69E-03 |
| CRday 2h 6mo. r1 | 4.77E-01 | 3.63E-01 | 7.69E-02 | 7.21E-01 | 5.63E-01 | 6.59E-01 | 2.66E-04 | 5.49E-01 | 4.24E-01 | 3.21E-01 |
| CRday 2h 6mo. r2 | 1.95E-04 | 7.21E-01 | 6.31E-01 | 3.41E-01 | 1.96E-01 | 5.40E-01 | 1.20E-01 | 4.06E-01 | 3.19E-02 | 1.97E-01 |
| CRday 2h 19mo. r1 | 3.08E-03 | 9.90E-01 | 7.29E-01 | 1.20E-01 | 4.02E-01 | 4.69E-01 | 2.33E-01 | 3.90E-03 | 3.19E-02 | 1.99E-03 |
| CRday 2h 19mo. r2 | 1.65E-02 | 7.08E-01 | 1.87E-01 | 1.31E-01 | 1.38E-01 | 5.73E-02 | 6.48E-01 | 1.07E-02 | 2.99E-01 | 1.53E-01 |
| CRday 12h 6mo. r1 | 9.67E-03 | 5.89E-01 | 1.49E-02 | 6.85E-02 | 1.00E+00 | 9.50E-02 | 2.04E-01 | 3.19E-02 | 8.15E-01 | 9.87E-02 |
| CRday 12h 6mo. r2 | 2.61E-02 | 3.74E-01 | 1.27E-01 | 4.12E-01 | 5.58E-01 | 7.99E-03 | 1.99E-01 | 8.45E-01 | 6.98E-01 | 1.04E-02 |
| CRday 12h 19mo. r1 | 2.32E-02 | 8.74E-01 | 7.91E-02 | 5.42E-01 | 2.17E-01 | 5.95E-01 | 7.71E-01 | 6.34E-03 | 1.41E-01 | 2.92E-02 |
| CRday 12h 19mo. r2 | 4.63E-03 | 8.92E-02 | 5.54E-01 | 3.05E-01 | 3.39E-01 | 6.89E-02 | 4.81E-01 | 1.49E-01 | 7.36E-01 | 1.16E-01 |
| CRnight 2h 6mo. r1 | 1.11E-02 | 1.87E-01 | 1.93E-01 | 6.13E-01 | 1.26E-01 | 8.37E-03 | 8.26E-01 | 2.63E-01 | 6.18E-01 | 9.75E-01 |
| CRnight 2h 6mo. r2 | 1.49E-03 | 2.06E-01 | 5.68E-01 | 3.65E-01 | 5.73E-01 | 6.76E-01 | 2.05E-01 | 1.13E-01 | 1.86E-01 | 1.62E-02 |
| CRnight 2h 19mo. r1 | 1.63E-05 | 4.59E-02 | 6.72E-01 | 3.50E-01 | 7.62E-01 | 6.90E-04 | 8.67E-01 | 1.09E-03 | 1.10E-02 | 3.73E-03 |
| CRnight 2h 19mo. r2 | 1.79E-03 | 1.18E-01 | 3.43E-01 | 9.54E-01 | 9.82E-01 | 2.63E-01 | 8.23E-01 | 7.73E-01 | 4.04E-01 | 1.39E-01 |
| CRnight 12h 6mo. r1 | 1.94E-05 | 1.37E-02 | 5.69E-01 | 1.68E-01 | 1.03E-01 | 5.90E-01 | 2.34E-01 | 7.45E-01 | 2.33E-02 | 4.88E-04 |
| CRnight 12h 6mo. r2 | 3.74E-03 | 1.80E-03 | 1.09E-01 | 1.52E-01 | 4.05E-01 | 2.68E-01 | 5.41E-01 | 4.58E-01 | 2.02E-01 | 4.06E-03 |
| CRnight 12h 19mo. r1 | 7.88E-04 | 6.61E-04 | 8.19E-01 | 6.75E-01 | 1.96E-01 | 2.50E-01 | 6.88E-01 | 5.81E-03 | 6.98E-01 | 4.42E-03 |
| CRnight 12h 19mo. r2 | 4.49E-04 | 1.46E-03 | 1.51E-01 | 8.27E-02 | 4.01E-01 | 2.09E-01 | 2.60E-01 | 7.98E-02 | 4.25E-01 | 1.48E-02 |
| CRspread 6mo. r1 | 3.24E-02 | 1.84E-01 | 1.85E-01 | 7.14E-01 | 4.05E-01 | 2.75E-02 | 6.25E-01 | 9.71E-01 | 1.56E-02 | 1.73E-02 |
| CRspread 6mo. r2 | 4.96E-02 | 3.87E-02 | 5.86E-01 | 7.28E-01 | 4.05E-01 | 8.53E-01 | 6.73E-02 | 5.06E-01 | 5.03E-01 | 1.20E-01 |
| CRspread 19mo. r1 | 3.78E-04 | 2.23E-01 | 5.81E-01 | 7.20E-01 | 4.14E-01 | 3.28E-02 | 8.76E-01 | 3.74E-02 | 7.43E-01 | 3.89E-01 |
| CRspread 19mo. r2 | 2.31E-05 | 1.87E-02 | 1.86E-01 | 4.89E-01 | 8.87E-01 | 5.99E-02 | 5.98E-01 | 3.67E-01 | 5.04E-01 | 6.68E-02 |

**Table S11:** An assesment of the periodicity or not of the cell cycle genes across the conditions studied in Acosta-Rodríguez *et al.* (*10*). The p-values from Cosinor for the timeseries from each of the genes in each feeding condition.

#### Genes identified in Acosta-Rodríguez *et al.*

**Ageing related genes.** In their study (*10*) Acosta-Rodríguez *et al.* found that 29% of the liver transcriptome is susceptible to aging-related changes under any condition tested. Among those genes they highlighted *Adora1, Serpine1, Themis, Spon2, Zcchc11, Gnmt, Agxt, Hmgcr, Got, Lepr, Lpin1, Pfkfb3, Scap, Hsd17b2, Hsd3b5, Cyp* and *Slc,*.

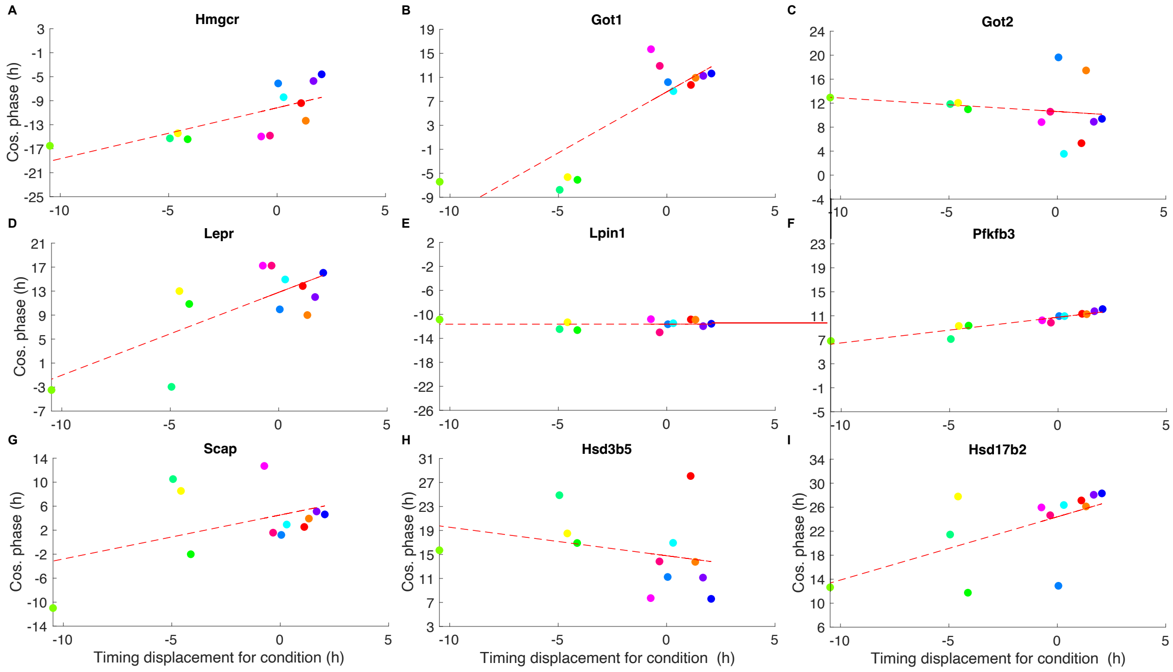


J

| | Hmgcr | Got1 | Got2 | Lepr | Lpin1 | Pfkfb3 | Scap | Hsd17b2 | Hsd3b5 |
|---|---|---|---|---|---|---|---|---|---|
| P-value | 7.58E-03 | 1.58E-03 | 4.29E-01 | 8.85E-02 | 9.84E-01 | 7.33E-06 | 2.57E-02 | 3.60E-01 | 3.69E-01 |
| R-squared | 0.53 | 0.65 | 0.06 | 0.26 | 0.00 | 0.88 | 0.41 | 0.08 | 0.08 |

K

AL 6 mo. · AL 19 mo. · CR day 2h 6 mo. · CR day 2h 19 mo. · CR day 12h 6 mo. · CR day 12h 19 mo. · CR night 2h 6 mo. · CR night 2h 19 mo. · CR night 12h 6 mo. · CR night 12h 19 mo. · CR spread 6 mo. · CR spread 19 mo.

**Figure S26: Ageing genes 1** PCPs for a subset of the ageing genes from (*10*). The Zhang *et al.* data was used for training, timecourse-matched normalisation was used for test data and $l_{\text{thresh}}$ = -8.

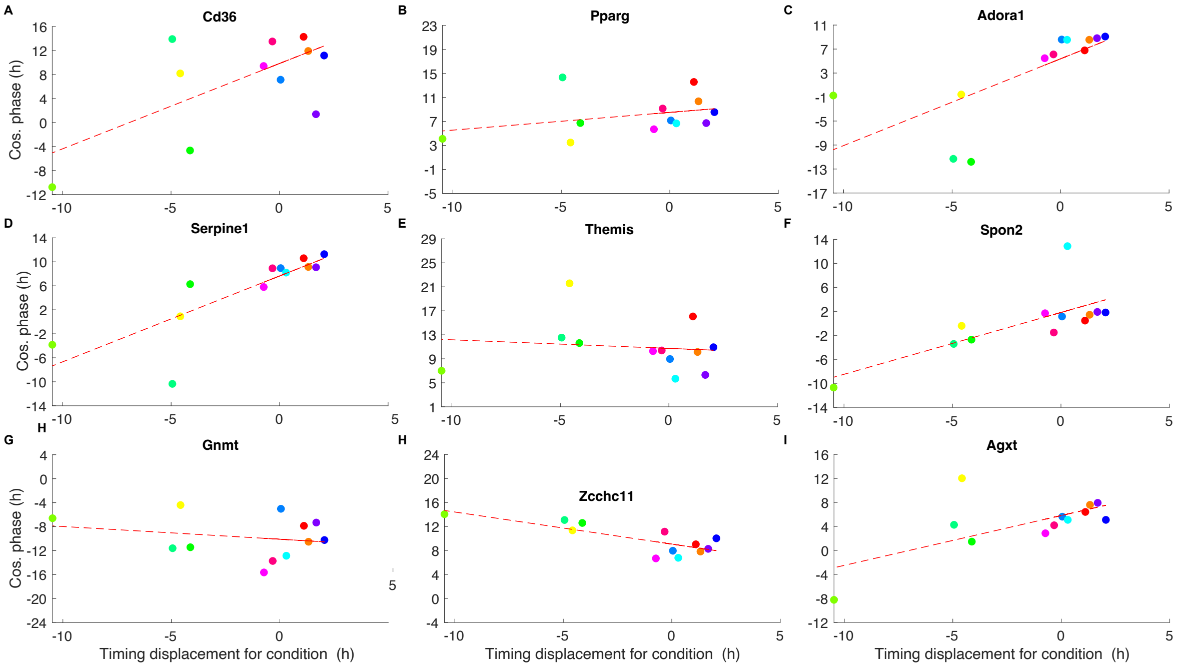


J

| | Cd36 | Pparg | Adora1 | Serpine1 | Themis | Spon2 | Zcchc11 | Gnmt | Agxt |
|---|---|---|---|---|---|---|---|---|---|
| p-value | 3.51E-01 | 2.99E-01 | 9.12E-04 | 1.28E-03 | 7.06E-01 | 3.38E-02 | 2.08E-03 | 4.83E-01 | 7.76E-02 |
| r-squared | 0.09 | 0.11 | 0.68 | 0.66 | 0.01 | 0.38 | 0.63 | 0.05 | 0.28 |

K

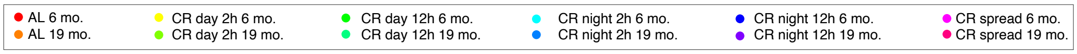


**Figure S27: Ageing genes 2** PCPs for the remaining ageing genes from (*10*). The Zhang *et al.* data was used for training, timecourse-matched normalisation was used for test data and $l_{\text{thresh}}$ = -8.

**Fasting genes** They also identified fasting genes (i.e. that were differentially expressed only in AL and CR-spread but remained constant in all the other four CR groups with some degree of fasting) i.e. *Col12a1, Plag1, Chaf1a* and *Hal*.

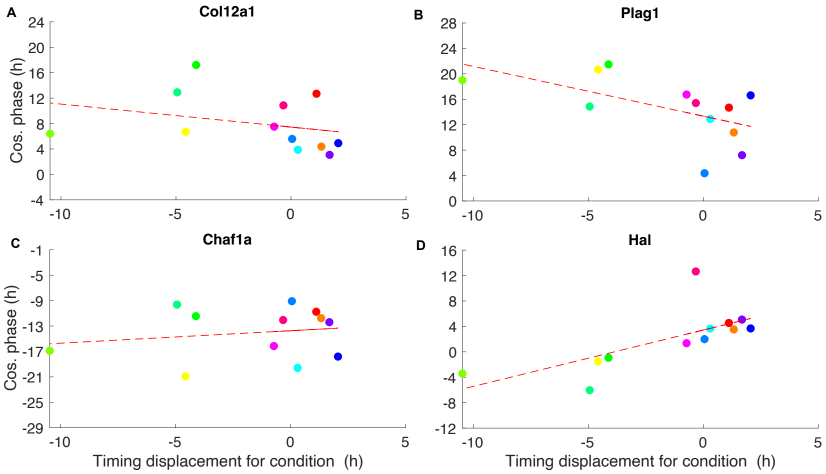


E

| Row | Col12a1 | Plag1 | Chaf1a | Hal |
|---|---|---|---|---|
| p-value | 3.32E-01 | 5.29E-02 | 5.71E-01 | 1.26E-02 |
| r-squared | 0.09 | 0.33 | 0.03 | 0.48 |

F

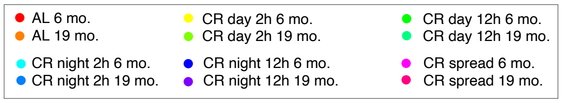


**Figure S28: Fasting genes A-D.** PCPs for the fasting genes from (*10*). The Zhang *et al.* data was used for training, timecourse-matched normalisation was used for test data and $l_{\text{thresh}}$ = -8.

**Timing related genes.** To study the beneficial effect of feeding time they identified timing related genes. These were defined as genes that maintained similar levels at young and old ages only in the CR-night fed groups with the longest life spans but were differentially expressed in the AL, CR-spread, and CR-day fed groups. None of these genes appeared to change with the clock in a coherent way.

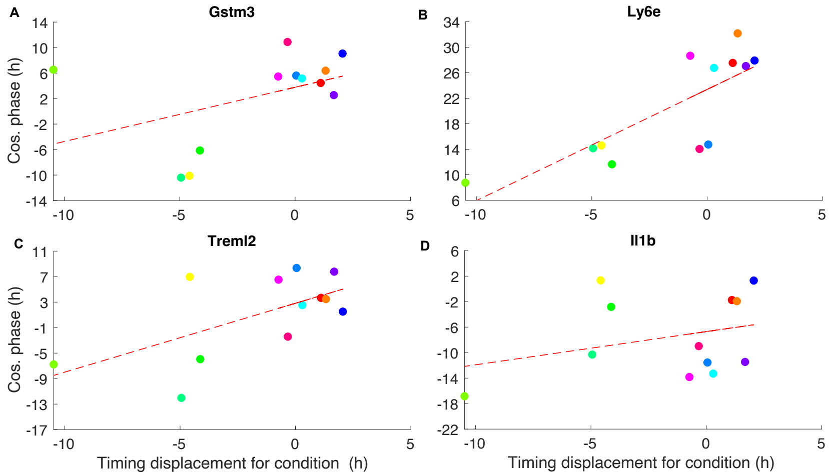


E

| | Gstm3 | Ly6e | Treml2 | Il1b |
|---|---|---|---|---|
| p-value | 1.52E-01 | 1.36E-01 | 4.62E-03 | 9.45E-01 |
| r-squared | 0.19 | 0.21 | 0.57 | 0.00 |

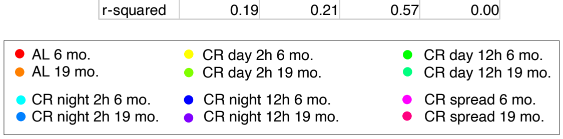


**Figure S29: Timing-related genes.** PCPs for timing-related genes from (*10*) that are not in the the TimeTeller training REP. The Zhang *et al.* data was used for training, timecourse-matched normalisation was used for test data and $l_{\text{thresh}}$ = -8.

**Genes affecting circadian cycling.** To investigate the effects on circadian cycling, using strict criteria for cyclicality, they observed that circadian cycling genes were both lost and gained with age. Three of the four circadian clock genes, *Arntl, Nr1d1, Per1* and *Per2* had a lower amplitude in old mice as did the metabolic pathway genes, *Gys2* and *Pck1*. Of these, we observe that *Arntl, Nr1d1* and *Per2* maintain strong coherence and *Pck1* maintains moderate coherence.

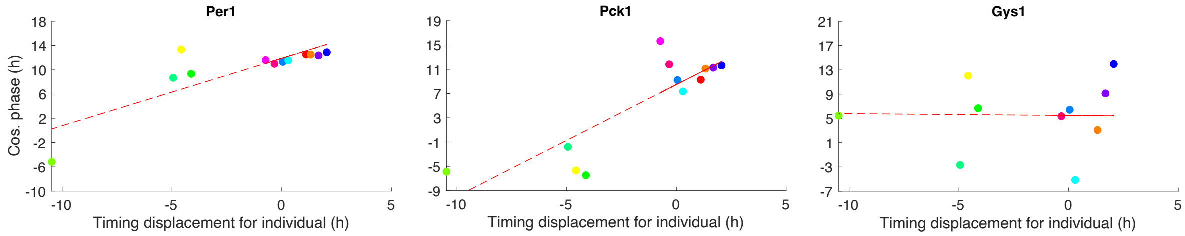


| | Arntl | Nr1d1 | Per1 | Per2 | Gys1 | Pck1 |
|---|---|---|---|---|---|---|
| p-value | 7.68E-12 | 6.91E-04 | 1.76E-01 | 3.46E-09 | 7.97E-01 | 3.15E-03 |
| r-squared | 0.99 | 0.70 | 0.17 | 0.97 | 0.01 | 0.60 |

**Figure S30: Genes affecting circadian cycling.** PCPs for three of the the genes affecting circadian cycling from (*10*). The genes in this class which are not shown (*Nr1d1*, *Per2* and *Arntl*) are part of the REP and are shown in Fig. 5 of **I**. The Zhang *et al.* data was used for training, timecourse-matched normalisation was used for test data and $l_{\text{thresh}}$ = -8.

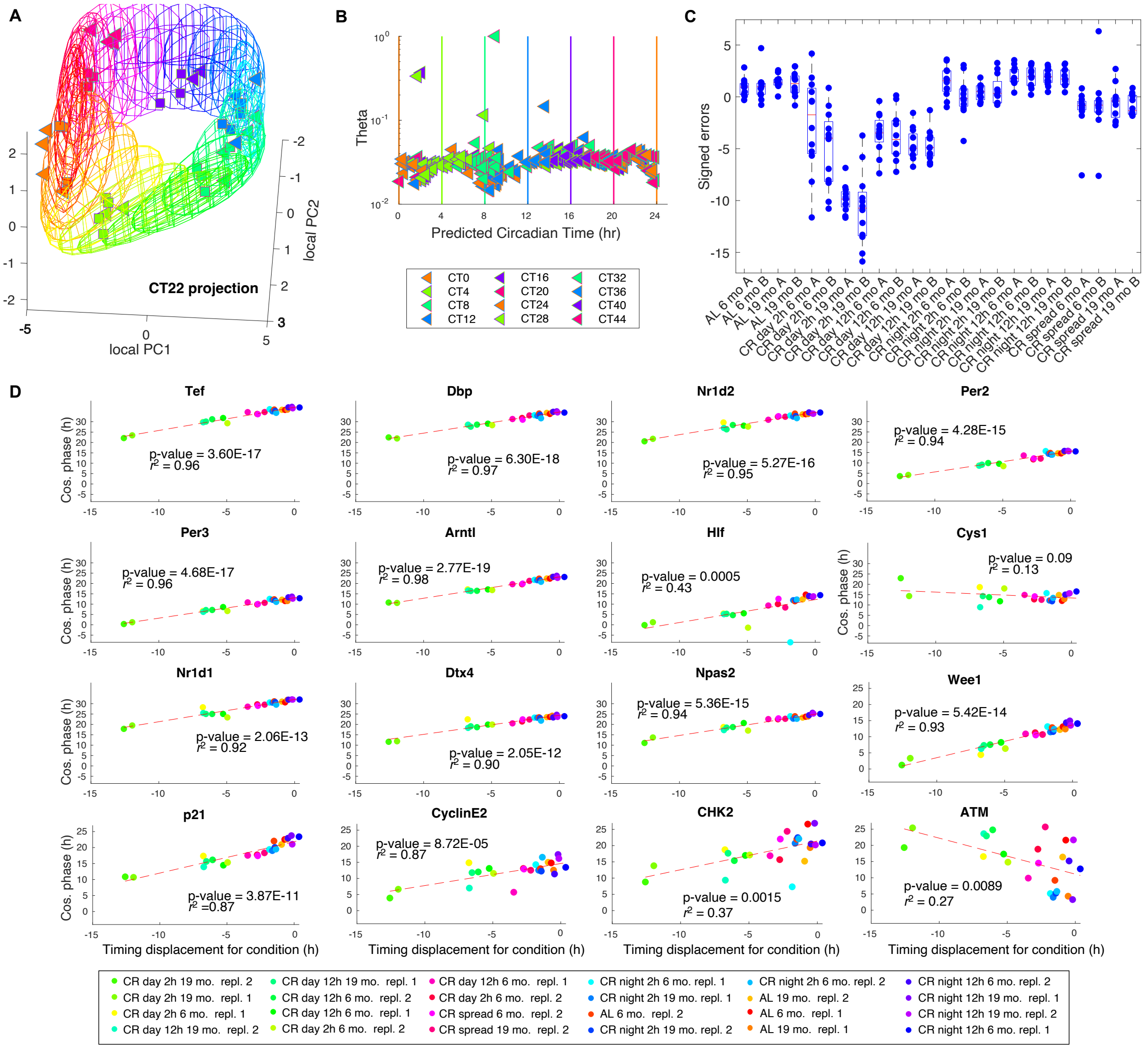


**Figure S31: Analysis of the Acosta-Rodríguez *et al.* data using a different value of $l_{\text{thresh}}$.** This uses $l_{\text{thresh}} = -12$ to check consistency with the results for $l_{\text{thresh}} = -8$. It also treats the two replicates for each age×feeding condition separately so that the consistency across replicates can be seen. **A.** Visualisation of the Acosta-Rodríguez *et al.* data (triangles) showing how this data sits in the Zhang *et al.* training data (squares). Timing is given by colour. **B.** Plots of the $\Theta$ value against the estimated time $T$. The vertical lines show the true time with colours indicating the sampling time as shown in the various legends. **C.** Box plots of the timing error for the different conditions. The mean value of each box plot gives the timing displacement for the condition. Replicates are treated separately so that the coherence of the results for the replicates can be assessed. **D.** PDP plots in which the timing displacement for the conditions is linearly regressed against the gene phase of each gene in the rhythmic profile used for training. For the genes in the REP except *Cys1* there is a strong linear relationship between the gene phases measured by Cosinor and the timing displacement as shown by small p-values (denoting a definite positive slope) and the large values of $r^2$. For the cell cycle genes *Wee1*, *p21* and *CyclinE2* there is a reasonable linear relationship but for *CHK2* and *ATM* this is not the case.

#### S9.12 Mure *et al.* data

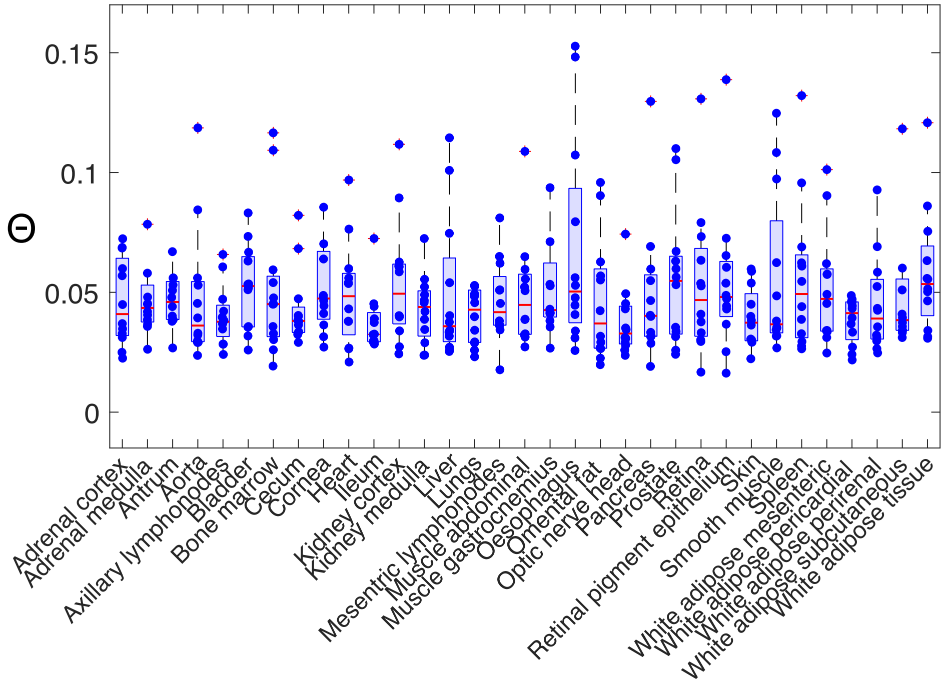


**Figure S32:** This shows the $\Theta$ values for the 33 tissues studied. The following tissues have 1 high outlier which is not shown: cornia, adrenal med., white adipose, lungs, muscle gast. . Only the central 18 tissues were used for training and $l_{\text{thresh}} = -12$.

| Gene | Arntl | Ciart | Nr1d2 | Nr1d1 | Tef | Per3 | Npas2 | Dbp | Per1 | RorC | Cry1 | Per2 | Cry2 |
|---|---|---|---|---|---|---|---|---|---|---|---|---|---|
| P-value | 4.4E-7 | 3.0E-8 | 3.4E-3 | 3.0E-5 | 3.0E-4 | 1.7E-3 | 4.6E-02 | 5.2E-7 | 1.9E-3 | 6.1E-1 | 1.9E-2 | 1.9E-4 | 5.2E-1 |
| $R^2$ | 0.57 | 0.63 | 0.25 | 0.43 | 0.35 | 0.28 | 0.12 | 0.56 | 0.27 | 0.01 | 0.17 | 0.37 | 0.01 |

**Table S12: Mure *et al.* data** The p-value and $r^2$ values for the linear regression of the gene phase of each gene shown in the plots against the timing displacements of the the 33 tissues for the Mure *et al.* data. TimeTeller has been trained on the central tissues only and $l_{\text{thresh}}$ = -12.

#### S9.13 Analysing stopped clocks

In addition to considering the *Nr1d1*KO which is associated with a partially functional clock, we also considered three examples where the clock is largely stopped and TimeTeller highlights some interesting facts about these. Both Weger *et al.* (*6*) and Yeung *et al.* (*23*) investigated the effect of knocking out *Arntl* in liver. For both datasets, the TimeTeller visualisation clearly shows clustering to CT20-24 for the KO samples as previously demonstrated by Hughey *et al.* (*22*) using ZeitZeiger.

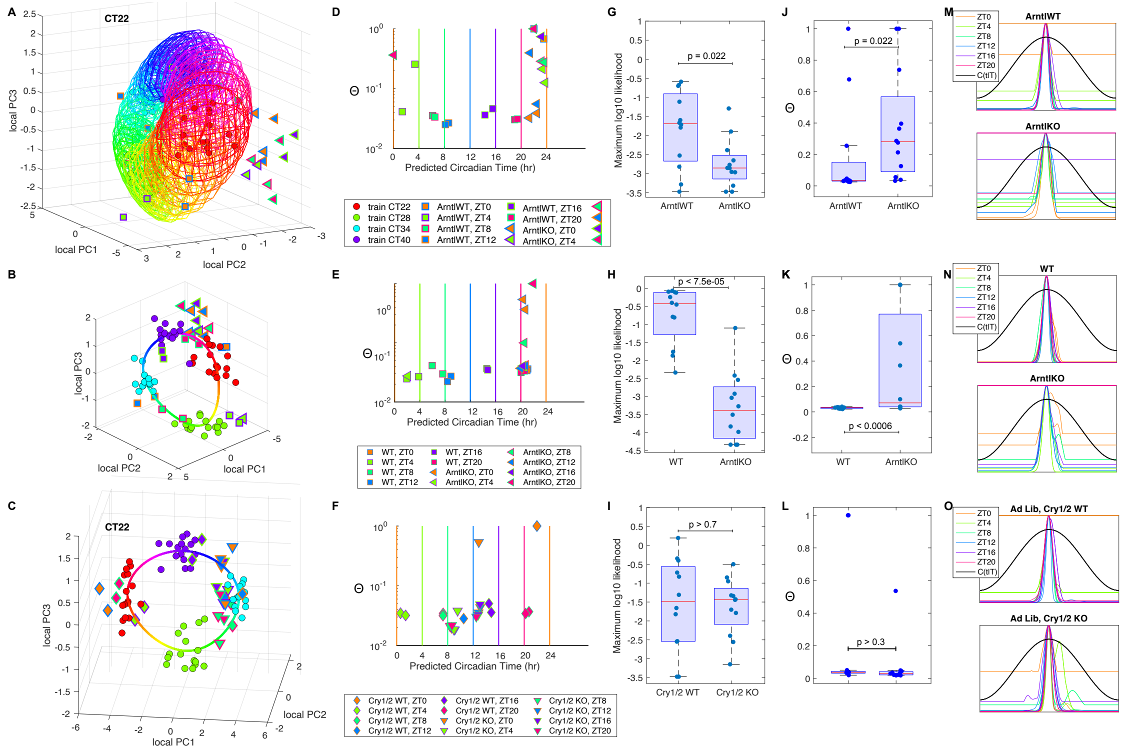


**Figure S33: Visualisation and quantitative analysis of *Arntl* and *Cr1/Cry2* knockouts.** Row 1: Weger *et al. Arntl*; row 2: Yeung *et al. Arntl*; row 3: Weger *et al. Cr1/Cry2*. All shown p-values are from the Wilcoxon Rank-Sum test. **A, B & C.** The visualisation of the data for each dataset compared to the training data. In each case the clustering of the KO data about a particular timepoint is evident. **D, E & F.** Plots of the $\Theta$ value against the estimated time $T$. The vertical lines show the true time with colours indicating the sampling time as shown in the various legends. These plots show the clustering of the KO clocks around a specific time of the day. **G, H & I.** Boxplots of the maximum likelihood $(\mathrm{ML})$ values for the three datasets. From G and H we see that for both the Weger *et al.* and Yeung *et al.* dataset involving *Arntl* there is a significant decrease in the $\mathrm{ML}$ for the KOs suggesting the presence of lowML type dysfunction for these where the KO data has moved away from the training clock mean trajectory. Moreover, from G and H, using the method explained in Sect. S2.3, these likelihoods suggest choosing $l_{\text{thresh}} = -8$ for the Weger *et al. Arntl* data and $l_{\text{thresh}} = -10$ for the Yeung *et al.* data. This is because of the lower likelihoods in the Yeung *et al.* KO data. The Weger *et al. Cry1/Cry2* data is different in that when $l_{\text{thresh}} = -8$, the $\mathrm{ML}$ values shown in I are very similar between WT and KO, and this result holds true for $6 \leq l_{\text{thresh}} \leq 12$. **J, K & L.** Boxplots of the $\Theta$ values for the three datasets. For the both sets of *Arntl* KO data the $\Theta$ values are significantly higher than for the WT data. However, for the *Cry1/Cry2* data there is no such difference. **M, N & O.** Centred LRFs for the three datasets. **M & N.** For the two *Arntl* KO datasets the only significant contribution to $\Theta$ is coming from flat parts of the LRF that are above $C(t|T)$ and therefore we conclude that the dysfunction is primarily of lowML type.

In addition to the visualisation we see that although the $\Theta$ values for the KO appear generally larger than the WT ones, they are not significantly so. There is a significant difference in the maximum likelihood values for the WT and KO groups for the Weger *et al.* data but not for the Yeung *et al.* data. This provides quantitative evidence that the clock in the liver became frozen at a particular state very close to a wild-type clock state and corresponding to a relatively narrow time range.

Similar results are found for the data from Weger *et al.* (*6*) studying the *Cry1/Cry2* double KO in liver. There is no significant difference between the Θs and maximum likelihoods for the WT and KO but the timings for the KO sample are clustered around CT8-12. Previously Hughey *et al.* (*24*) demonstrated using ZeitZeiger that *Cry1/2* KO microarray samples from (*25*) also displayed time predictions of CT8-13. Our results support the conclusion that *Cry1/2* KO breaks the molecular oscillator such that the expression of E-box driven genes such as *Per2* and *Dbp* remains constantly high, as the *Per/Cry* repressor complex cannot form to repress E-box driven gene expression. Since E-box driven genes are at their highest expression at CT8-12, this is the timepoint that best represents the gene expression that results from *Cry1/2* KO. As for the *Arntl* KO we suspect the clock has been frozen because a saddle-node SNIC bifurcation has occured. A critic might comment that since in such a KO the e-box activators are missing, therefore, the clock should look like the maximally repressed state. But this does not explain why the observed state is so close to the WT limit cycle.

### S10 Precision assessment without time stamps: Feng *et al.* and Cadenas *et al.*

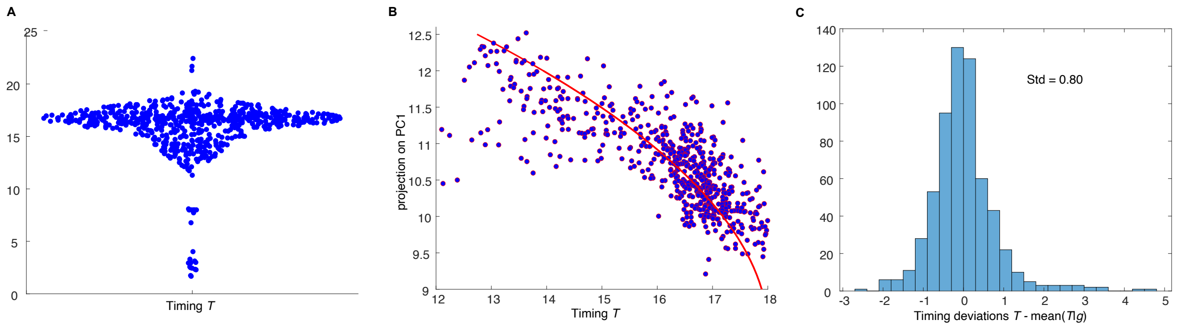


**Figure S34: A.** Swarm chart showing the TimeTeller timings $T$ of the Cadenas *et al.* data. **B.** A scatter plot of the projection $\tilde{g}$ of the data REVs $g$ onto the first PC (vertical axis) against the TimeTeller timing $T$. The smooth red curve approximates the means of $P(T|\tilde{g})$. **C.** The distribution of the deviations i.e. the horizontal difference between each data point $(T, \tilde{g})$ and the mean of $P(T|\tilde{g})$. The standard deviation of this distribution is shown.

In this section we explain how we calculate an upper bound on the variance of $P(T|g)$ from data where $T$ is the TimeTeller estimated time and $g$ is the gene expression vector or REV. This approach can also be used to get MAEs or MdAEs for this distribution. $P(T|g)$ gives the distribution of the TimeTeller estimate conditional on the gene expression state $g$. The methods currently used to quantify the precision of phase estimation algorithms focus on $P(T|t)$ where $t$ is the sample time. This restricts such assessments to data that is time stamped. Moreover, for an individual sample the value $T - t$ is strongly biased by any chronotype for the individual, tissue or condition associated with the sample. For our human data we saw a range from -3.5h to 2.5h. Therefore, one can argue that it is more natural to instead try to assess the variance of $P(T|g)$ instead. We now describe in more detail how we do this.

We assume that we have a dataset and a determination of $T$ for each data point. This might be the straightforward assessment of $T$ by TimeTeller or it might be some adjusted value of $T$ using TimeTeller such as the use of second peaks as done for the Feng *et al.* data. In addition, we will have the value of the REV $g$ for each datapoint. We use the Feng *et al.* (*3*) data and that discussed in Cadenas *et al.* (*14*) as examples.

We carry out PCA analysis as described in Sect. S7.1 where we take for the vectors $v_j$ the REVs $g$ in our dataset. We then project the REVs onto the first principal component $\mathbf{b}_1$ to get projected values $\tilde{g} = g \cdot \mathbf{b}_1$. From this we can produce a scatter plot as in Figs. 3I & S34B where the projected value $\tilde{g}$ of $g$ is plotted against the corresponding value of $T$. We then use kernel based smoothing or related methods to obtain a smooth curve $T = m(\tilde{g})$ giving the mean of the $T$ values as a function of $\tilde{g}$ (Figs. 3I & S34B). For the Feng *et al.* data the kernels used are normal with variance 0.5. Then for each data point (with timing $T_i$ and REV $g_i$) we calculate the quantity $T_i - m(\tilde{g}_i))$. These are the timing deviations. The resulting distributions are shown in (Figs. 3J & S34C). We calculate the standard deviation (std) of these (overall std) and also check that it does not deviate much from the std found of we restrict to a subinterval of $\tilde{g}$ values with a reasonable number of data values in it (local std). This gives an estimate of the std of the distribution $P(T|\tilde{g})$ which is an upper bound of the std of $P(T|g)$.

For the Cadenas *et al.* data the analysis has been restricted to the data between $T = 10$ and $T = 20$.

### S11 Supplementary information about synthetic datasets: 1. Local PCs versus global PCs, 2. Θ dependence on the efficiency of a *Bmal1* knockdown

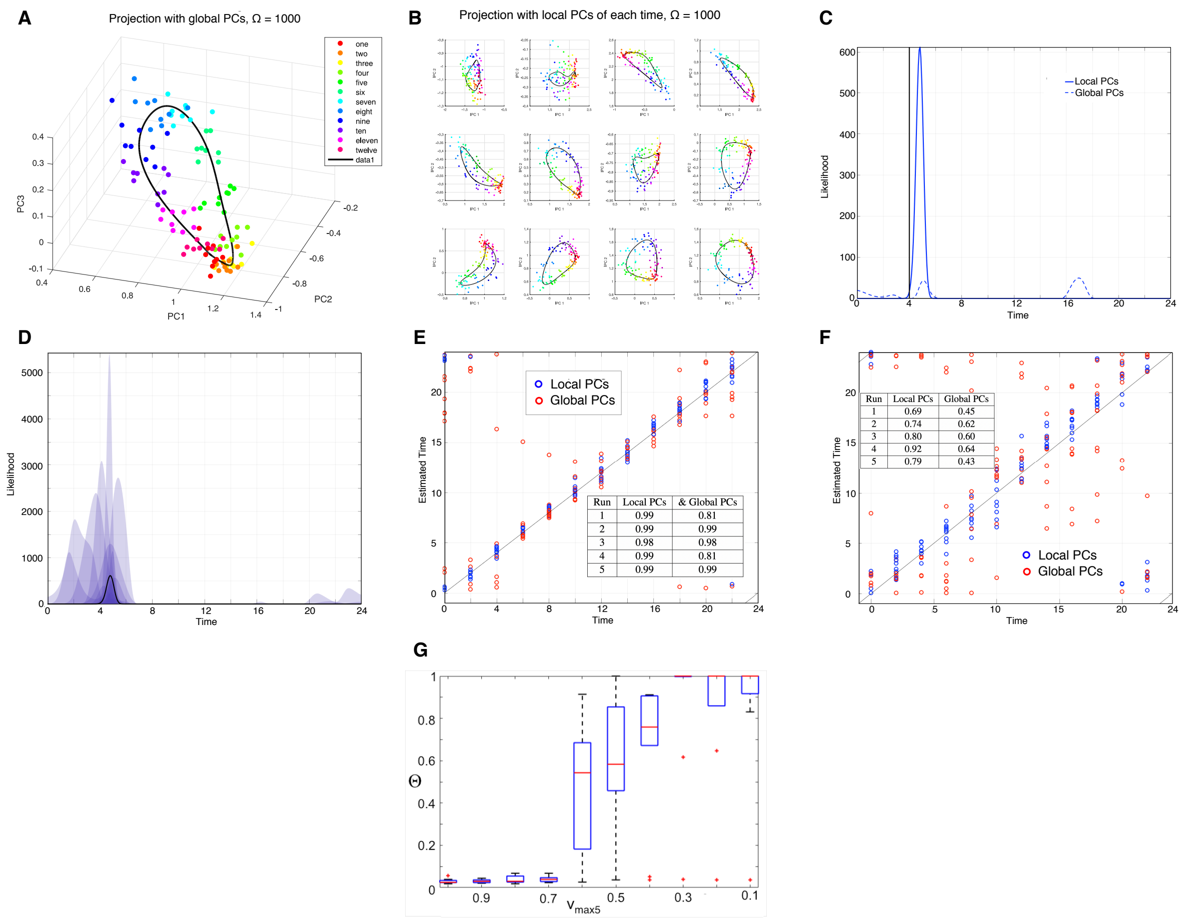

**Figure S35: A-F. The construction of the TimeTeller probability model and likelihood function uses projections calculated locally (local PCs) which are then combined.** We compared the effectiveness of this local approach to the simpler one using a single principal component projection of all the training data (global PCs). To do this we used simulated data from the stochastic Relogio model (Note S12.1) using systems sizes $\Omega = 1000$ and $\Omega = 100$ and 12 time points around the day. The latter system size produces particularly noisy data. **(A)** A global PC projection of the data when $\Omega = 1000$. **(B)** The 12 local PC projections. **(C)** Example likelihood curves for the same data point for global and local PCs. The global PC likelihood is shown with the dashed line and the local PC likelihood is shown in bold, for the same training and test data. The black vertical line shows the true time $T$ of the sample. Notice the improvement in accuracy when using local PCs and the MLE peak at $T + 12$ for the global PCs. **(D)** Plot showing how individual local PC likelihoods are averaged to produce the combined likelihood. The likelihood curves for the local PCs are (geometrically) averaged, to get the overall likelihood (Note S2.2). This helps to overcome problems with symmetry as described in Fig. S1 and seen in (C). **(E)** Scatter plot showing estimated versus real time for low noise ($\Omega = 1000$) simulations, using local and global PCA. Correlation is generally very high, with some obvious symmetry issues with the global PC estimations. The red point at (8, 13) is the estimate resulting from the dotted likelihood in (C). The inset table shows $R^2$ values for linear fit to $x = y$ of estimated versus real time, for $\Omega = 1000$ simulated data. Values are generally accurate due to low noise. $R^2$s are similar except for runs 1 and 4, where the accuracy of the local PC method is higher than for global PC method. **(F)** As (E) but for the high noise $\Omega = 100$ simulations. The training and test data are identical. From (E) and (F) and the inset tables we observe that the method using local PCs is more accurate than the method using global PCs. The inset table is as for (E) but for $\Omega = 100$. Values vary due to high noise in the simulated data. $R^2$s for local PCs are significantly higher than for global PC. **(G) $\Theta$ dependence on the efficiency of a *Bmal1* knockdown.** We investigated how $\Theta$ increases as we increase the efficiency of a *Bmal1* knockdown using synthetic data from the stochastic Relogio model with intermediate systems size $\Omega = 500$, giving a moderate level of stochasticity. *Bmal1* is crucial for the proper functioning of the circadian clock (*26*). The knockdown of *Bmal1* is modelled by decreasing the rate $V_{max5}$ of *Bmal1* transcription from the WT value of 1 to the full knock-down value of 0.1, in steps of 0.1. For each of ten stochastic simulations, 10 equally spaced time-points were extracted as the test sets. The box plots show that there is an (almost) monotonic relationship between $\Theta$ and the change in the $V_{max5}$ parameter and that, as $V_{max5}$ falls below 0.7, the functioning of the clock is severely affected.

### S12 Stochastic modelling

There have been a number of stochastic models of the circadian clocks (e.g., (*27–29*)). In particular, an early detailed stochastic mammalian model of circadian rhythms was published by Forger *et al.* (*30*), who adapted his previous 74 equation ODE model. Instead of attempting to replicate this existing stochastic model of mammalian circadian rhythms, we have created a stochastic version of the Relogio model (*31*) with the adaptation based on methods in (*27*). This stochastic model incorporates a system size parameter, $\Omega$. We consider it has units L/nM and a value in the range 100-1000 based on typical cell size estimates (*32*). The individual trajectories were calculated in MATLAB, using an implementation of the Gillespie algorithm or the pc-LNA (*32*).

#### S12.1 The stochastic Relogio model

The stochastic Relogio model has 44 reactions using 19 variables and 72 parameters, where parameters with units of concentration are scaled with system size parameter accordingly. As $\Omega \to \infty$, the stochastic solution converges to the ODE solutions. The 44 reaction rates for the stochastic Relogio model are shown in Note S12.1. In order to generate enough trajectories to estimate a distribution in a reasonable time the pc-LNA was used (*32*).

##### ODE

$$
\begin{aligned}
\frac{\mathrm{d}CCK/BML}{\mathrm{d}t} &= kf_{x1}BML_N - kd_{x1}CCK/BML - d_{x1}CCK/BML\\
\frac{\mathrm{d}P_N^*/C_N}{\mathrm{d}t} &= ki_{z4}P_C^*/C_C - ke_{x2}P_N^*/C_N - d_{x2}P_N^*/C_N\\
\frac{\mathrm{d}P_N/C_N}{\mathrm{d}t} &= ki_{z5}P_C/C_C - ke_{x3}P_N/C_N - d_{x3}P_N/C_N\\
\frac{\mathrm{d}BML_N}{\mathrm{d}t} &= ki_{z8}BML_c + kd_{x1}CCK/BML - kf_{x1}BML_N - d_{x7}BML_N\\
\frac{\mathrm{d}Reverb_n}{\mathrm{d}t} &= ki_{z6}RVRB_c - d_{x5}Reverb_n\\
\frac{\mathrm{d}Ror_n}{\mathrm{d}t} &= ki_{z7}ROR_c - d_{x6}Ror_n\\
\frac{\mathrm{d}Per}{\mathrm{d}t} &= V_{1max}\left(\frac{(1+a\left(\frac{CCK/BML}{k_{t1}}\right)^b}{1+\left(\frac{P_N^*/C_N+P_N/C_N}{k_{i1}}\right)^c\left(\frac{CCK/BML}{k_{t1}}\right)^b+\left(\frac{CCK/BML}{k_{t1}}\right)^b}\right) - d_{y1}Per\\
\frac{\mathrm{d}Cry}{\mathrm{d}t} &= V_{2max}\left(\frac{1+d\left(\frac{CCK/BML}{k_{t2}}\right)^e}{1+\left(\frac{P_N^*/C_N+P_N/C_N}{k_{i2}}\right)^f\left(\frac{CCK/BML}{k_{t2}}\right)^e+\left(\frac{CCK/BML}{k_{t2}}\right)^e}\frac{1}{1+\left(\frac{Reverb_n}{k_{i21}}\right)^{f_1}}\right) - d_{y2}Cry\\
\frac{\mathrm{d}Reverb}{\mathrm{d}t} &= V_{3max}\left(\frac{1+g\left(\frac{CCK/BML}{k_{t3}}\right)^v}{1+\left(\frac{P_N^*/C_N+P_N/C_N}{k_{i3}}\right)^w\left(\frac{CCK/BML}{k_{t3}}\right)^v+\left(\frac{CCK/BML}{k_{t3}}\right)^v}\right) - d_{y3}Reverb\\
\frac{\mathrm{d}Ror}{\mathrm{d}t} &= V_{4max}\left(\frac{1+h\left(\frac{CCK/BML}{k_{t4}}\right)^p)}{1+\left(\frac{P_N^*/C_N+P_N/C_N}{k_{i4}}\right)^q\left(\frac{CCK/BML}{k_{t4}}\right)^p+\left(\frac{CCK/BML}{k_{t4}}\right)^p}\right) - d_{y4}Ror\\
\frac{\mathrm{d}Bmal}{\mathrm{d}t} &= V_{5max}\left(\frac{1+i\left(\frac{Ror_n}{k_{t5}}\right)^n}{1+\left(\frac{Reverb_n}{k_{i5}}\right)^m+\left(\frac{Ror_n}{k_{t5}}\right)^n}\right) - d_{y5}Bmal\\
\frac{\mathrm{d}Cry_C}{\mathrm{d}t} &= k_{p2}(Cry+y_{20}) + kd_{z4}P_C^*/C_C + kd_{z5}P_C/C_C - kf_{z5}Cry_CPer_C - kf_{z4}Cry_CPer_C^* - d_{z1}Cry_C\\
\frac{\mathrm{d}Per_C}{\mathrm{d}t} &= k_{p1}(Per+y_{10}) + kd_{z5}P_C/C_C + kd_{phz3}Per_C^* - kf_{z5}Per_CCry_C - kph_{z2}Per_C - d_{z2}Per_C\\
\frac{\mathrm{d}P_C^*/C_C}{\mathrm{d}t} &= kf_{z4}Cry_CPer_C^* + ke_{x2}P_N^*/C_N - ki_{z4}P_C^*/C_C - kd_{z4}P_C^*/C_C - d_{z4}P_C^*/C_C
\end{aligned}
\tag{5}
$$

$$\frac{\mathrm{d}Per*_C}{\mathrm{d}t} = kph_{z2}Per_C + kd_{z4}P^*_C/C_C - kd_{phz3}Per^*_C - kf_{z4}Per^*_C Cry_C - d_{z3}Per^*_C$$

$$\frac{\mathrm{d}P_C/C_C}{\mathrm{d}t} = kf_{z5}Cry_C Per_C + ke_{x3}P_N/C_N - ki_{z5}P_C/C_C - kd_{z5}P_C/C_C - d_{z5}P_C/C_C$$

$$\frac{\mathrm{d}RVRB_c}{\mathrm{d}t} = k_{p3}(Reverb + y_{30}) - ki_{z6}RVRB_c - d_{z6}RVRB_c$$

$$\frac{\mathrm{d}ROR_c}{\mathrm{d}t} = k_{p4}(Ror + y_{40}) - ki_{z7}ROR_c - d_{z7}ROR_c$$

$$\frac{\mathrm{d}BML_c}{\mathrm{d}t} = k_{p5}(Bmal + y_{50}) - ki_{z8}BML_c - d_{z8}BML_c$$

**Stochastic model.**

19 state variables. 44 reactions. 72 parameters.

$$a(1) = kf_{x1}BML_N \quad a(2) = kd_{x1}CCK/BML \quad a(3) = d_{x1}CCK/BML$$
$$a(4) = ki_{z4}P^*_C/C_C \quad a(5) = ke_{x2}P^*_N/C_N \quad a(6) = d_{x2}P^*_N/C_N$$
$$a(7) = ki_{z5}P_C/C_C \quad a(8) = ke_{x3}P_N/C_N \quad a(9) = d_{x3}P_N/C_N$$
$$a(10) = ki_{z8}BML_c \quad a(11) = d_{x7}BML_N \quad a(12) = ki_{z6}RVRB_c$$
$$a(13) = ki_{z7}ROR_c \quad a(15) = d_{x6}Ror_n \tag{6}$$

$$a(16) = V_{1max}\left(\frac{(1 + a\left(\frac{CCK/BML}{k_{t1}}\right)^b}{1 + \left(\frac{P^*_N/C_N + P_N/C_N}{k_{i1}}\right)^c\left(\frac{CCK/BML}{k_{t1}}\right)^b + \left(\frac{CCK/BML}{k_{t1}}\right)^b}\right)$$
$$a(17) = d_{y1}Per$$
$$a(18) = V_{2max}\left(\frac{1 + d\left(\frac{CCK/BML}{k_{t2}}\right)^e}{1 + \left(\frac{P^*_N/C_N + P_N/C_N}{k_{i2}}\right)^f\left(\frac{CCK/BML}{k_{t2}}\right)^e + \left(\frac{CCK/BML}{k_{t2}}\right)^e}\frac{1}{1 + \left(\frac{Reverb_n}{k_{i21}}\right)^{f_1}}\right)$$
$$a(19) = d_{y2}Cry$$
$$a(20) = V_{3max}\left(\frac{1 + g\left(\frac{CCK/BML}{k_{t3}}\right)^v}{1 + \left(\frac{P^*_N/C_N + P_N/C_N}{k_{i3}}\right)^w\left(\frac{CCK/BML}{k_{t3}}\right)^v + \left(\frac{CCK/BML}{k_{t3}}\right)^v}\right)$$
$$a(21) = d_{y3}Reverb$$
$$a(22) = V_{4max}\left(\frac{1 + h\left(\frac{CCK/BML}{k_{t4}}\right)^p)}{1 + \left(\frac{P^*_N/C_N + P_N/C_N}{k_{i4}}\right)^q\left(\frac{CCK/BML}{k_{t4}}\right)^p + \left(\frac{CCK/BML}{k_{t4}}\right)^p}\right) \tag{7}$$

$$a(23) = d_{y4}Ror$$
$$a(24) = V_{5max}\left(\frac{1 + i\left(\frac{Ror_n}{k_{t5}}\right)^n}{1 + \left(\frac{Reverb_n}{k_{i5}}\right)^m + \left(\frac{Ror_n}{k_{t5}}\right)^n}\right)$$

$$
\begin{aligned}
&a(25) = d_{y5}Bmal, \quad a(26) = k_{p2}Cry, \quad a(27) = kd_{z4}P_C^*/C_C, \\
&a(28) = kd_{z5}P_C/C_C, \quad a(29) = kf_{z5}Cry_C Per_C, \quad a(30) = kf_{z4}Cry_C Per_C^*, \\
&a(31) = d_{z1}Cry_C, \quad a(32) = k_{p1}Per, \quad a(33) = kd_{phz3}Per_C^*, \\
&a(34) = kph_{z2}Per_C, \quad a(35) = d_{z2}Per_C, \quad a(36) = d_{z3}P_C^*/C_C, \\
&a(37) = d_{z4}Per_C^*, \quad a(38) = d_{z5}P_C/C_C, \quad a(39) = k_{p3}Reverb, \\
&a(40) = d_{z6}RVRB_c, \quad a(41) = k_{p4}Ror, \quad a(42) = d_{z7}ROR_c, \\
&a(43) = k_{p5}Bmal, \quad a(44) = d_{z8}BML_c. \qquad (8)
\end{aligned}
$$

**Parameters**

| Parameter | Value | Description |
|---|---|---|
| $aa$ | 12 | Per |
| $d$ | 12 | Cry |
| $g$ | 5 | Rev-Erb |
| $h$ | 5 | Ror |
| $i$ | 12 | Bmal |
| $b$ | 5 | Per-activation |
| $c$ | 7 | Per-inhibition |
| $e$ | 6 | Cry-activation rate |
| $f$ | 4 | Cry-inhibition |
| $f1$ | 1 | Cry-inhibition |
| $v$ | 6 | Rev-Erb-activation |
| $w$ | 2 | Rev-Erb-inhibition |
| $p$ | 6 | Ror-activation |
| $q$ | 3 | Ror-inhibition |
| $n$ | 2 | Bmal-activation |
| $m$ | 5 | Bmal-inhibition |
| $y_{10}$ | 0*omega | Per |
| $y_{20}$ | 0*omega | Cry |
| $y_{30}$ | 0*omega | Rev-Erb |
| $y_{40}$ | 0*omega | Ror |
| $y_{50}$ | 0*omega | Bmal |
| $d_{z1}$ | 0.23 | CRYC |
| $d_{z2}$ | 0.25 | PERC |
| $d_{z3}$ | 0.6 | PERC* |
| $d_{z4}$ | 0.2 | PERC*/CRYC |
| $d_{z5}$ | 0.2 | PERC/CRYC |
| $d_{z6}$ | 0.31 | REV-ERBC |
| $d_{z7}$ | 0.3 | RORC |
| $d_{z8}$ | 0.73 | BMALC |
| $d_{y1}$ | 0.3 | Per |
| $d_{y2}$ | 0.2 | Cry |
| $d_{y3}$ | 2 | Rev-Erb |
| $d_{y4}$ | 0.2 | Ror |
| $d_{y5}$ | 1.6 | Bmal |
| $d_{x1}$ | 0.08 | CLOCK/BMAL ** -23 PD Huge Phase deriv -38 PD |
| $d_{x2}$ | 0.06 | PER*N/CRYN ** |
| $d_{x3}$ | | PERN/CRYN |
| $d_{x5}$ | 0.17 | REV-ERBN |

| | | |
|---|---|---|
| $d_{x6}$ | 0.12 | RORN ** -28 PD |
| $d_{x7}$ | 0.15 | BMALN |
| $kf_{x1}$ | 2.3 | CLOCK/BMAL-complex formation [hour-1] |
| $kd_{x1}$ | 0.01 | CLOCK/BMAL-complex dissociation [hour-1] |
| $kf_{z4}$ | 1/omega | PERC*/CRYC-complex formation [(a.u.$\times$hour)-1] |
| $kd_{z4}$ | 1 | PERC*/CRYC-complex dissociation [hour-1] |
| $kf_{z5}$ | 1/omega | PERC/CRYC-complex formation [(a.u.$\times$hour)-1] |
| $kd_{z54}$ | 1 | PERC/CRYC-complex dissociation [hour-1] |
| $ki_{z4}$ | 0.2 | PERC*/CRYC |
| $ki_{z5}$ | 0.1 | PERC/CRYC |
| $ki_{z6}$ | 0.5 | REV-ERBC |
| $ki_{z7}$ | 0.1 | RORC ** 12.9 PD |
| $ki_{z8}$ | 0.1 | BMALC ** 15.6 PD |
| $ke_{x2}$ | 0.02 | PER*N/CRYN ** -21 PD |
| $ke_{x3}$ | 0.02 | PERN/CRYN |
| $kph_{z2}$ | 2 | PERC-phosphorylation rate |
| $kd_{phz3}$ | 0.05 | PERC*-dephosphorylation rate |
| $k_{p1}$ | 0.4 | PERC |
| $k_{p2}$ | 0.26 | CRYC |
| $k_{p3}$ | 0.37 | REV-ERBC |
| $k_{p4}$ | 0.76 | RORC |
| $k_{p5}$ | 1.21 | BMALC |
| $k_{t1}$ | 3*omega | Per-activation rate |
| $k_{i1}$ | 0.9*omega | Per-inhibition rate |
| $k_{t2}$ | 2.4*omega | Cry-activation rate PLOSGEN PERTUR 5.3 TO SIM RAS |
| $k_{i2}$ | 0.7*omega | Cry-inhibition rate |
| $k_{i21}$ | 5.2*omega | Cry-inhibition rate |
| $k_{t3}$ | 2.07*omega | Rev-Erb-activation rate |
| $k_{i3}$ | 3.3*omega | Rev-Erb-inhibition rate |
| $k_{t4}$ | 0.9*omega | Ror-activation rate |
| $k_{i4}$ | 0.4*omega | Ror-inhibition rate **15.5 PD |
| $k_{t5}$ | 8.35*omega | Bmal-activation rate |
| $k_{i5}$ | 1.94*omega | Bmal-inhibition rate |
| $V_{1max}$ | 1*omega | Per |
| $V_{2max}$ | 2.92*omega | Cry |
| $V_{3max}$ | 1.9*omega | Rev-Erb |
| $V_{4max}$ | 10.9*omega | Ror |
| $V_{5max}$ | 1*omega | Bmal |

**Table S13**

### Bibliography


1. M. E. Hughes, L. DiTacchio, K. R. Hayes, C. Vollmers, S. Pulivarthy, J. E. Baggs, S. Panda, and J. B. Hogenesch, "Harmonics of circadian gene transcription in mammals," *PLoS genetics*, vol. 5, no. 4, p. e1000442, 2009.

2. J. O. Boyle, Z. H. Gümüş, A. Kacker, V. L. Choksi, J. M. Bocker, X. K. Zhou, R. K. Yantiss, D. B. Hughes, B. Du, B. L. Judson, K. Subbaramaiah, and A. J. Dannenberg, "Effects of cigarette smoke on the human oral mucosal transcriptome," *Cancer Prevention Research*, vol. 3, no. 3, pp. 266–278, 2010.

3. L. Feng, J. R. Houck, P. Lohavanichbutr, and C. Chen, "Transcriptome analysis reveals differentially expressed lncRNAs between oral squamous cell carcinoma and healthy oral mucosa.," *Oncotarget*, vol. 8, no. 19, pp. 31521–31531, 2017.

4. R. Zhang, N. F. Lahens, H. I. Ballance, M. E. Hughes, and J. B. Hogenesch, "A circadian gene expression atlas in mammals: Implications for biology and medicine," *Proceedings of the National Academy of Sciences*, pp. 2–7, oct 2014.

5. K. Kinouchi, C. Magnan, N. Ceglia, Y. Liu, M. Cervantes, N. Pastore, T. Huynh, A. Ballabio, P. Baldi, S. Masri, *et al.*, "Fasting imparts a switch to alternative daily pathways in liver and muscle," *Cell reports*, vol. 25, no. 12, pp. 3299–3314, 2018.

6. B. D. Weger, C. Gobet, F. P. David, F. Atger, E. Martin, N. E. Phillips, A. Charpagne, M. Weger, F. Naef, and F. Gachon, "Systematic analysis of differential rhythmic liver gene expression mediated by the circadian clock and feeding rhythms," *Proceedings of the National Academy of Sciences*, vol. 118, no. 3, 2021.

7. B. Fang, L. J. Everett, J. Jager, E. Briggs, S. M. Armour, D. Feng, A. Roy, Z. Gerhart-Hines, Z. Sun, and M. A. Lazar, "Circadian enhancers coordinate multiple phases of rhythmic gene transcription in vivo," *Cell*, vol. 159, no. 5, pp. 1140–1152, 2014.

8. J. L. Barclay, J. Husse, B. Bode, N. Naujokat, J. Meyer-Kovac, S. M. Schmid, H. Lehnert, and H. Oster, "Circadian desynchrony promotes metabolic disruption in a mouse model of shiftwork," *PLoS ONE*, vol. 7, no. 5, 2012.

9. G. Le Martelot, D. Canella, L. Symul, E. Migliavacca, F. Gilardi, R. Liechti, O. Martin, K. Harshman, M. Delorenzi, B. Desvergne, W. Herr, B. Deplancke, U. Schibler, J. Rougemont, N. Guex, N. Hernandez, and F. Naef, "Genome-Wide RNA Polymerase II Profiles and RNA Accumulation Reveal Kinetics of Transcription and Associated Epigenetic Changes During Diurnal Cycles," *PLoS Biology*, vol. 10, no. 11, 2012.

10. V. Acosta-Rodriguez, F. Rijo-Ferreira, M. Izumo, P. Xu, M. Wight-Carter, C. Green, and J. Takahashi, "Circadian alignment of early onset caloric restriction promotes longevity in male c57bl/6j mice.," *Science*, vol. 376, no. 6598, pp. 1192–1202, 2022.

11. K. B. Koronowski, K. Kinouchi, P.-S. Welz, J. G. Smith, V. M. Zinna, J. Shi, M. Samad, S. Chen, C. N. Magnan, J. M. Kinchen, *et al.*, "Defining the independence of the liver circadian clock," *Cell*, vol. 177, no. 6, pp. 1448–1462, 2019.

12. L. S. Mure, H. D. Le, G. Benegiamo, M. W. Chang, L. Rios, N. Jillani, M. Ngotho, T. Kariuki, O. Dkhissi-Benyahya, H. M. Cooper, *et al.*, "Diurnal transcriptome atlas of a primate across major neural and peripheral tissues," *Science*, vol. 359, no. 6381, p. eaao0318, 2018.

13. G. Bjarnason, A. Seth, Z. Wang, N. Blanas, M. Straume, and T. Martino, "Diurnal rhythms (dr) in gene expression in human oral mucosa: Implications for gender differences in toxicity, response and survival and optimal timing of targeted therapy (rx)," *Journal of Clinical Oncology*, vol. 25, no. 18_suppl, pp. 2507–2507, 2007.

14. C. Cadenas, L. van de Sandt, K. Edlund, M. Lohr, B. Hellwig, R. Marchan, M. Schmidt, J. Rahnenführer, H. Oster, and J. G. Hengstler, “Loss of circadian clock gene expression is associated with tumor progression in breast cancer.,” *Cell cycle (Georgetown, Tex.)*, vol. 13, pp. 3282–91, oct 2014.

15. V. Nygaard, E. A. Rødland, and E. Hovig, “Methods that remove batch effects while retaining group differences may lead to exaggerated confidence in downstream analyses,” *Biostatistics*, vol. 17, no. 1, pp. 29–39, 2016.

16. G. Casella and R. L. Berger, *Statistical inference*, vol. 2. Duxbury Pacific Grove, CA, 2002.

17. F. Takens, “Detecting strange attractors in turbulence,” in *Dynamical systems and turbulence, Warwick 1980*, pp. 366–381, Springer, 1981.

18. T. Sauer, J. A. Yorke, and M. Casdagli, “Embedology,” *Journal of Statistical Physics*, vol. 65, no. 3, pp. 579–616, 1991.

19. J. Huke, “Embedding nonlinear dynamical systems: A guide to takens’ theorem,” 2006.

20. H. Press, A. Teukolsky, T. Vetterling, and P. Flannery, *Numerical Recipes in C++. The Art of Computer Programming*. Cambridge University Press New York, 2002.

21. D. Broomhead, R. Indik, A. Newell, and D. Rand, “Local adaptive galerkin bases for large-dimensional dynamical systems,” *Nonlinearity*, vol. 4, no. 2, p. 159, 1991.

22. J. J. Hughey, T. Hastie, and A. J. Butte, “ZeitZeiger: supervised learning for high-dimensional data from an oscillatory system,” *Nucleic Acids Research*, p. gkw030, 2016.

23. J. Yeung, J. Mermet, C. Jouffe, J. Marquis, A. Charpagne, F. Gachon, and F. Naef, “Transcription factor activity rhythms and tissue-specific chromatin interactions explain circadian gene expression across organs,” *Genome research*, vol. 28, no. 2, pp. 182–191, 2018.

24. J. J. Hughey, T. Hastie, and A. J. Butte, “Zeitzeiger: supervised learning for high-dimensional data from an oscillatory system,” *Nucleic acids research*, vol. 44, no. 8, pp. e80–e80, 2016.

25. C. Vollmers, S. Gill, L. DiTacchio, S. R. Pulivarthy, H. D. Le, and S. Panda, “Time of feeding and the intrinsic circadian clock drive rhythms in hepatic gene expression,” *Proceedings of the National Academy of Sciences*, vol. 106, no. 50, pp. 21453–21458, 2009.

26. E. D. Buhr and J. S. Takahashi, “Molecular components of the mammalian circadian clock,” *Handbook of Experimental Pharmacology*, vol. 217, no. 217, pp. 3–27, 2013.

27. D. Gonze, J. Halloy, and A. Goldbeter, “Robustness of circadian rhythms with respect to molecular noise.,” *Proceedings of the National Academy of Sciences of the United States of America*, vol. 99, no. 2, pp. 673–678, 2002.

28. D. Gonze, J. Halloy, and a. Goldbeter, “Deterministic and stochastic models for circadian rhythms,” *Pathologie-biologie*, vol. 51, no. 4, pp. 227–230, 2003.

29. M. L. Guerriero, O. E. Akman, G. V. Ooijen, and D. Chiarugi, “Stochastic models of cellular circadian rhythms in plants help to understand the impact of noise on robustness and clock structure,” *Frontiers in Plant Science*, vol. 5, no. October, pp. 1–6, 2014.

30. D. B. Forger and C. S. Peskin, “A detailed predictive model of the mammalian circadian clock.,” *Proceedings of the National Academy of Sciences of the United States of America*, vol. 100, no. 25, pp. 14806–11, 2003.

31. A. Relógio, P. O. Westermark, T. Wallach, K. Schellenberg, A. Kramer, and H. Herzel, “Tuning the mammalian circadian clock: robust synergy of two loops.,” *PLoS computational biology*, vol. 7, p. e1002309, dec 2011.

32. G. Minas and D. A. Rand, “Long-time analytic approximation of large stochastic oscillators: Simulation, analysis and inference,” *PLoS Computational Biology*, vol. 13, no. 7, pp. 1–23, 2017.
